# A workflow for practical training in ecological genomics using Oxford Nanopore long-read sequencing

**DOI:** 10.1101/2024.09.03.610948

**Authors:** Robert Foster, Heleen De Weerd, Nathan Medd, Tim Booth, Caitlin Newman, Helen Ritch, Javier Santoyo-Lopez, Urmi Trivedi, Alex D. Twyford

## Abstract

Long-read single molecule sequencing technologies continue to grow in popularity for genome assembly and provide an effective way to resolve large and complex genomic variants. However, uptake of these technologies for teaching and training is hampered by the complexity of high molecular weight DNA extraction protocols, the time required for library preparation and the costs for sequencing, as well as challenges with downstream data analyses. Here, we present a full long-read workflow optimised for teaching, that covers each stage from DNA extraction, to library preparation and sequencing, to data QC and genome assembly and characterisation, that can be completed in under two weeks. We use a specific case study of plant identification, where students identify an anonymous plant sample by sequencing and assembling the genome and comparing it to other samples and to reference databases. In testing, long-read genome skimming of nine wild-collected plant species extracted with a modified kit-based approach produced an average of 8Gb of Oxford Nanopore data, enabling the complete assembly of plastid genomes, and partial assembly of nuclear genomes. In the classroom, all students were able to complete the protocols, and to correctly identify their plant samples based on BOLD searches of barcoding loci extracted from the plastid genome, coupled with phylogenetic analyses of whole plastid genomes. We supply all the learning material and raw data allowing this to be adapted to a range of teaching settings.

## Introduction

Long-read sequencing has become a mainstay of genomics, with benefits including the generation of reads of sufficient length to read through complex repeat regions, and decreasing costs making it increasingly accessible for a diverse range of genomic applications (De Coster et al., 2021; Glinos et al., 2022; Logsdon et al., 2020). Long-read sequencing was hailed as Nature’s Method of the Year 2022 due to developments increasing its utility across genomics (Marx, 2023). Major innovations include ultra-long-reads of over 4Mb with Oxford Nanopore Technologies (ONT) Ultra-Long DNA Sequencing Kit, and ‘near perfect’ highly accurate long-reads with Pacific Biosciences (PacBio) High Fidelity (HiFi) data. Long-read sequencing has underpinned major new research, such as the publication of the first gapless telomere-to-telomere assembly for all 22 human autosomes and the X chromosome (Nurk et al., 2022). These technologies make the prospect of sequencing reference genomes for the diversity of eukaryotic life increasingly possible (Lewin et al., 2022).

While long-read sequencing is now commonly used for research, what is the best approach to bring long-read sequencing into the classroom? There is a huge demand to train the next generation of genome scientists, however most teaching and training still relies on short-read sequencing and typically focuses on ‘easy’ species for sequencing and assembly such as bacteria (Drew & Triplett, 2008; Hotaling et al., 2018). Providing training that is more representative of the challenges faced by researchers working with non-model species is extremely valuable, and where this has been done it has proved successful, such as with the *de novo* assembly and analysis of earwig genomes by students in Chile (Kobayashi et al., 2023). However, such training is far from routine, and often needs to overcome issues at each stage of the workflow. Before sequencing, issues include extracting a suitable quantity and quality of DNA in a classroom setting without dangerous chemicals and laborious protocols, and preparing suitable high-quality libraries. Both good quality DNA and libraries are essential, and have a major impact on the quality of the sequencing data that is generated. In terms of the sequencing, the ONT MinION has been widely used for teaching (Kamathewatta et al., 2019; Watsa et al., 2019; Zaaijer et al., 2016) but the output remains relatively low and expensive per Gb, while the output of many long-read production sequencers, such as the PacBio Revio and the ONT PromethION, while relatively cheap per Gb, have a high unit cost necessitating a degree of multiplexing that can be hard to manage. As such, careful consideration needs to be given to the sequencing strategy. Finally, the management and analysis of large sequence datafiles, and the complexity of downstream analyses (including *de novo* genome assembly and comparative genomic analyses), all pose difficulties.

Here, we focus on developing training in the application of long-read genomics to organismal identification, with a case study in plants. Organismal identification is central to many questions in molecular ecology, from profiling species diversity and abundance in microbial samples with metagenomics, to the authentication of herbal and medicinal products with DNA barcoding (Hollingsworth et al., 2016). This application is ideal for teaching as it is conceptually simple compared to many other areas of molecular ecology (e.g. it does not rely on background knowledge of theoretical population genetics), but has the benefits of involving a representative range of tasks common to other research activities. Our case study uses plants as they are frequently overlooked in laboratory teaching activities in favour of more experimentally tractable groups such as bacteria. This is in part due to their complexity in terms of lab protocols (e.g. frequent need for species-specific protocols to overcome issues such as diverse secondary compounds, Schalamun et al., 2019), and in downstream analyses (i.e. no single barcoding region provides universal species-level resolution, CBOL Plant Working Group, 2009). However, the key role of plants in natural ecosystems, their ease of collection and fewer ethical considerations than many animal groups, and the huge potential of genomic sequencing to improve species identification over DNA barcoding, makes them an excellent case study (Antonelli et al., 2020). In particular, we use common plant species collected at a site in Scotland, allowing us to tie our work into the Darwin Tree of Life Project, which aims to sequence high quality reference genomes for all species in Britain and Ireland (DToL Consortium, 2022).

We teach the principles of species identification focusing on plastid genomes, due to their small size, low repeat content, and abundance in genomic DNA datasets that make them easy to assemble, as well as the extensive availability of plastid data in DNA barcoding reference databases, and the manageable size of plastid genomes for comparative genomic analyses (Twyford & Ness, 2017). We adopt a ‘long-read genome skim’ approach, i.e. generate low coverage long-read data, which proves more affordable than the generation of high coverage data, and still teaches similar principles and would be anticipated to provide similarly high quality plastid genomes. While plastid genome assembly is relatively simple, there are still a number of notable challenges (Turudić et al., 2021). Most plastids are composed of a long single copy (LSC) region, two identical inverted repeats (IRs) and a small single copy (SSC) region. The two identical IRs pose a problem for most standard genome assembly pipelines, which typically assemble circular plastid genomes into three contigs. Secondly, plastids exist as two isomers in the cell, with alternating SSC orientation (Walker et al., 2015). Many genome assemblers report one or other assembly orientation, in a non-standardised fashion, posing an issue for comparative analyses between plastid genomes. These two issues in particular are why various plastid specific genome assembly algorithm have been developed (Dierckxsens et al., 2017; Jin et al., 2020).

Here, we develop an applied practical genomics course aimed at early career researchers, such as first year PhD students. The expectation is that these students will be familiar with some laboratory methods such as PCR, and will have a working knowledge of computing, but have not previously performed genomic sequencing or bioinformatic analyses. The course is intended to fit within a two- week full-time teaching period, such as a short summer school, run with a small cohort of ∼12 students. Our training development aimed to take students through each stage of the genomics workflow, providing a balance of training in wet lab skills and bioinformatics. For the lab work, we chose to use Oxford Nanopore sequencing because: (1) there is an active user community and a range of easy to implement protocols; (2) there is flexibility in sequencing output for small test runs on the Flongle and MinION to production sequencing on the PromethION; (3) ONT has the potential to generate the longest reads out of currently available technologies, making it excellent for teaching long-read sequencing and of high value for many projects such as in *de novo* genome assembly, (4) skills learnt in training can be directly transferred into individual laboratories as many groups own MinIONs. To make our protocol interactive, we gave each student an anonymous (unlabelled) plant sample, with the aim of using the genomic sequencing and bioinformatic analyses taught in the class to provide a correct identification. We first document the development of our training material, including laboratory and bioinformatic methods. We then report the results from training two cohorts of ecological genomics students. Finally, we consider how this teaching provides benefits over other training options, and the potential future utility of long-read sequencing for plant identification.

## Methods

### Wet lab protocol development

To develop a protocol suitable for the classroom, we focused on testing various protocols that could streamline delivery relative to standard long-read workflows, namely: (1) the utility of silica dried plant material, which may compromise read length due to drying that causes fragmentation, but is easy to handle and removes the requirement for liquid nitrogen; (2) kit based extractions, including both the DNeasy Plant Mini Kit, which is popular for Sanger sequencing and Illumina short read sequencing, and the Nucleon PhytoPure kit, which has been used in a range of genomics applications; (3) the use of additional cleanups to compensate for the more rapid extraction protocol that may include coeluting compounds. The performance of these methods was judged based both on DNA and library QC as well as sequencing quality on different ONT platforms.

We sampled commonly cultivated or wild plant species at the King’s Buildings’ Campus of the University of Edinburgh, UK (coordinates 55.924, -3.173). The campus covers 35 hectares and includes a wide range of ornamental trees, shrubs and herbaceous plants, as well as native species in artificial meadows and growing as weeds. Species for sequencing were chosen for their phylogenetic diversity, being diploids with relatively small genome sizes (haploid genome sizes <2Gb), based on available genome size estimates in the Plant DNA C-value database (Table 1 (Pellicer & Leitch, 2020)). Leaf material was collected during the growth season into desiccating silica material. Identical samples were used in testing and in the taught courses, apart from *Antirrhinum majus* which was replaced with *Antirrhinum hispanicum* for teaching.

**Table 1.**
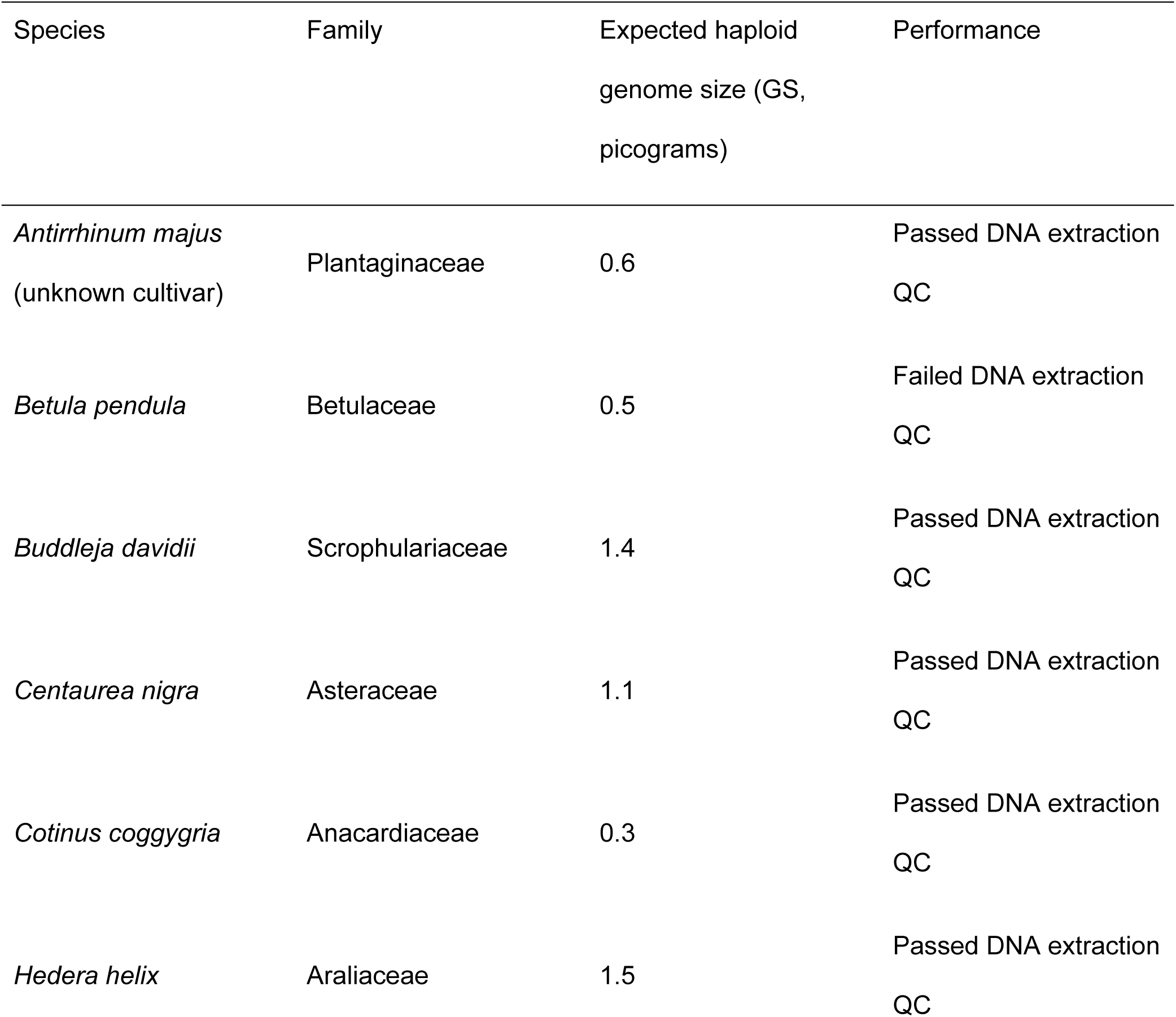
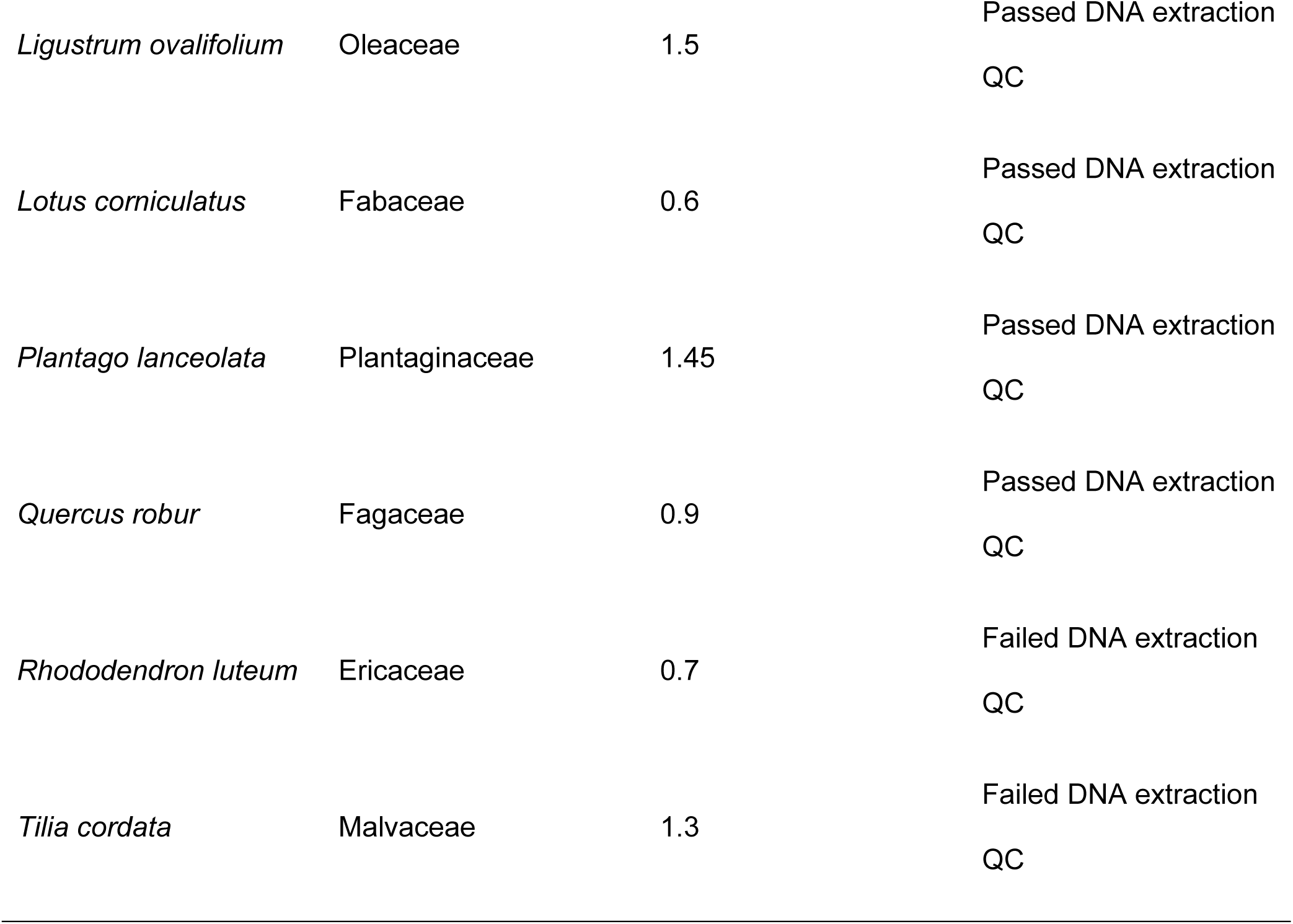
Species used for protocol development. Genome sizes from the Plant DNA C-value database.

Firs, we tested DNA extraction using the Qiagen DNeasy Plant Mini Kit following the manufacturer’s instructions. Dried leaf tissue weighing 10.3 - 27.5 mg was ground to a fine powder using a PowerMasher II device and BioMasher II tubes (Nippi) placed in dry ice. DNA concentrations were estimated using a Qubit HS dsDNA Assay kit, and size distributions checked with a Femto Pulse using a Genomic DNA 165 kb kit (Agilent). A 15-minute incubation at room temperature with a 1:1 volume ratio of AMPure XP beads (Beckman Coulter) was performed to concentrate samples. Two 80% ethanol washes were performed on a magnet, with samples eluted in 48ul of nuclease-free water.

Second, we tested the Nucleon PhytoPure kit (Cytiva), paired with the DNeasy PowerClean Pro Cleanup Kit (Qiagen) to remove polysaccharides and polyphenols which may inhibit sequencing. Dry leaf tissue weighing 13.3 - 16.0 mg was used as an input, extractions and cleanups were performed following manufacturer’s instructions, and samples were eluted in 50ul EB.

Third, we performed small-scale testing of the Qiagen DNeasy Plant Mini Kit as used initially, but additionally with the DNeasy PowerClean Pro Cleanup Kit. This was performed on two test species, following the approaches described above.

The concentration and purity of the extractions was checked using a Qubit HS dsDNA Assay kit and Qubit HS RNA kit, the absorption ratio measurements taken with a NanoDrop ND-1000, and the size distribution checked using a Tapestation 2200 and Genomic DNA ScreenTape and Reagents. Three sampled species that performed poorly were excluded due to low DNA concentrations (Table 1, see Results). Sequencing libraries were prepared from 1ug of concentrated DNA extraction per species (or all the DNA available if <1ug), using the ligation sequencing kit (SQK-LSK109) with native Barcoding Expansion (PCR-free, EXP-NBD104 and EXP-NBD114), following the manufacturer’s protocol. A 700 ng library pool was prepared using a different amount of input DNA for each species, based on their genome sizes (Table S1). Pools were QC’d using the Femto Pulse and Qubit HS dsDNA assay, as above. Pools for trial 1 (multi-species Qiagen DNeasy Plant Mini) and 2 (multi-species Nucleon PhytoPure + cleanup) were prepared and loaded on a PromethION flow cell (version 9.4.1), while the third trial (two-species Qiagen DNeasy Plant Mini + cleanup) was run on a Flongle flowcell in a MinION.

### Bioinformatics protocol development

Overall, our aim with the bioinformatics teaching was to introduce the Linux command shell, then guide students through data QC, nuclear genome and plastid genome assembly, and comparative sequence analysis and phylogenetic analyses. At the end students should have sufficient evidence to identify their species. Teaching material was adapted to provide a background to Linux and the Bash shell. Introductory topics were prepared as an interactive demo, followed by self-led tutorials. The course materials covered filesystem navigation and paths, obtaining on-line help, copying and moving files, gzip compression, viewing and searching in text files, and shell pipelines.

Initial preparation involved installing the NanoPack (De Coster & Rademakers, 2023) software using Bioconda (Grüning et al., 2018) and MambaForge (https://github.com/conda-forge/miniforge). The raw ONT data was basecalled with Guppy version 6.3.8 with default parameters, for use in downstream genome assemblies.

Primary QC of the read data was performed with NanoPlot (version 1.41), part of the NanoPack package, and reports examined, particularly with regard to the length and quality distribution of their reads. The mean quality score reported was used to set the quality trimming cutoff used with chopper (version 0.5.0, also part of NanoPack), and reads under 500bp were also filtered out. Then NanoPlot was re-run on the filtered reads to confirm the results of filtering.

For nuclear genome assemblies, we aimed to illustrate how to run the pipeline and the increasing ease of accessing the nuclear genome, however given the relatively low sequence coverage per sample the assemblies run on our data are expected to be fragmented and incomplete. The trimmed reads were assembled using Redbean (version 2.5, Ruan & Li, 2020). The quality of the assemblies were assessed with QUAST (version 5.2, Gurevich et al., 2013), and corrected using Racon.

Plastid assembly was performed using ptGAUL (version 1.0.5, Zhou et al., 2023). We used a plastid reference database from NCBI refseq version 2.1, with samples chosen to possess the same orientation of the Small Single Copy (SSC), which can be in one of two orientations in plastid genomes, to map the reads as part of the pipeline. The edges created were compared to sequences for the reference library to identify any issues, which were fixed manually. These edges were combined with combine_gfa.py script provided by ptGAUL pipeline and used to generate two paths (i.e. plastid isomers, with the SSC in one of two directions).

PGA (Qu et al., 2019) was used for the annotation of the plastids. A reference plastid of the species or a related species was downloaded for each species from NCBI and used as a reference (Table S4).

Species identity was initially assessed from the sequence data by extracting two widely used plant DNA barcoding regions, *matK* and *rbcL*, based on the plastid genome annotation. These sequences were searched in BOLD (Ratnasingham & Hebert, 2007), with matches in the search table providing the first evidence of the potential identity of the species, and giving high certainty as to the plant family it belongs.

The 9 plant species sequenced belong to 8 different plant families; for each family up to 15 related plastid sequences were downloaded from NCBI (Table S5). These were intended to provide contextual sequence information and a range of genetic distances in phylogenetic analyses, rather than represent comprehensive sampling of the whole family. If the focal species had been sequenced before then the previous assembly was included. Based on the DNA barcoding analysis, above, which was used to infer the family, the user could then select the relevant family alignment and add in their plastid genome assembly using mafft (v7.520, Katoh & Standley, 2013). To test the impact of SSC orientation on alignment quality the option was given to include both paths (e.g. forward and reverse SSC orientation). Tree building was performed with the graphical user interface software ugene for (Okonechnikov et al., 2012) for student cohort 1, and the command line software IQ-Tree 2 (Minh et al., 2020) using automated model selection for student cohort 2. Trees were also constructed using a k-mer based approach with SANS serif v2.3_3A (Rempel & Wittler, 2021), which is suitable for closely related taxa, such as here with a species-level alignment of low-variation plastid sequences. SANS serif provides a quick introduction to k-mers, and also partly overcomes issues with incorrect SSC orientation (i.e. only mismatching k-mers spanning the SSC-IR junction rather than the whole SSC will provide conflicting phylogenetic signal).

### Course delivery and evaluation

We delivered the course to two classes of students (12 or 14 students) in 2023. Students were recruited based on their training needs, and ensuring inclusion from underrepresented groups. Most students were studying for PhDs, though there were some undergraduates, MSc, postdocs and Junior PIs, and students had a range of expertise, from beginners to more experienced users in either the lab or in bioinformatics. The course was taught over six days of a two-week period, with three days in the wet lab to start, followed by a break when data were generated on the PromethION, then three days of bioinformatic analysis. Students were each randomly assigned one silica dried plant sample, without being given its species identity, with some students assigned the same species. DNA extractions were performed with the best performing extraction approach (see Results), followed by library preparation (ONT Kit 109 for cohort 1, updated to Kit 14 for cohort 2). Pools of student libraries were made by Edinburgh Genomics facility staff, before being used in demonstrations of loading a sequencer, using ONT Flongles. Pools were then run on the ONT PromethION to generate sufficient data for assembly. Basecalling was done by facility staff ready for the students to analyse following the bioinformatics methods, above.

For the bioinformatics, each learner on the course was assigned an individual virtual machine (VM) running on the Amazon EC2 commercial cloud service. The VM’s are configured to present a remote XFCE4 desktop via TigerVNC server as well as having some key packages and data pre-installed. For the first day of teaching (introduction to linux and data QC) the t3.medium size (2vCPU, 4GiB RAM) was adequate, but to enable nuclear and plastid genome assembly we modified the VM size to c4.8xlarge (36vCPU, 60GiB RAM) providing sufficient resources for rapid analysis. The cloud hosting service enables the size of the VM’s to be altered at any time, albeit with the proviso that all VM instances are shut down and rebooted.

Full information on both the wet lab and bioinformatics student-led protocols are provided in the Supplementary Materials (Supplementary Protocol 1 and Supplementary Protocol 2).

Each student completed the full lab and bioinformatic exercise. Interpretation of the species identity was initially based on the BOLD database comparison table for *matK* and *rbcL* extracted from the whole plastid genome assembly, and subsequently on the placement of the focal species in the two phylogenetic trees. For the second cohort, data on inferred species ID was gathered after the BOLD searches, and after all analyses were completed. After the classes were complete, we assessed the quality of the plastid genomes for each group (test data, cohort 1 and cohort 2) by aligning the assemblies with previously published assemblies (if available) using mafft, and calling SNPs using snp-dists (https://github.com/tseemann/snp-dists). Based on this we recorded the number of mismatches (SNP variants or indels). Manual curation was required for assemblies where errors were present; in particular we altered sequencing coverage used by ptGAUL, and reorientated scaffolds.

Regions were also reverse complemented to be in the same orientation as other sequences in the alignment.

Once the course was complete, evaluation surveys were sent to participants.

## Results

### Wet lab protocol development

Leaves from the 12 species extracted with the Qiagen DNeasy Plant Mini Kit revealed substantial variation in the quantity and quality of DNA (Table 2). There was a ∼67-fold difference in total yield, from 0.1ug to 6.8ug, from a ∼3-fold difference in tissue input. The average molecular weight of the DNA extracted was generally low (<20kb) from the perspective of starting a long-read library preparation; however, this DNA did not require additional shearing. Levels of impurities inferred from

**Table 2.**
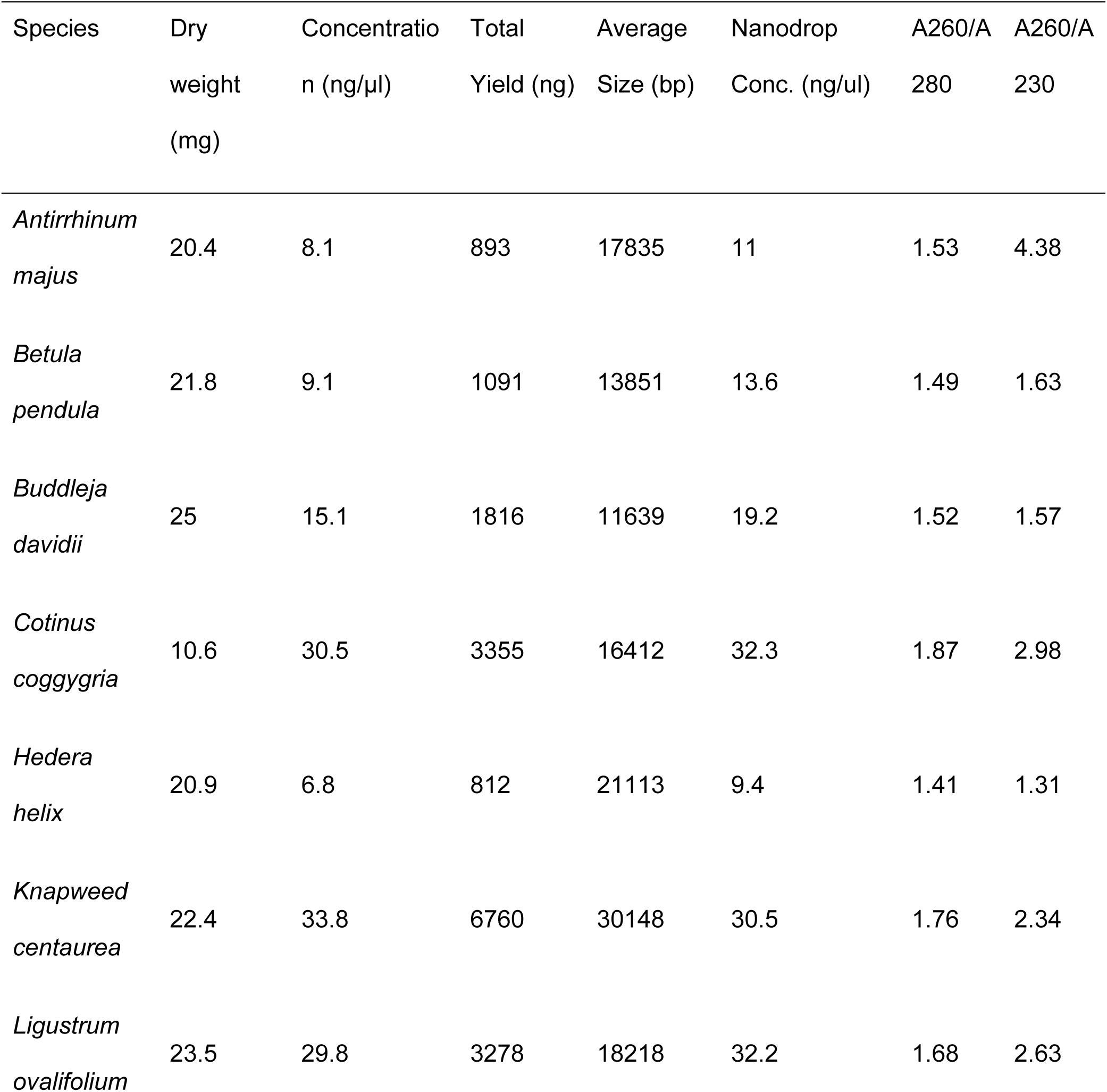
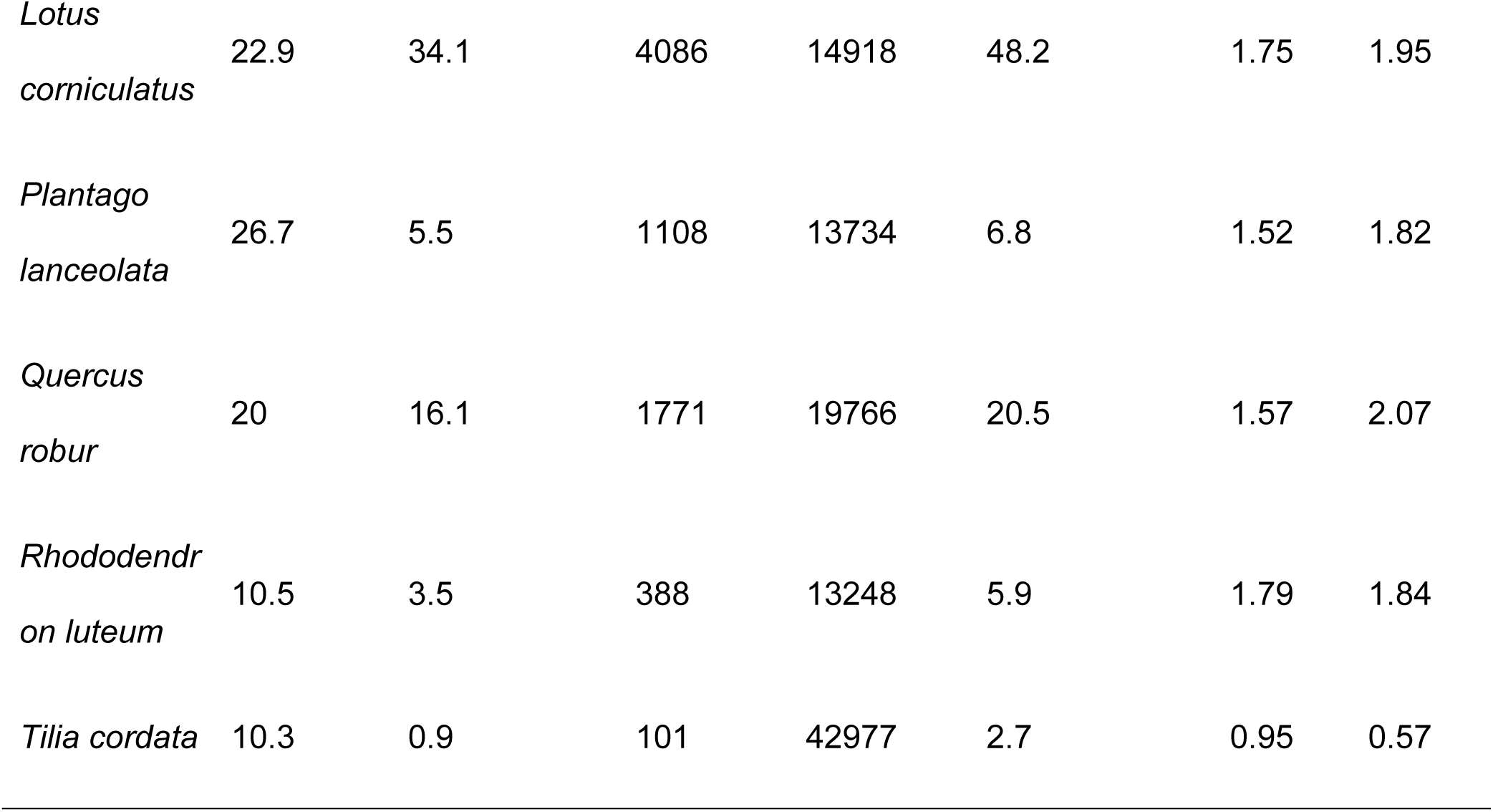
Quality Control of DNA Extractions from the DNeasy Plant Mini Kit.

Nanodrop ratios were highly variable, and ranged from acceptable for *Knapweed centaurea*, to high and likely to be problematic for *Tilia cordata*. Based on these DNA QC results, it was decided to exclude *T. cordata* and continue with library preparation and sequencing for the other species.

Library preparations with the standard ONT protocols at the time (kit v109) and native barcoding expansion was performed for the 11 species, with a 700ng pool weighting samples by their expected genome sizes adapter-ligated and run on a PromethION R9.4.1 flowcell. The results were very poor, with low sequencing yield (<12 Gb). This was due to a rapid accumulation of saturated (blocked) pores (Figure 1a), with fewer than 2,000 active pores after 12 hours, potentially indicating the presence of co-eluting compounds in the DNA extracts and libraries.

**Figure 1.**
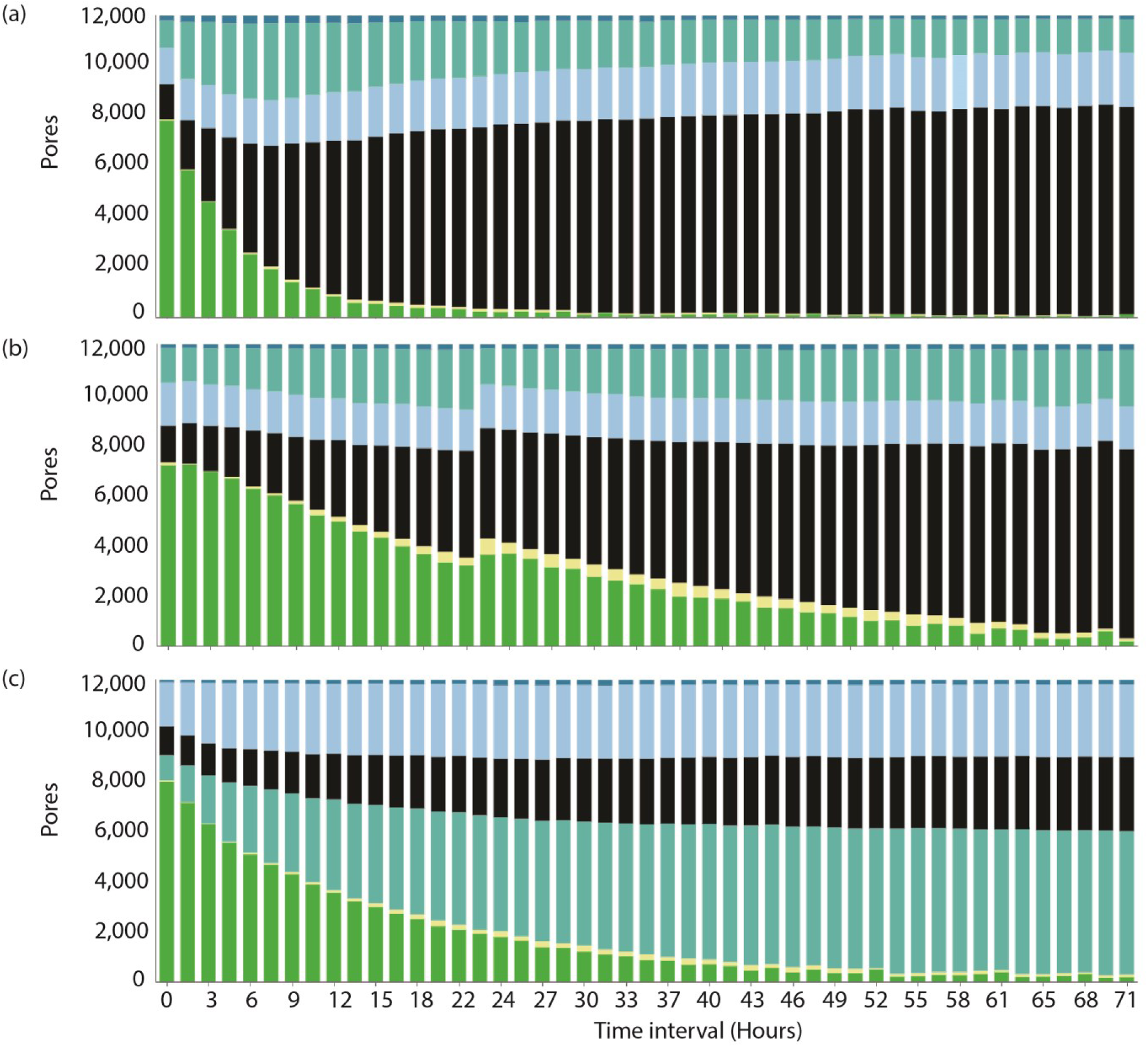
Sequence data generation for (a) Qiagen DNeasy extractions run on the ONT PromethION without cleanup, (b) Nucleon PhytoPure extractions plus Powerclean Pro cleanup, with nuclease wash and library reloaded after 24 hours, (c) Qiagen Dneasy extraction with cleanup, generated by class 1 students. Note how data without cleanup in (a) have fewer active pores. Pores are classified at each time point into: green, pore available for sequencing and black, pores saturated. Additional colours represent: light blue, zero, no current; turquoise, unavailable (may be partially with a nuclease wash and re-loading); dark blue, inactive, no longer suitable for sequencing; yellow, reserved pore, will return to sequencing when required.

Following these issues with extracting DNA using the Qiagen kit alone, we tested an alternative of the Nucleon PhytoPure kit (Cytiva) followed by cleanups with the PowerClean Pro kit (Qiagen). This extraction method generally gave very good results (Table S2), though *Tilia cordata*, *Betula pendula* and *Rhododendron luteum* performed poorly and were excluded from further testing. This library pool was then run on the ONT PromethION, and was optimised for high output for use as test data for bioinformatics protocol development, and thus included a nuclease flush and reloading with fresh library resulting in pore recovery after ∼24 hours (Fig 2b). A total of 82.3Gb of data were generated, with the statistics per species summarized in Table S3. However, this kit requires chloroform and mecaptoethanol, which are poorly suited to a classroom setting, so we explored whether cleanups with the PowerClean Pro kit (Qiagen) alone might rescue Qiagen DNeasy extractions.

**Figure 2.**
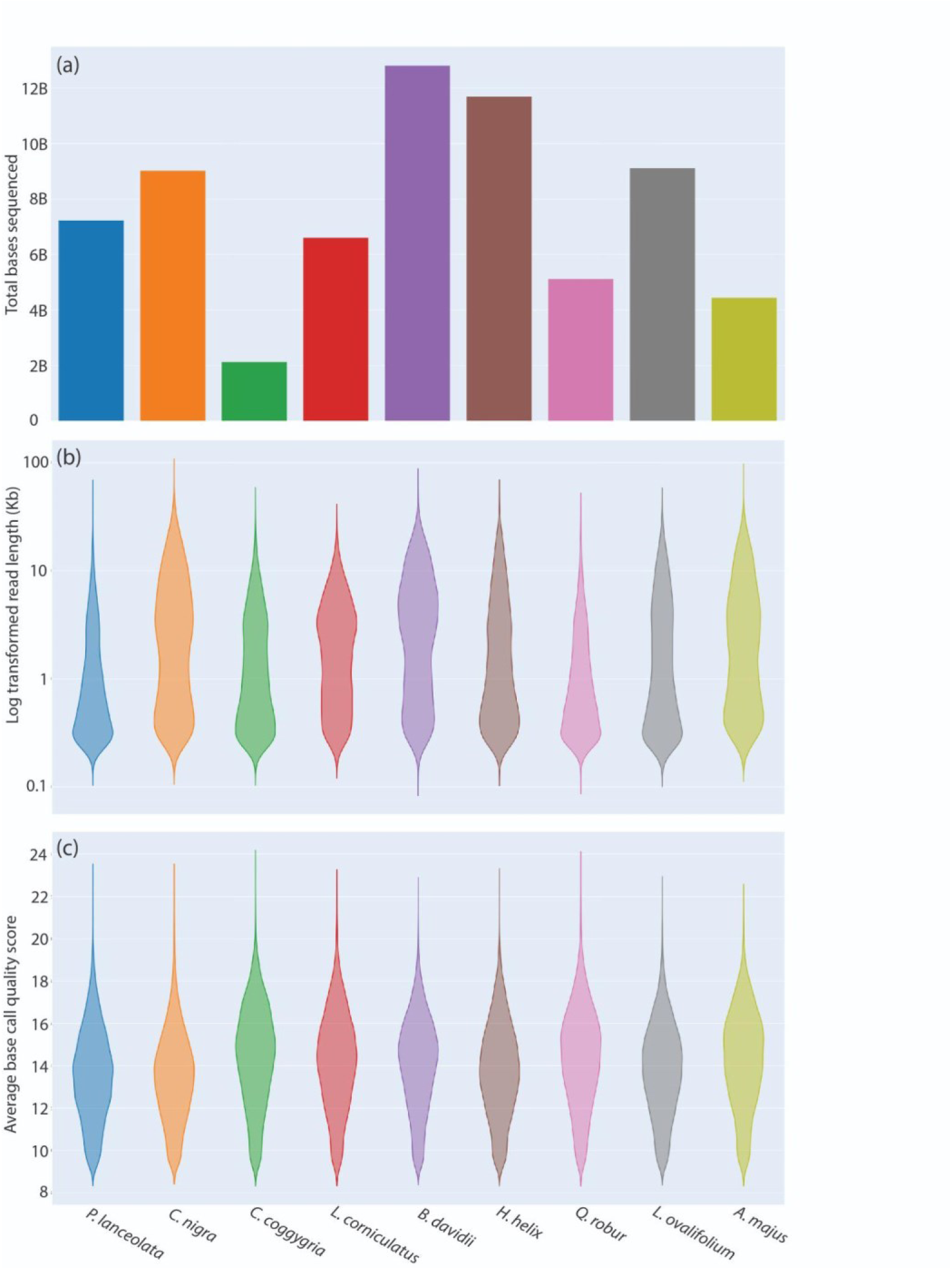
Sequence properties from the Oxford Nanopore sequencing of plant samples in the test data. (a) Total sequencing output in bases, (b) Log transformed read length, (c) Average base quality.

We performed a small-scale test of whether the PowerClean Pro columns may recover DNA from the DNeasy Plant Mini kit, using two of the better performing species from the previous run. The samples had a high purity following the cleanup, with a great improvement in the A260/A280 ratios, indicating these samples should be suitable for nanopore sequencing. Pooled libraries were loaded on a Flongle, which displayed only a small, gradual increase in the number of saturated pores across the run. Therefore, this approach was adopted for the course.

### Wet lab course delivery

Following successful testing, the Qiagen DNeasy Plant Mini kit followed by the PowerClean Pro columns were subsequently used for the two taught classes. Cohort 1 had 11 out of 12 students successfully extract suitable DNA (with the student with a failed extraction supplied a replacement DNA sample), while Cohort 2 had 12 out of 14 students successfully extract suitable DNA (2 supplied with replacement DNA).

All students successfully managed to generate sequence data from their libraries. The total data generated by cohort 1 was 33.1 Gb, with a mean of 2.8Gb for each of 12 samples, while cohort 2 generated 43.0 Gb giving a mean of 3.0Gb for each of 14 samples (representative sequence run output in Figure 1C). This ONT sequence output from student data, which included a cleanup treatment, indicated over double the number of active pores after 9 hours than the initial development test that used a Qiagen extraction without a cleanup. Further data QC was performed as the first stage of bioinformatics delivery (below).

### Bioinformatics protocol development

We used the multi-species Nucleon PhytoPure + cleanup sequencing run as test data for bioinformatics protocol development; this run included between 2.13 and 12.81 Gb of data per sample, with read length averages per sample ranging from 1.55 to 4.64 Kb (read N50 3.6 - 10.3Kb), though there were some outliers (with a longest read of 309 Kb, Figure 2). Base quality was relatively uniform across sequencing libraries, with a mean of 13, Figure 2c).

Nuclear genome assemblies based on the filtered long-read data took less than 30 minutes using RedBean, with the assemblies being characterized by ∼5 - 27,000 scaffolds with a span of 55 - 641 Mb (Table 3). These assemblies represent partial nuclear genomes, spanning on average 32.7% of the size of the nuclear genome based on previously published flow cytometry estimates for the species. However, two assemblies spanned less than 20% of the genome size (*Plantago lanceolata* 3.8%, *Quercus robur* 11.3%), with these taxa the two with the shortest read lengths as well as being at the larger end of the distribution of genome sizes sequenced.

**Table 3.** Nuclear genome assembly statistics for the plant samples sequenced with ONT in the test run.

| Species | Number of contigs | Largest contig | Total length | GC (%) | N50 | L50 |
| --- | --- | --- | --- | --- | --- | --- |
| *A. majus* | 9,624 | 364,924 | 284,004,051 | 34.33 | 47376 | 1771 |
| *B. davidii* | 17,987 | 798,919 | 600,547,147 | 35.01 | 68914 | 2133 |
| *C. coggygia* | 5,464 | 269,542 | 73,214,531 | 34.22 | 17676 | 1284 |
| *C. nigra* | 21,160 | 326,088 | 487,465,112 | 35.53 | 36697 | 3701 |
| *H. helix* | 24,231 | 256,826 | 563,301,186 | 31.25 | 35334 | 4699 |
| *L. corniculatus* | 13,167 | 354,996 | 243,082,846 | 35.58 | 27288 | 2626 |
| *L. ovalifolium* | 26,622 | 810,908 | 640,508,770 | 32.85 | 36177 | 5507 |
| *P. lanceolata* | 7,057 | 138,393 | 54,576,997 | 36.9 | 9528 | 1743 |
| *Q. robur* | 10,545 | 96,505 | 101,572,099 | 33.81 | 12777 | 2517 |

Plastid genomes assembled directly from the long-read data using ptGAUL resulted in complete assemblies for all 9 test samples (Table 4). Despite the relatively low total sequencing coverage, plastid data were highly represented in all libraries, with coverage between 998-fold and 5281-fold, representing 6.8 - 34.3% of reads per dataset. The intermediate assembly files included three contigs corresponding to the LSC, IR, and SSC, with these being effectively stitched together to produce a single circular contig 150-167Kb in length.

**Table 4.**
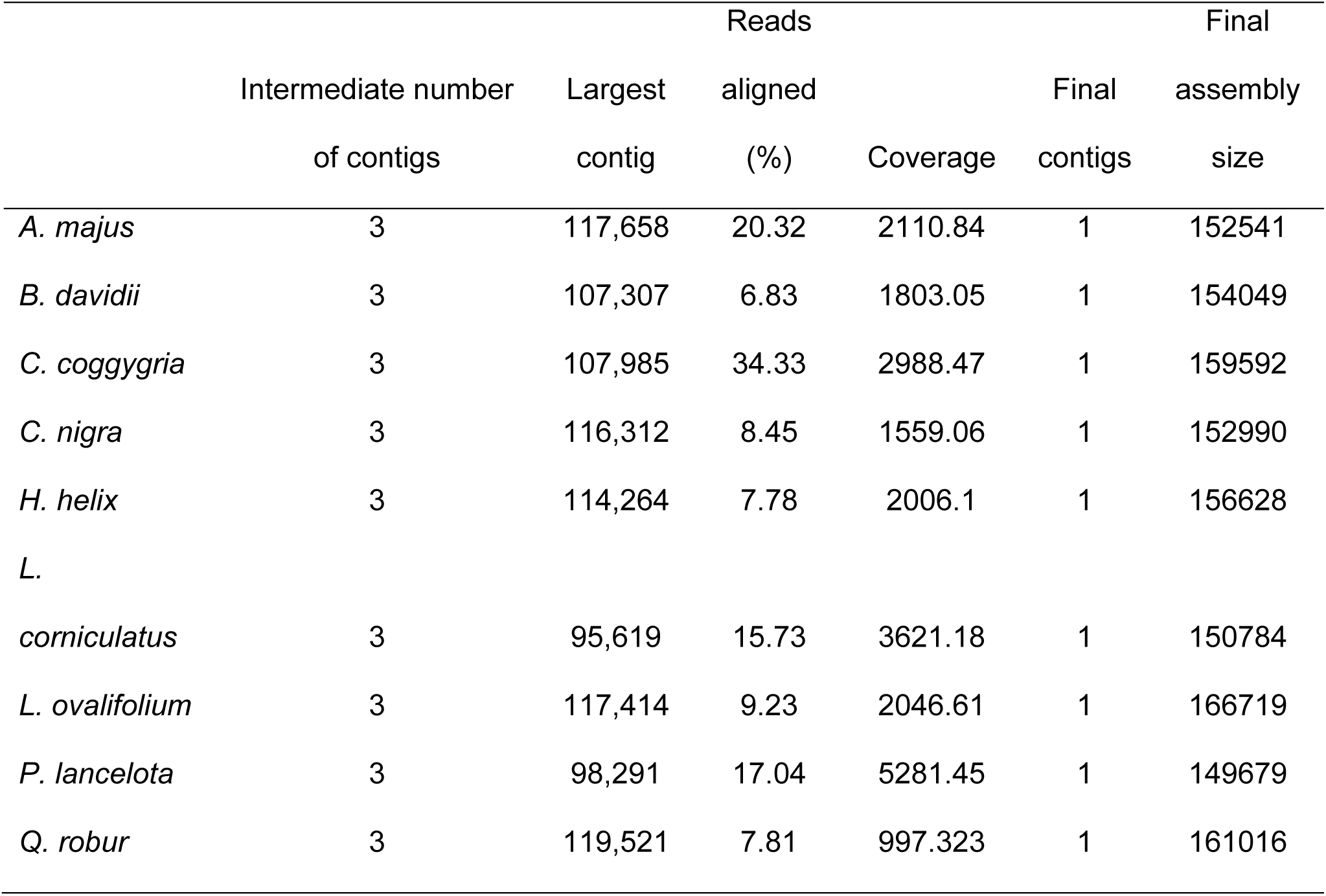
Plastid genome assembly summary statistics for the samples sequenced in the test run. Reads aligned refers to the percentage of reads in the total dataset that map to the finished plastome assembly.

The reference sequence alignments of plastids varied from 158 Kb in Scrophulariaceae, to 175Kb in Fabaceae (Tables S6 and S7). Phylogenetic analyses incorporating the newly generated plastomes were quick to perform, either with ugene, or with IQ-Tree including model finding (Table S8), and generally resulted in the clear placement of the newly generated sequences, often with path 1 and path 2 (alternative SSC arrangement) clustering together, with the alternative arrangement on a long branch. In many cases the newly generated plastid came out sister to the correct species (if included) or a near relative (if the focal species has not previously been sequenced/represented in the reference alignment). K-mer based trees were generally highly consistent with the maximum likelihood phylogeny, with shorter branch lengths between alternative paths (Figure 3).

**Figure 3.**
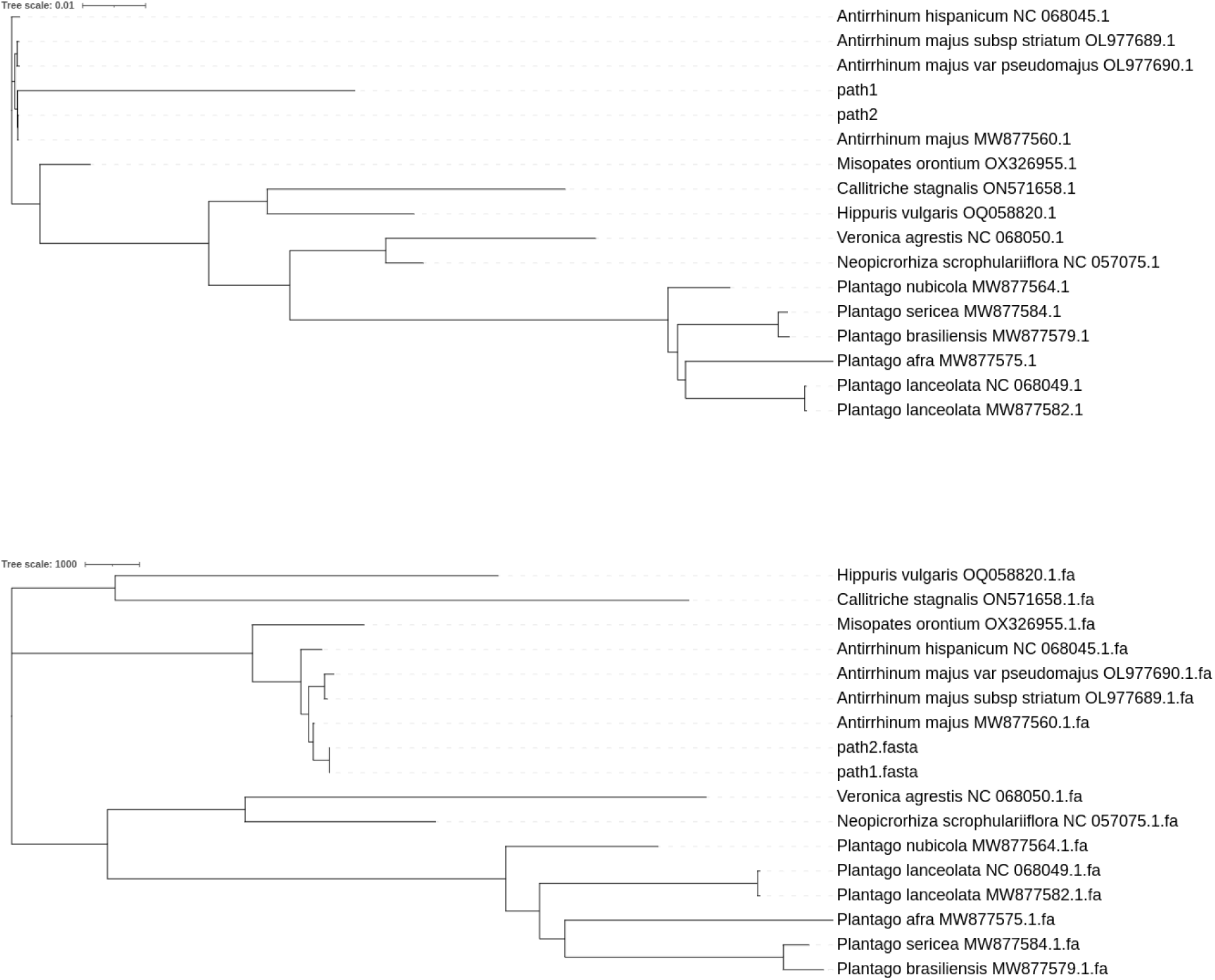
Example phylogenetic trees based on newly assembled plastid genomes, here for the focal taxon *Antirrhinum majus*. Both orientations of the SSC are included (labelled path1 and path2). (a) IQ- Tree analysis using the sequence alignment, (b) SANS serif analysis based on k-mers.

### Bioinformatics course delivery

Cohort 1 produced on average 2.7Gb of ONT data per sample, with average read lengths between 1.4 and 3 Kb, and with an average quality of 13, with the longest read being 785Kb (Table S9). In contrast cohort 2 generated more data per sample but with shorter reads, with on average 3Gb of data per sample, an average read length between 0.7 and 2.8Kb, and an average quality of 12, with the longest read being 292Kb (Table S10). The most notable difference between the test data and the student cohorts were in read length, with a read length N50 across samples in the test data being 6.8Kb, while the student cohorts were 5.0Kb and 3.2Kb for cohort 1 and cohort 2 respectively. Both the test data and student data used the same plant species and the same silica dried material, thus differences are due to the extraction and cleanup approach, or the user.

Nuclear genome assemblies based on the filtered student long-read data were characterized by as few as 653 scaffolds for the smallest genome (∼0.3Gb *C. coggygria*, Cohort 1) to 14,836 for the largest genome (*Ligustrum ovalifolium*, 1.5Gb, Tables S11 and S12). The assemblies partially span the nuclear genome, and are mostly fragmented into thousands of contigs, with some large contigs. While there is considerable variation between samples that were sequenced on multiple occasions, there are some general features, for example four replicate sequencing libraries of the larger genome of *L. ovalifolium* had the longest contig lengths of any samples in the datasets (∼800Kb).

Plastid assemblies from student data were notably variable (Tables S13 & S14), with a number being erroneously large (e.g. all in cohort 1 were >159 Kb and with one outlier of *C. coggygria* being 197Kb; when most land plant plastids are expected to be in the ∼155Kb size range). This resulted in mismatches between student assemblies and the reference data assemblies we generated, above (Table S15 and S16). In cases where student plastid assemblies were characterised by misassembles, typically due to low coverage and short read lengths, these erroneous assemblies still clustered with their nearest relative in k-mer based tre building. In contrast, ML trees led to longer branch lengths making evolutionary relationships and taxon identity harder to interpret. However, further work identified errors are only present in the full assemblies, and not the intermediate sequence scaffolds. Almost all errors were resolved through manual curation of the scaffolds. Once scaffolds were in the correct order and overlaps between sequences correctly identified, the curated assemblies were the correct size. Despite some assembly errors all student plastid genomes had 99.6+% sequence similarity to the reference after curation.

All students successfully annotated the plastid genomes, including the barcoding genes *matK* and *rbcL*. Barcode searches using the BOLD identification engine were more likely to be good matches (returning higher similarity scores, and hitting the correct species) for *matK* than *rbcL*, with the combined evidence from both loci giving reliable family level inference for all samples. In many cases hits were to the correct genus and species, too, though there were some errors and conflicting signal between loci (Figure 4). For cohort 2, where information was collated on the predictions of species identity, 7 students correctly inferred the species based on just the BOLD matches of *matK* and *rbcL*, 5 others identified the correct genus, and 2 others only identified the correct family. After tree building, twelve out of thirteen students who entered their final guess were correct (with one student naming the correct species plus an alternative congeneric taxon).

**Figure 4.**
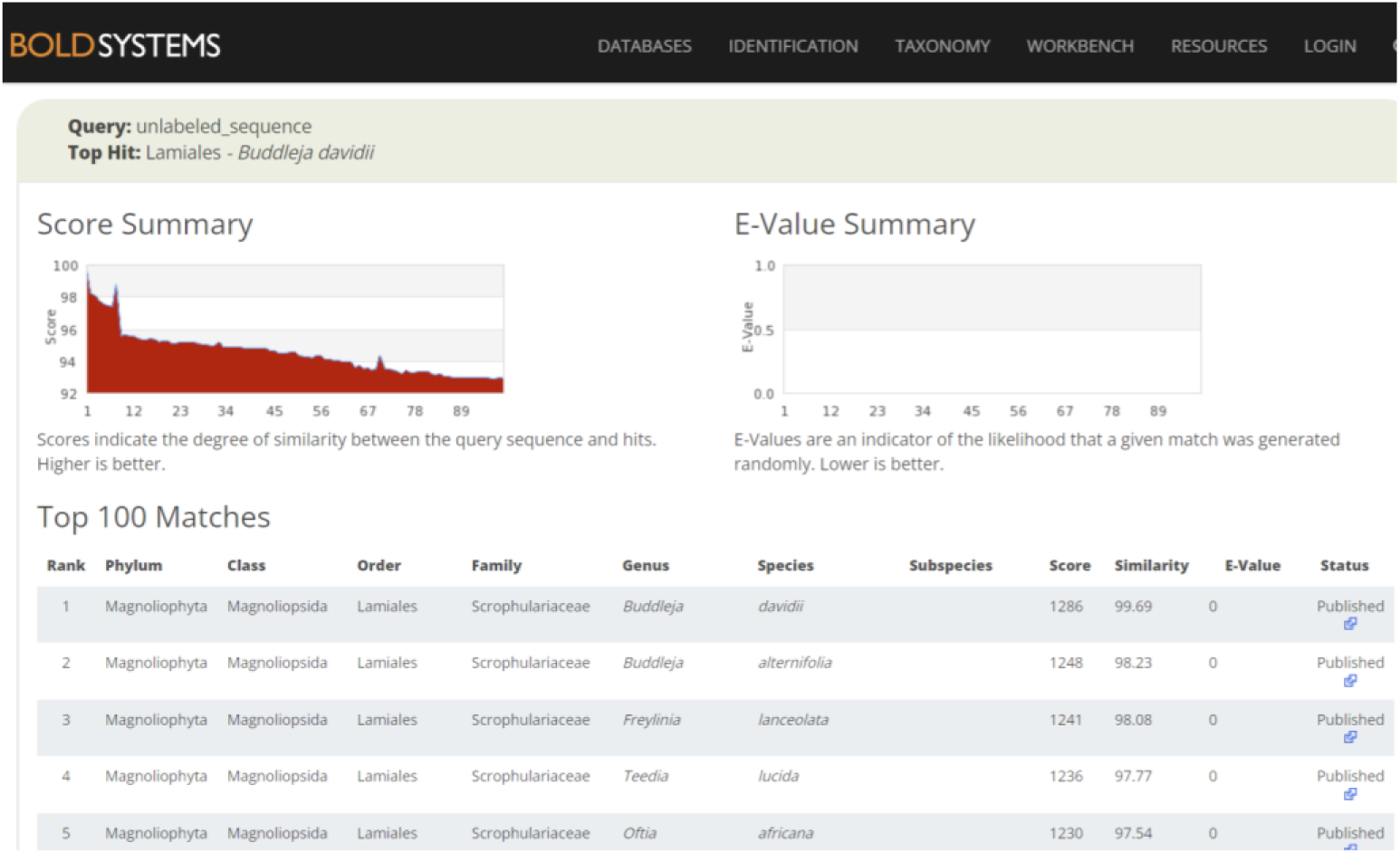
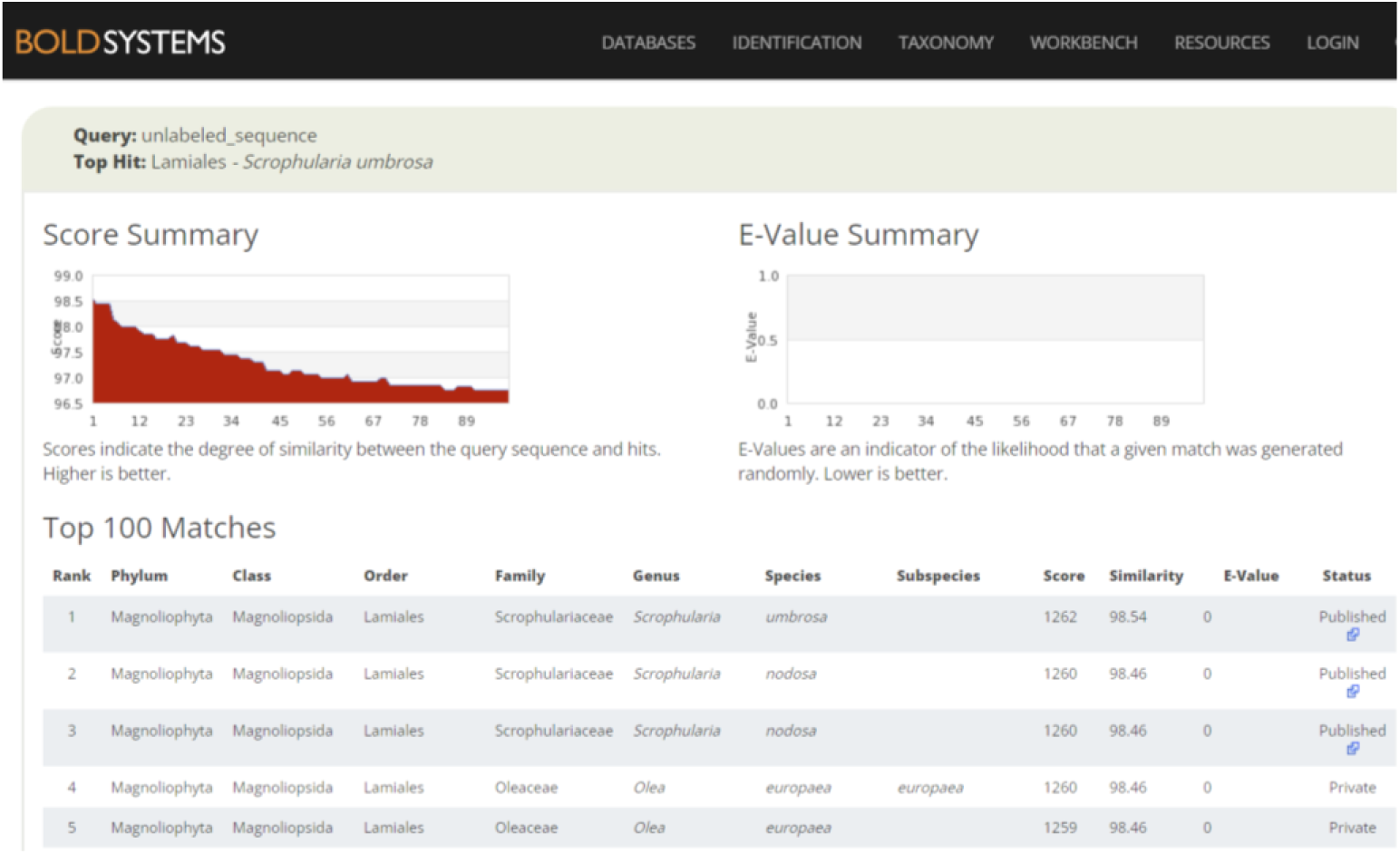
Representative screenshots from BOLD for searches made for the barcoding sequences extracted from the newly assembled *Buddleja davidii* plastome. Top is *matK* and bottom *rbcL*. Note in this case *matK* and *rbcL* have conflicting top hits.

Course questionnaires revealed a high degree of satisfaction, with most students rating the course ‘excellent’, and highlighting different favorite parts (either with the lab, bioinformatics or the whole workflow). An area highlighted for improvement was offering each student to load a MinION to gain individual experience with sequencing.

## Discussion

We have developed a long-read sequencing workflow for plant genomics, and successfully delivered the course to two cohorts of students. By combining multiple readily available laboratory kits, and a range of bioinformatic tools, we have produced a full sample-to-sequence-to-analysis workflow suitable for the classroom. This development has required careful consideration to overcome challenges associated with working with plants, in particular selecting species with small genome sizes, and focusing downstream analyses on plastid genomes that are highly represented in genomic DNA extracts. Below, we consider the wider utility, potential limitations, and areas for future development.

### Development and delivery of a long-read sequencing workflow for the classroom

DNA extraction kits have been widely adopted for first- and second-generation sequencing approaches for their ease of use and reliability in recovering large quantities of clean DNA. However, newer third generation sequencing methods typically require more DNA and a higher quality (Kang et al., 2023). These quality requirements can be hard to assess, for example seemingly suitable high molecular weight DNA may be nicked and this may only become apparent when the library is sequenced, resulting in short reads of limited value (https://www.molecularecologist.com/2018/04/26/dna-extraction-for-pacbio-sequencing/). We have shown that while the widely used Qiagen Plant DNA kit may not be suitable on its own for *de novo* genome sequencing with ONT, due to co-eluting compounds that block the sequencing pores, the DNA can be partially recovered using a cleanup column. While the read lengths are not optimal for applications such as generating chromosomally complete genome assemblies, it proved highly effective in a classroom setting, avoiding the cost, time and use of dangerous chemicals required in many HMW protocols, and resulted in useful genomic data.

While overall our protocol fulfilled our aims, the biggest issue remaining is reliably generating sufficient sequencing output per sample given the relatively large size of flowering plant genomes compared to other species usually used for laboratory training. To minimise costs, multiplexing on a high throughput platform such as the PromethION is necessary. Our approach allowed us to be relatively cost effective, at under £250 per sample, including library preparation and sequencing consumable costs of around £200, and computing costs of around £14. However, ONT is highly variable in its sequencing output, with flow cells producing an order of magnitude of variation. This is hard to predict and depends on sample-specific attributes such as DNA fragment length and the abundance and types of co-eluting secondary compounds. We partly circumvented the need for high coverage by focusing our efforts on analysing the plastid genome, which is naturally enriched in genomic DNA extractions. However even here we encountered issues particularly with the lower coverage student data (see below). Future work should aim to ensure multiplexing that recovers sufficient coverage, either through reducing the number of samples per flowcell, using smaller genome size species, or optimising data output from the sequencing platform (e.g. cleaner DNA blocking fewer pores, size selection to remove small fragments, or wash flow cells and rerun with fresh library samples).

Alternatively, lower coverage may prove suitable if the sequencing quality is higher, for example with improvements with ONT data such as duplex sequencing, or by using PacBio HiFi.

### Application of long-read sequencing in plant identification

Our course focused on the specific application of plant identification, an important field that often relies on low quality degraded material (e.g. old herbarium specimens or mixed environmental samples).

While most research in this area relies on DNA barcoding loci, we found a significant improvement when moving from BOLD searches with *matK* and *rbcL*, to phylogenetic analysis using complete plastid genomes. Here, the larger number of sequence characters coupled with the contextual information provided by tree building helped in the identification of these common plant species. As such, whole plastid genomes may prove particularly useful when nuclear data is hard to recover, and can provide discrimination gains relative to standard barcodes such as in diverse floras and some large genera (Song et al., 2023). While promising in certain settings, plastid genomes alone will prove ineffective for species ID when there is insufficient reference data, and will not resolve identification challenges in more complex groups where plastid genomes do not track species boundaries (e.g. due to hybridization, Wang et al., 2018).

Plastid genomes are widely used in a range of applications such as studies of degraded herbarium specimens, environmental DNA sequencing and broad-scale phylogenetics. Here, genome skimming is usually performed with short reads (Straub et al., 2012), which often produce complete and largely error free plastid assemblies (Twyford & Ness, 2017). In contrast, our long-read assemblies, perhaps counterintuitively, were less accurate and required more curation. This is due to various reasons, including those that are general to the approach as well as those specific to our samples/data. Firstly, a number of attributes of the data generated were suboptimal. We found sequencing coverage to be critical, with our initial test data generating sufficient data to be largely error free following curation, while lower coverage student data produced erroneously large plastid assemblies, which had to be corrected by altering coverage thresholds followed by manual curation. Read length is also important, with the ptGAUL paper reporting reliable assembly results can be achieved with long-reads (>5Kb N50, Zhou et al., 2023), however this read length threshold was only met by two students out of fourteen in our second cohort. Secondly, short-read data typically have few errors (often Q30) while most current ONT data is more error prone (Q10), translating into errors in the assemblies. For the second cohort of students, we used the new ONT v14 chemistry, which should allow duplex reads and therefore give lower errors, however only 2.5% of reads were duplex reads. Thirdly, the bioinformatic pipelines for plastid genome assembly from short-read data are more mature and have developed over the years to account for a wide range of potential issues (such as heteroplasmy, Dierckxsens, Mardulyn, & Smits, 2020), whereas long-read plastid assembly pipelines are still in their infancy.

Despite this, the ptGAUL paper reports most assembly issues are encountered with PCR based library methodologies such as plastid capture or long range PCR rather than whole genome sequencing (Zhou et al., 2023), and long-read specific pipelines for organelle assembly are improving at a rapid pace (Uliano-Silva et al., 2023).

A major benefit of genome skimming is the possibility to not only assemble plastid genomes, but also other regions such as mitochondrial genomes, nuclear ribosomal DNA and high copy nuclear repeats, while questions remain as to the utility of genome skimming for recovering other nuclear genomic data (Straub et al., 2012). Here, we show that 5-11X nuclear genome data provides sufficient coverage to generate a de novo assembly representing approximately a third of the nuclear genome. While these partial genome assemblies may prove problematic for comparative genomic analyses due to mismatched missing data across samples and lack of coverage to correct errors, they may still prove valuable for genome profiling applications such as repeat characterisation, as well as for preliminary marker development. Here again, improvements in sequencing read quality of long-read data will make these types of data increasingly useful for rapid plant genomic characterisation without a reference.

Our work focused on plant identification from a single site, where our plastid genome approach works well and species prove mostly easy to identify. In many cases this represents an easy test case, particularly as the British flora is intensely studied, with a well-worked out taxonomy and an extensive barcode database (Jones et al., 2021). Moreover, the British flora contains ∼1500 flowering plant species from ∼500 genera, therefore there are relatively few congeners in most groups (except groups such as sedges, which were not included in sequencing). For more diverse sites and genera, the approach used here could be adapted from species identification, to species discovery and documentation. This would be particularly useful in underexplored taxa with no existing genomic data.

## Supporting information

Supplementary Protocol 2

Supplementary Protocol 1

## Data accessibility

The full bioinformatics learning materials for the course are available as Jupyter notebooks on the Edinburgh Genomics Github: https://github.com/EdinburghGenomics/NERC_EcologicalGenomics. A static version of the wet laboratory and bioinformatics guides used for teaching are available in the Supporting Information. All data associated with the project are available on ENA in project PRJEB76543.

## Benefit sharing

All resources developed for this course are openly available and teachers are encouraged to adopt any part for their own classes.

All plant samples were collected at the University of Edinburgh, with permission.

## Acknowledgements

We thank all of the participants in the two NERC Advanced Training in Ecological Genomics Courses (April and June 2023) for their involvement, feedback and allowing us to share their data. We also thank Edinburgh Genomics staff for preparing materials and assisting in delivery, in particular Ana Vieira, Matt Arno and Stewart Laing. Thanks to Katerina Guschanski for supporting course developments and ranking applications. We also thank ONT for providing reagents.

## Author contributions

Alex Twyford conceived the study and secured funding. Robert Foster led the development of the wet lab protocols. Heleen De Weerd, Nathan Medd, Tim Booth and Urmi Trivedi developed the bioinfor- matics protocols. All authors contributed to the delivery of the teaching. Alex Twyford drafted the pa- per with input from co-authors.

## Citations

Antonelli, A., Smith, R., Fry, C., Simmonds, M. S., Kersey, P. J., Pritchard, H., … Ainsworth, A. (2020). State of the World’s Plants and Fungi. Royal Botanic Gardens (Kew); Sfumato Foundation.

CBOL Plant Working Group (2009). A DNA barcode for land plants. Proceedings of the National Academy of Sciences, 106(31), 12794–12797.

De Coster, W., & Rademakers, R. (2023). NanoPack2: population-scale evaluation of long-read sequencing data. Bioinformatics, 39(5), btad311. doi:10.1093/bioinformatics/btad311.

De Coster, W., Weissensteiner, M. H., & Sedlazeck, F. J. (2021). Towards population-scale long-read sequencing. Nature Reviews Genetics, 22(9), 572–587. doi:10.1038/s41576-021-00367-3.

Dierckxsens, N., Mardulyn, P., & Smits, G. (2017). NOVOPlasty: de novo assembly of organelle genomes from whole genome data. Nucleic acids research, 45(4), e18–e18.

Dierckxsens, N., Mardulyn, P., & Smits, G. (2020). Unraveling heteroplasmy patterns with NOVOPlasty. NAR Genomics and Bioinformatics, 2(1), lqz011.

Drew, J. C., & Triplett, E. W. (2008). Whole Genome Sequencing in the Undergraduate Classroom: Outcomes and Lessons from a Pilot Course. Journal of Microbiology & Biology Education, 9(1), 3–11. doi:doi:10.1128/jmbe.v9.89

DToL Consortium (2022). Sequence locally, think globally: The Darwin tree of life project. Proceedings of the National Academy of Sciences, 119(4), e2115642118.

Glinos, D. A., Garborcauskas, G., Hoffman, P., Ehsan, N., Jiang, L., Gokden, A., … Garimella, K. (2022). Transcriptome variation in human tissues revealed by long-read sequencing. Nature, 608(7922), 353–359.

Grüning, B., Dale, R., Sjödin, A., Chapman, B. A., Rowe, J., Tomkins-Tinch, C. H., … The Bioconda, T. (2018). Bioconda: sustainable and comprehensive software distribution for the life sciences. Nature Methods, 15(7), 475–476. doi:10.1038/s41592-018-0046-7

Gurevich, A., Saveliev, V., Vyahhi, N., & Tesler, G. (2013). QUAST: quality assessment tool for genome assemblies. Bioinformatics, 29(8), 1072–1075. doi:10.1093/bioinformatics/btt086

Hollingsworth, P. M., Li, D.-Z., van der Bank, M., & Twyford, A. D. (2016). Telling plant species apart with DNA: from barcodes to genomes. Phil. Trans. R. Soc. B, 371(1702), 20150338.

Hotaling, S., Slabach, B. L., & Weisrock, D. W. (2018). Next-generation teaching: a template for bringing genomic and bioinformatic tools into the classroom. Journal of Biological Education, 52(3), 301–313. doi:10.1080/00219266.2017.1357650

Jin, J.-J., Yu, W.-B., Yang, J.-B., Song, Y., DePamphilis, C. W., Yi, T.-S., & Li, D.-Z. (2020). GetOrganelle: a fast and versatile toolkit for accurate de novo assembly of organelle genomes. Genome Biology, 21, 1–31.

Jones, L., Twyford, A. D., Ford, C. R., Rich, T. C., Davies, H., Forrest, L. L., … Hollingsworth, P. M. (2021). Barcode UK: A complete DNA barcoding resource for the flowering plants and conifers of the United Kingdom. Molecular ecology resources, 21(6), 2050–2062.

Kamathewatta, K. I., Bushell, R. N., Young, N. D., Stevenson, M. A., Billman-Jacobe, H., Browning, G. F., & Marenda, M. S. (2019). Exploration of antibiotic resistance risks in a veterinary teaching hospital with Oxford Nanopore long read sequencing. PLOS ONE, 14(5), e0217600. doi:10.1371/journal.pone.0217600

Kang, M., Chanderbali, A., Lee, S., Soltis, D. E., Soltis, P. S., & Kim, S. (2023). High-molecular-weight DNA extraction for long-read sequencing of plant genomes: An optimization of standard methods. Applications in Plant Sciences, 11(3), e11528. 10.1002/aps3.11528

Katoh, K., & Standley, D. M. (2013). MAFFT multiple sequence alignment software version 7: improvements in performance and usability. Molecular Biology and Evolution, 30(4), 772–780.

Kobayashi, S., Maldonado, J. E., Gaete, A., Araya, I., Aguado-Norese, C., Cumplido, N., … School Earwig Genome, C. (2023). DNA sequencing in the classroom: complete genome sequence of two earwig (Dermaptera; Insecta) species. Biological Research, 56(1), 6. doi:10.1186/s40659-023-00414-9

Lewin, H. A., Richards, S., Lieberman Aiden, E., Allende, M. L., Archibald, J. M., Bálint, M., … Bertorelle, G. (2022). The earth BioGenome project 2020: Starting the clock. In (Vol. 119, pp. e2115635118): National Acad Sciences.

Logsdon, G. A., Vollger, M. R., & Eichler, E. E. (2020). Long-read human genome sequencing and its applications. Nature Reviews Genetics, 21(10), 597–614.

Marx, V. (2023). Method of the year: long-read sequencing. Nature Methods, 20(1), 6–11. doi:10.1038/s41592-022-01730-w

Minh, B. Q., Schmidt, H. A., Chernomor, O., Schrempf, D., Woodhams, M. D., Von Haeseler, A., & Lanfear, R. (2020). IQ-TREE 2: new models and efficient methods for phylogenetic inference in the genomic era. Molecular Biology and Evolution, 37(5), 1530–1534.

Nurk, S., Koren, S., Rhie, A., Rautiainen, M., Bzikadze, A. V., Mikheenko, A., … Phillippy, A. M. (2022). The complete sequence of a human genome. Science, 376(6588), 44–53. doi:doi:10.1126/science.abj6987

Okonechnikov, K., Golosova, O., Fursov, M., & Team, U. (2012). Unipro UGENE: a unified bioinformatics toolkit. Bioinformatics, 28(8), 1166–1167.

Pellicer, J., & Leitch, I. J. (2020). The Plant DNA C-values database (release 7.1): an updated online repository of plant genome size data for comparative studies. New phytologist, 226(2), 301–305.

Qu, X.-J., Moore, M. J., Li, D.-Z., & Yi, T.-S. (2019). PGA: a software package for rapid, accurate, and flexible batch annotation of plastomes. Plant Methods, 15(1), 50. doi:10.1186/s13007-019-0435-7

Ratnasingham, S., & Hebert, P. D. (2007). BOLD: The Barcode of Life Data System (http://www.barcodinglife.org). Molecular Ecology Notes, 7(3), 355–364.

Rempel, A., & Wittler, R. (2021). SANS serif: alignment-free, whole-genome-based phylogenetic reconstruction. Bioinformatics, 37(24), 4868–4870.

Ruan, J., & Li, H. (2020). Fast and accurate long-read assembly with wtdbg2. Nature Methods, 17(2), 155–158. doi:10.1038/s41592-019-0669-3

Schalamun, M., Nagar, R., Kainer, D., Beavan, E., Eccles, D., Rathjen, J. P., … Schwessinger, B. (2019). Harnessing the MinION: An example of how to establish long-read sequencing in a laboratory using challenging plant tissue from *Eucalyptus pauciflora*. Molecular ecology resources, 19(1), 77–89. 10.1111/1755-0998.12938

Song, F., Li, T., Yan, H.-F., Feng, Y., Jin, L., Burgess, K. S., & Ge, X.-J. (2023). Plant DNA barcode library for native flowering plants in the arid region of northwestern China. Molecular ecology resources, 23(6), 1389–1402. 10.1111/1755-0998.13797

Straub, S. C., Parks, M., Weitemier, K., Fishbein, M., Cronn, R. C., & Liston, A. (2012). Navigating the tip of the genomic iceberg: Next-generation sequencing for plant systematics. American Journal of Botany, 99(2), 349–364.

Turudić, A., Liber, Z., Grdiša, M., Jakše, J., Varga, F., & Šatović, Z. (2021). Towards the well- tempered chloroplast DNA sequences. Plants, 10(7), 1360.

Twyford, A. D., & Ness, R. W. (2017). Strategies for complete plastid genome sequencing. Molecular ecology resources, 17(5), 858–868.

Uliano-Silva, M., Ferreira, J. G. R. N., Krasheninnikova, K., Blaxter, M., Mieszkowska, N., Hall, N., … Darwin Tree of Life, C. (2023). MitoHiFi: a python pipeline for mitochondrial genome assembly from PacBio high fidelity reads. BMC Bioinformatics, 24(1), 288. doi:10.1186/s12859-023-05385-y

Walker, J. F., Jansen, R. K., Zanis, M. J., & Emery, N. C. (2015). Sources of inversion variation in the small single copy (SSC) region of chloroplast genomes.

Wang, X., Gussarova, G., Ruhsam, M., de Vere, N., Metherell, C., Hollingsworth, P. M., & Twyford, A. D. (2018). DNA barcoding a taxonomically complex hemiparasitic genus reveals deep divergence between ploidy levels but lack of species-level resolution. AoB PLANTS, 10(3), ply026–ply026. doi:10.1093/aobpla/ply026

Watsa, M., Erkenswick, G. A., Pomerantz, A., & Prost, S. (2019). Genomics in the jungle: using portable sequencing as a teaching tool in field courses. bioRxiv, 581728. doi:10.1101/581728

Zaaijer, S., Columbia University Ubiquitous Genomics, c., & Erlich, Y. (2016). Using mobile sequencers in an academic classroom. eLife, 5, e14258. doi:10.7554/eLife.14258

Zhou, W., Armijos, C. E., Lee, C., Lu, R., Wang, J., Ruhlman, T. A., … Jones, C. D. (2023). Plastid Genome Assembly Using Long-read data. Molecular ecology resources, 23(6), 1442–1457. 10.1111/1755-0998.13787

