## Supplementary material for "A workflow for practical training in ecological genomics using Oxford Nanopore long-read sequencing": Table S1

**Table S1.** Pooling of samples from the Qiagen DNeasy kit extraction for the first run of the ONT PromethION. Genome sizes from the Plant DNA C-value database.

| Species | Genome size (1C, picograms) | Proportion of pool | Proportion of 700ng |
|---|---|---|---|
| *Antirrhinum majus* | 0.6 | 0.057 | 39.8 |
| *Betula pendula* | 0.5 | 0.047 | 33.2 |
| *Buddleja davidii* | 1.4 | 0.133 | 92.9 |
| *Centaurea nigra* | 1.1 | 0.104 | 73 |
| *Cotinus coggygria* | 0.3 | 0.028 | 19.9 |
| *Hedera helix* | 1.5 | 0.142 | 99.5 |
| *Ligustrum ovalifolium* | 1.5 | 0.142 | 99.5 |
| *Lotus corniculatus* | 0.6 | 0.057 | 39.8 |
| *Plantago lanceolata* | 1.45 | 0.137 | 96.2 |
| *Quercus robur* | 0.9 | 0.085 | 59.7 |
| *Rhododendron luteum* | 0.7 | 0.066 | 46.4 |
| *Tilia cordata* | | - | - |
| **Sum** | 10.5 | 1 | 700 |

**Table S2.** Quality Control of purified DNA extractions from the Nucleon PhytoPure kit.

| Species | Dry weight (mg) | Concentration (ng/µl) | Total Yield (ng) | Peak Size (bp) | Nanodrop Conc. (ng/µl) | A260/ A280 | A260/ A230 |
|---|---|---|---|---|---|---|---|
| *Antirrhinum majus* | 15.4 | 26.8 | 1340 | 26080 | 22.4 | 2.09 | 2.41 |
| *Betula pendula* | 15.7 | 1.5 | 77 | 16404 | 0.08 | 6.15 | 1.37 |
| *Buddleja davidii* | 13.5 | 49.2 | 2460 | 22702 | 43.2 | 1.9 | 2.09 |
| *Centaurea nigra* | 14.5 | 58.6 | 2930 | 26109 | 57.4 | 1.92 | 2.16 |
| *Cotinus coggygria* | 14 | 5.8 | 288 | 13139 | 7.4 | 1.79 | 1.23 |
| *Hedera helix* | 16 | 24 | 1200 | 21670 | 19.1 | 1.91 | 2.15 |
| *Ligustrum ovalifolium* | 14.6 | 6 | 302 | 23427 | 4.4 | 4.41 | 1.66 |
| *Lotus corniculatus* | 13.3 | 21.6 | 1080 | 22449 | 22.1 | 2.09 | 1.91 |

| | | | | | | | |
|---|---|---|---|---|---|---|---|
| *Plantago lanceolata* | 13.8 | 7.1 | 356 | 14977 | 8.1 | 2.96 | 1.26 |
| *Quercus robur* | 14.3 | 11.5 | 575 | 19356 | 9.6 | 2.2 | 1.53 |
| *Rhododendron luteum* | 15.2 | 2.9 | 147 | 56895 | 5.2 | 1.76 | 0.68 |
| *Tilia cordata* | 14.9 | 0.8 | 40 | 19159 | 1.3 | 4.15 | 0.48 |

**Table S3.** Data summary for the ONT PromethION run from the samples extracted using the Nucelon Phytopure Kit, which was used as test data for bioinformatics development.

| Species | #Reads | Yield (Gb) | Max Length | N50 | Mean length |
|---|---|---|---|---|---|
| *P. lanceolata* | 4,680,121 | 7.23 | 220,062 | 3,558 | 1,545.40 |
| *Q. robur* | 3,173,351 | 5.12 | 286,859 | 3,644 | 1,614.10 |
| *C. coggygria* | 994,193 | 2.12 | 71,199 | 4,590 | 2,138.50 |
| *L. corniculatus* | 2,365,111 | 6.6 | 309,986 | 5,009 | 2,794.70 |
| *L. ovalifolium* | 3,058,239 | 9.1 | 173,581 | 7,456 | 2,981.30 |
| *H. helix* | 3,828,755 | 11.69 | 177,817 | 7,813 | 3,054.60 |
| *A. majus* | 1,113,168 | 4.4 | 132,834 | 9,130 | 3,990.60 |
| *B. davidii* | 2,761,247 | 12.81 | 277,294 | 9,656 | 4,640.60 |
| *C. nigra* | 2,014,654 | 9.02 | 182,675 | 10,308 | 4,477.50 |

**Table S4.** Accessions for references used for plastid genome annotation using PGA.

| Species | NCBI accession ID |
|---|---|
| *Antirrhinum majus* | OL977689.1 |
| *Buddleja davidii* | NC069921.1 |
| *Centaurea cyanus* | NC066898.1 |
| *Cotinus coggygria* | NC054342.1 |
| *Hedera helix* | OK539585.1 |
| *Ligustrum ovalifolium* | MH559272.1 |
| *Lotus corniculatus* | MT528596.1 |
| *Plantago lanceolata* | MW877582.1 |
| *Quercus robur* | OW028778.1 |

**Table S5.** Family and species used to generate reference alignments for phylogenetic analyses of plastid genomes.

| **Family** | **Species** | **GenbankID** |
|---|---|---|
| Anacardiaceae | *Cotinus coggygria* | NC_054342.1 |
| Anacardiaceae | *Cotinus coggygria* var. *coggygria* | ON167906.1 |
| Anacardiaceae | *Cotinus coggygria* var. *glaucophyllus* | ON167903.1 |
| Anacardiaceae | *Cotinus coggygria* var. *pubescens* | ON167905.1 |
| Anacardiaceae | *Cotinus nanus* | NC_067635.1 |
| Anacardiaceae | *Cotinus szechuanensis* | NC_067637.1 |
| Anacardiaceae | *Buchanania latifolia* | OM000214.1 |
| Anacardiaceae | *Rhus potaninii* | MT230556.1 |
| Anacardiaceae | *Choerospondias axillaris* | OL405119.1 |
| Araliaceae | *Hedera helix* | OK539585.1 |
| Araliaceae | *Hedera nepalensis var sinensis* | MK130890.1 |
| Araliaceae | *Hedera rhombea* | MT991757.1 |
| Araliaceae | *Aralia caesia* | ON493677.1 |
| Araliaceae | *Brassaiopsis hainla* | KC456164.1 |
| Araliaceae | *Cephalaralia cephalobotrys* | MW183403.1 |
| Araliaceae | *Dendropanax morbifer* | NC_027607.1 |
| Araliaceae | *Schefflera heptaphylla* | NC_029764.1 |
| Araliaceae | *Panax ginseng* | AY582139.1 |
| Asteraceae | *Centaurea diffusa* | NC_024286.1 |
| Asteraceae | *Bidens biternata* | MW551951.1 |
| Asteraceae | *Dahlia pinnata* | OP006582.1 |
| Asteraceae | *Elephantopus scaber* | NC_061907.1 |
| Asteraceae | *Inula linariifolia* | NC_063571.1 |
| Asteraceae | *Lactuca perennis* | NC_066752.1 |

| | | |
|---|---|---|
| Asteraceae | *Neopallasia pectinata* | NC_053639.1 |
| Asteraceae | *Saussurea leontodontoides* | MK953477.1 |
| Asteraceae | *Adenostemma madurense* | OP598193.1 |
| Fabaceae | *Anthyllis barba-jovis* | ON009079.1 |
| Fabaceae | *Coronilla valentina* | ON009080.1 |
| Fabaceae | *Lotus japonicus* | AP002983.1 |
| Fabaceae | *Securigera varia* | MW125582.1 |
| Fabaceae | *Indigofera stachyodes* | MZ768851.1 |
| Fabaceae | *Cullen corylifolium* | NC_042700.1 |
| Fabaceae | *Cajanus cajan* | KX672004.1 |
| Fagaceae | *Quercus robur* | MG678035.1 |
| Fagaceae | *Quercus yunnanensis* | MW829658.1 |
| Fagaceae | *Quercus fabri* | NC_061594.1 |
| Fagaceae | *Castanea crenata* | MN402457.1 |
| Fagaceae | *Castanopsis fabri* | NC_072671.1 |
| Fagaceae | *Fagus crenata* | NC_041252.1 |
| Fagaceae | *Lithocarpus glaber* | MZ750954.1 |
| Fagaceae | *Lithocarpus konishii* | ON422319.1 |
| Fagaceae | *Quercus gambleana* | NC_069210.1 |
| Oleaceae | *Ligustrum lucidum* | MH394207.1 |
| Oleaceae | *Abeliophyllum distichum* | KT274029.1 |
| Oleaceae | *Chengiodendron matsumuranum* | OP545809.1 |
| Oleaceae | *Forsythia viridissima* | MW856917.1 |
| Oleaceae | *Fraxinus chinensis* | MW599993.1 |
| Oleaceae | *Myxopyrum hainanense* | MN908148.1 |
| Oleaceae | *Olea brachiata* | MT560008.1 |
| Oleaceae | *Syringa oblata* | MT025818.1 |

| | | |
|---|---|---|
| Oleaceae | *Nestegis lanceolata* | MH817917.1 |
| Plantaginaceae | *Antirrhinum hispanicum* | NC_068045.1 |
| Plantaginaceae | *Antirrhinum majus* | MW877560.1 |
| Plantaginaceae | *Antirrhinum majus* subsp. *striatum* | OL977689.1 |
| Plantaginaceae | *Antirrhinum majus* var. *pseudomajus* | OL977690.1 |
| Plantaginaceae | *Misopates orontium* | OX326955.1 |
| Plantaginaceae | *Hippuris vulgaris* | OQ058820.1 |
| Plantaginaceae | *Callitriche stagnalis* | ON571658.1 |
| Plantaginaceae | *Neopicrorhiza scrophulariiflora* | NC_057075.1 |
| Plantaginaceae | *Veronica agrestis* | NC_068050.1 |
| Plantaginaceae | *Plantago afra* | MW877575.1 |
| Plantaginaceae | *Plantago brasiliensis* | MW877579.1 |
| Plantaginaceae | *Plantago lanceolata* | MW877582.1 |
| Plantaginaceae | *Plantago lanceolata* | NC_068049.1 |
| Plantaginaceae | *Plantago nubicola* | MW877564.1 |
| Plantaginaceae | *Plantago sericea* | MW877584.1 |
| Scrophulariaceae | *Buddleja alternifolia* | MN395662.1 |
| Scrophulariaceae | *Buddleja davidii* | NC_069921.1 |
| Scrophulariaceae | *Buddleja forrestii* | OP007421.1 |
| Scrophulariaceae | *Buddleja japonica* | OP007355.1 |
| Scrophulariaceae | *Buddleja nivea* | OP007398.1 |
| Scrophulariaceae | *Buddleja yunnanensis* | OP007364.1 |
| Scrophulariaceae | *Leucophyllum frutescens* | MN044638.1 |
| Scrophulariaceae | *Bontia daphnoides* | MN044637.1 |
| Scrophulariaceae | *Eremophila pilosa* | NC_068056.1 |

**Table S6.** Sequence alignment lengths per family based on the reference plastomes.

| Family | Alignment length |
| --- | --- |
| Anacardiaceae | 167,994 |
| Araliaceae | 159,169 |
| Asteraceae | 160,816 |
| Fabaceae | 175,536 |
| Fagaceae | 164,833 |
| Oleaceae | 159,953 |
| Plantaginaceae | 173,349 |
| Scrophulariaceae | 158,283 |

**Table S7.** Sequence alignment lengths per family based on the reference plastomes and the newly assembled plastid genomes. Assemblies based on the ONT test data, and include the two paths (alternate SSC orientations).

| Species (Family) | Alignment length |
| --- | --- |
| *A. majus* (Plantaginaceae) | 176,344 |
| *B. davidii* (Scrophulariaceae) | 161,177 |
| *C. coggygria* (Anacardiaceae) | 170,715 |
| *C. nigra* (Asteraceae) | 164,073 |
| *H. helix* (Araliaceae) | 162,080 |
| *L. corniculatus* (Fabaceae) | 178,981 |
| *L. ovalifolium* (Oleaceae) | 174,538 |
| *P. lanceolota* (Plantaginaceae) | 176,134 |
| *Q. robur* (Fagaceae) | 168,113 |

**Table S8.** Best-fitting evolutionary models per alignment, calculated with ModelFinder implemented in IQ-Tree.

| Focal species (Family) | Best-fitting model |
| --- | --- |
| *A. majus* (Plantaginaceae) | TVM+F+R2 |

| | |
|---|---|
| *B. davidii* (Scrophulariaceae) | TVM+F+R3 |
| *C. coggygria* (Anacardiaceae) | TVM+F+R2 |
| *C. nigra* (Asteraceae) | TVM+F+R2 |
| *H. helix* (Araliaceae) | TVM+F+I+G4 |
| *L. corniculatus* (Fabaceae) | TVM+F+R2 |
| *L. ovalifolium* (Oleaceae) | TVM+F+R2 |
| *P. lanceolota* (Plantaginaceae) | GTR+F+R4 |
| *Q. robur* (Fagaceae) | TVM+F+R2 |

**Table S9.** Summary of raw data generated by student Cohort 1 in April 2023 on the ONT PromethION.

| Participant ID | Species | Mean read length | Mean read quality | #Reads | N50 read length | Total bases |
|---|---|---|---|---|---|---|
| A1 | *Q. robur* | 1,703.80 | 13.4 | 1,694,131 | 3,867 | 2,886,401,183 |
| A2 | *L. ovalifolium* | 2,237.70 | 13 | 1,155,558 | 5,246 | 2,585,771,793 |
| A3 | *H. helix* | 1,351.60 | 12.9 | 1,741,375 | 3,661 | 2,353,628,367 |
| A4 | *A. majus* | 1,882.50 | 13.4 | 1,339,905 | 3,603 | 2,522,410,261 |
| A5 | *L. ovalifolium* | 1,439.60 | 12.9 | 2,700,496 | 3,668 | 3,887,645,283 |
| A6 | *B. davidii* | 1,934.50 | 13.1 | 3,358,152 | 4,551 | 6,496,368,956 |
| A7 | *H. helix* | 1,793.70 | 12.9 | 1,699,316 | 5,363 | 3,048,090,732 |
| A8 | *B. pendula* | 2,549.80 | 13.1 | 455,919 | 5,282 | 1,162,488,229 |
| A9 | *C. coggygria* | 2,240.90 | 13.4 | 263,460 | 4,835 | 590,377,383 |
| A10 | *P. lanceolata* | 3,058.90 | 12 | 22* | 6,279 | 67,295 |
| A11 | *L. ovalifolium* | 2,070.10 | 13 | 2,195,738 | 6,060 | 4,545,397,527 |
| A12 | *C. nigra* | 2,838.10 | 12.7 | 1,055,760 | 6,851 | 2,996,395,619 |

**Table S10.** Summary of raw data generated by student Cohort 2 in June 2023 on the ONT PromethION.

| Participant ID | Species | Mean read length | Mean read quality | #Reads | N50 read length | Total bases |
|---|---|---|---|---|---|---|
| J1 | *P. lanceolata* | 850.5 | 11.5 | 2,693,750 | 1,141 | 2,291,033,347 |
| J2 | *P. lanceolata* | 670.9 | 11.3 | 3,646,997 | 764 | 2,446,597,357 |

| | | | | | | |
|---|---|---|---|---|---|---|
| J3 | *C. nigra* | 1,066.00 | 11.7 | 2,329,335 | 1,390 | 2,483,033,672 |
| J4 | *C. nigra* | 1,970.30 | 11.5 | 1,851,677 | 4,453 | 3,648,449,670 |
| J5 | *L. corniculatus* | 1,457.10 | 12.4 | 1,568,598 | 2,710 | 2,285,538,038 |
| J6 | *B. davidii* | 945.5 | 12.2 | 4,370,878 | 1,164 | 4,132,846,430 |
| J7 | *C. coggygria* | 1,887.00 | 12.6 | 638,846 | 3,524 | 1,205,521,873 |
| J8 | *L. ovalifolium* | 2,805.80 | 12 | 617,932 | 5,721 | 1,733,767,955 |
| J9 | *H. helix* | 1,454.80 | 11.8 | 2,074,724 | 2,495 | 3,018,340,661 |
| J10 | *H. helix* | 1,952.70 | 11.8 | 2,283,763 | 4,166 | 4,459,596,287 |
| J11 | *L. ovalifolium* | 2,370.00 | 12.2 | 2,232,044 | 5,683 | 5,289,984,176 |
| J12 | *Q. robur* | 1,979.10 | 12.5 | 1,200,139 | 3,974 | 2,375,204,334 |
| J13 | *A. hispanicum* | 1,859.10 | 12.5 | 3,244,392 | 3,407 | 6,031,556,463 |
| J14 | *L. ovalifolium* | 2,260.90 | 12 | 681,466 | 4,870 | 1,540,738,952 |

**Table S11.** Summary statistics for nuclear genome assemblies generated by students from Cohort 1 using Redbean.

| Participant ID | Species | # contigs | Largest contig | Total length | N50 | L50 |
|---|---|---|---|---|---|---|
| A1 | *Q. robur* | 3912 | 184160 | 36771063 | 10984 | 1037 |
| A2 | *L. ovalifolium* | 2765 | 582238 | 24381336 | 9796 | 739 |
| A3 | *H. helix* | 1383 | 397103 | 11860239 | 9616 | 369 |
| A4 | *A. majus* | 2811 | 269397 | 23401188 | 9158 | 800 |
| A5 | *L. ovalifolium* | 3676 | 811708 | 29581985 | 9074 | 979 |
| A6 | *B. davidii* | 13025 | 391076 | 192877901 | 19646 | 3047 |
| A7 | *H. helix* | 3363 | 213588 | 32720855 | 11394 | 908 |
| A8 | *B. pendula* | 1154 | 328679 | 10185376 | 9417 | 319 |
| A9 | *C. coggygria* | 653 | 277069 | 5927586 | 10124 | 178 |
| A10 | *P. lanceolata* | 961 | 173415 | 9036282 | 11078 | 216 |
| A11 | *L. ovalifolium* | 9702 | 806831 | 107249807 | 13257 | 2554 |
| A12 | *C. nigra* | 3417 | 176330 | 33665608 | 11523 | 918 |

**Table S12.** Summary statistics for nuclear genome assemblies generated by students from Cohort 2 using Redbean.

| Participant ID | Species | # contigs | Largest contig | Total length | N50 | L75 |
|---|---|---|---|---|---|---|
| J1 | *P. lanceolata* | 245 | 128067 | 2015632 | 9950 | 115 |
| J2 | *P. lanceolata* | 157 | 80436 | 986997 | 8219 | 73 |
| J3 | *C. nigra* | 1214 | 127240 | 6910266 | 6974 | 618 |
| J4 | *C. nigra* | 5604 | 181804 | 45366461 | 9337 | 3065 |

| | | | | | | |
|---|---|---|---|---|---|---|
| J5 | *L. corniculatus* | 2511 | 351488 | 17941983 | 7916 | 1389 |
| J6 | *B. davidii* | 976 | 132554 | 4285639 | 5271 | 485 |
| J7 | *C. coggygria* | 1811 | 271916 | 13822814 | 7931 | 1052 |
| J8 | *L. ovalifolium* | 2293 | 576152 | 19133185 | 9106 | 1272 |
| J9 | *H. helix* | 1655 | 142122 | 9800835 | 6369 | 907 |
| J10 | *H. helix* | 8724 | 209073 | 71609027 | 9345 | 4842 |
| J11 | *L. ovalifolium* | 14836 | 810876 | 162546003 | 13396 | 7745 |
| J12 | *Q. robur* | 4102 | 180369 | 34763423 | 9564 | 2262 |
| J13 | *A. hispanicum* | 10441 | 270088 | 110849842 | 12959 | 5462 |
| J14 | *L. ovalifolium* | 2138 | 811255 | 16979244 | 8555 | 1193 |

**Table S13.** Summary statistics of plastid genome assemblies generated by Cohort 1, prior to manual curation. Assemblies are based on the ONT data assembled with ptGAUL.

| Participant ID | Species | # contigs | Largest contig | Total length | N50 |
|---|---|---|---|---|---|
| A1 | *Q. robur* | 3 | 107470 | 178523 | 107470 |
| A2 | *L. ovalifolium* | 3 | 122225 | 185508 | 122225 |
| A3 | *H. helix* | 3 | 105676 | 168211 | 105676 |
| A4 | *A. majus* | 3 | 114368 | 184585 | 114368 |
| A5 | *L. ovalifolium* | 3 | 111693 | 181441 | 111693 |
| A6 | *B. davidii* | 3 | 102926 | 170870 | 102926 |
| A7 | *H. helix* | 3 | 113519 | 175902 | 113519 |
| A8 | *B. pendula* | 3 | 99104 | 159353 | 99104 |
| A9 | *C. coggygria* | 3 | 128962 | 197043 | 128962 |
| A10 | *P. lanceolata* | 3 | 97381 | 160008 | 97381 |
| A11 | *L. ovalifolium* | 3 | 119971 | 194448 | 119971 |
| A12 | *C. nigra* | 3 | 101281 | 168099 | 101281 |

**Table S14.** Summary statistics of plastid genome assemblies generated by Cohort 2, prior to manual curation. Assemblies are based on the ONT data assembled with ptGAUL.

| Participant ID | Species | # contigs | Largest contig | Total length | N50 |
|---|---|---|---|---|---|
| J1 | *P. lanceolata* | 3 | 94541 | 148653 | 94541 |
| J2 | *P. lanceolata* | 3 | 90767 | 147479 | 90767 |
| J3 | *C. nigra* | 3 | 116275 | 177642 | 116275 |
| J4 | *C. nigra* | 4 | 68149 | 170651 | 42662 |
| J5 | *L. corniculatus* | 3 | 106212 | 168704 | 106212 |
| J6 | *B. davidii* | 2 | 112434 | 130465 | 112434 |

| | | | | | |
|---|---|---|---|---|---|
| J7 | *C. coggygria* | 2 | 99891 | 145684 | 99891 |
| J8 | *L. ovalifolium* | 4 | 108261 | 196261 | 108261 |
| J9 | *H. helix* | 6 | 103760 | 185998 | 103760 |
| J10 | *H. helix* | 3 | 108473 | 179005 | 108473 |
| J11 | *L. ovalifolium* | 3 | 113476 | 182933 | 113476 |
| J12 | *Q. robur* | 3 | 115978 | 172248 | 115978 |
| J13 | *A. hispanicum* | 1 | 139963 | 139963 | 139963 |
| J14 | *L. ovalifolium* | 2 | 109499 | 154406 | 109499 |

**Table S15.** Quality of plastid assemblies generated with ptGAUL from the first student cohort based on comparisons to the plastids assembled from the test data. Barcode refers to the unique molecular index used by a student. Samples without relevant references are omitted.

| Barcode | Test data species | Gaps | Mismatches |
|---|---|---|---|
| Barcode01 | *Q. robur* | 68 | 3 |
| Barcode02 | *L. ovalifolium* | 88 | 0 |
| Barcode03 | *H. helix* | 94 | 0 |
| Barcode05 | *L. ovalifolium* | 84 | 0 |
| Barcode06 | *B. davidii* | 76 | 0 |
| Barcode07 | *H. helix* | 103 | 0 |
| Barcode08 | *L. corniculatus* | 602 | 0 |
| Barcode09 | *C. coggygria* | 38 | 0 |
| Barcode10 | *P. lanceolata* | 40 | 3 |
| Barcode11 | *L. ovalifolium* | 191 | 0 |
| Barcode12 | *C. nigra* | 93 | 5 |

**Table S16.** Quality of plastid assemblies generated with ptGAUL from the second student cohort based on comparisons to the plastids assembled from the test data. Barcode refers to the unique molecular index used by a student.Samples without relevant references are omitted.

| **Barcode** | **Test data species** | **Gaps** | **Mismatches** |
|---|---|---|---|
| Barcode01 | *P. lanceolata* | 72 | 6 |
| Barcode02 | *P. lanceolata* | 65 | 6 |
| Barcode03 | *C. nigra* | 138 | 6 |
| Barcode04 | *C. nigra* | 130 | 6 |
| Barcode05 | *L. corniculatus* | 708 | 4 |
| Barcode06 | *B. davidii* | 117 | 1 |
| Barcode07 | *C. coggygria* | 105 | 0 |
| Barcode08 | *L. ovalifolium* | 503 | 3 |
| Barcode09 | *H. helix* | 133 | 3 |
| Barcode10 | *H. helix* | 130 | 1 |
| Barcode11 | *L. ovalifolium* | 258 | 2 |
| Barcode12 | *Q. robur* | 117 | 1 |
| Barcode14 | *L. ovalifolium* | 265 | 17 |
