## Supplementary Protocol 2 for "A workflow for practical training in ecological genomics using Oxford Nanopore long-read sequencing"

### NERC Advanced Training in Ecological Genomics:

### Bioinformatics

**Instructors:**

- Alex Twyford, Co-Academic Lead, Edinburgh Genomics
- Urmi Trivedi, Bioinformatics Team Leader, Edinburgh Genomics
- Heleen De Weerd, Bioinformatics analyst, Edinburgh Genomics
- Tim Booth, Bioinformatics analyst, Edinburgh Genomics
- Nathan Medd, Training Manager, Edinburgh Genomics

#### Course Book Contents

#### Connecting to our training virtual machines (VMs)

**1.** First, you will need a VNC (virtual network computing) viewer running on your PC or Mac. 'TigerVNC' is our preferred choice. Please follow this link to download it now:

***https://sourceforge.net/projects/tigervnc/files/stable/1.12.0/***

*For Windows you want the file called vncviewer64-1.12.0.exe, because you only need the viewer not the full server.*

**2.** Once TigerVNC is successfully downloaded, click on it to run. You will be greeted with a grey dialogue box asking for a VNC server connection address.

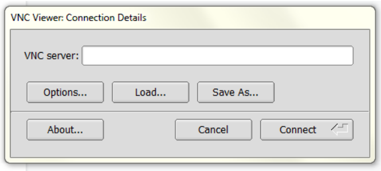

**3.** To discover the connection address your personal VM for the course, go this page:

***tinyurl.com/egtraining***

... and copy the 'Current VNC Address' next to your VM number from this page into your TigerVNC box.

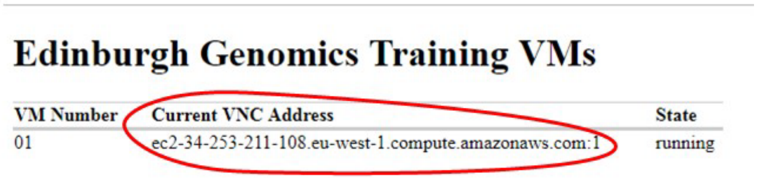

Clicked 'Connect' and when prompted enter the password **EdG3n0m1cs**. You should be greeted by a Linux desktop.

**Welcome to your VM for the course!** This VM is yours to play with throughout the course, it is a 'sandbox' that can be reset or restarted in an instant. **You can't break anything or delete anything important.**

Your VM will be shut down and rebooted overnight, so remember to **save your work and close down applications at the end of each day**. Any files or notes you want to have access to at home can be sent to yourself by email or dropped into an online folder, Google Drive, OneDrive etc. **Steps 2-3 will have to be repeated every morning of the course, because the IP addresses get recycled overnight.**

**How to use this tutorial:**

Example commands are shown like this:

```
$ ls
```

Type the command (without the $, which represents the prompt) into the terminal and press the [Enter] key to run it. For some example commands the expected output is shown directly underneath the command. When output ends with ..., it means that only a part of the real output is shown.

> **Extra information / tips**
>
> Information that is useful but not essential to the tutorial is shown like this.

In various places in the tutorial there are exercises with sample answers.

### Dry Lab: Day 1

### $ intro -to **Linux**

**Instructors:**

Tim Booth - Bioinformatician and Programmer

Nathan Medd - Training and Outreach Manager

### 1. Introduction to Linux

Genomic studies like the one we are engaged in produce vast amounts of data, usually in the form of very large text* files. Commands available on the Linux command line are particularly suited for working with such files and it is therefore arguably one of the most important tools in a bioinformatician's toolkit. The command-line enables one to search, manipulate and distil large text files that are difficult or impossible to handle with applications like Word or Excel. By chaining simple commands you can write pipelines to perform certain tasks, and run bioinformatics software for which no web or GUI interface is available.

** We'll explore the distinction between "text files" and "binary files" later.*

**In this section of the course you will learn to:**

- Navigate the file system using the Linux command shell (chapters 1.1 - 1.6)
- Use basic shell commands to view and filter data (chapters 1.7 - 1.9)
- Install new tools and use these to assess the quality of raw ONT reads (chapters 10-11)**

**You will use the following tools:**

- Ubuntu Linux with XFCE4 desktop: https://www.xubuntu.org
- Bash: https://en.wikipedia.org/wiki/Bash_(Unix_shell)
- GNU coreutils: https://www.gnu.org/software/coreutils
- Miniconda3: https://docs.conda.io/miniconda.html
- NanoPack toolkit: https://github.com/wdecoster/nanopack

#### 1.1 The shell and commands

The command-line interface in Linux is provided by a program called the **shell**. The most common shell program found in Linux distributions is **Bash** (the name is a pun on the original developer Stephen Bourne - **B**ourne **A**gain **Sh**ell). Bash is a command processor, i.e. a program that takes commands and passes them to the Linux kernel, which executes them.

The **shell** runs in a text window, where the user types **commands** and views outputs. This text window is called a **terminal emulator,** or simply a **terminal**. Bash can also read commands from a text file, called a **shell script**, in which case it doesn't necessarily need a terminal. (And you can also run other commands than a shell, like a text editor, in the terminal.)

> ➔ Open a terminal window now, by clicking the Terminal Emulator icon on the left on your Linux desktop

You can move and resize the terminal window using the mouse, or maximise it using the maximise icon, and increase and decrease the text size by selecting 'View > Zoom In' and 'View > Zoom Out' from the menu bar. You can open as many terminal windows as you like.

What you will see in your terminal window is the **command prompt**, which consists of a user name (in this case, training), the name of the machine this user is working on (eg. vm-01), and the name of the current directory (~), followed by a **$** sign that marks the end of the prompt:

```
training@vm-01:~$
```

For the sake of simplicity, and because the prompt can change, in our examples we will just use the **$** sign to represent the entire command prompt.

The basic structure of a Linux command line is:

**command [option(s)] [argument(s)]**

- command: the operation or program you want Linux to execute
- options (also called flags or switches): modify the way the command works
- arguments: filenames or other targets that direct the action of the command

The options and arguments are not always needed, when a command has some default behaviour.

Commands are executed by typing the command line at the command prompt, followed by pressing the [Enter] key (aka. the [Return] key, aka. ⏎).

Let's try two examples:

The **date** command simply displays the current date and time in the terminal. If run with no options it shows the local time, but if run with the "-u" **option** it shows UTC. By convention, options to Linux commands are always preceded by hyphens like this, while arguments are not.

```
$ date
Fri 19 Jun 15:35:03 BST 2020

$ date -u

Fri 19 Jun 14:35:20 UTC 2020
```

**Note:** You must put a space between the command name ("date") and the option ("-u"), but you must **not** put a space between the hyphen and the letter "u". Also, you must use all lower case.

- ➔ In your terminal, run the two "date" commands as shown.
- ➔ Break the rules in the note above - what happens?

The ls (list) command lists the contents of the current directory:

```
$ ls
Desktop    Public    Videos

Documents  Music     R       Templates      bin

Downloads  Pictures  RNAseq  Variants       igv
```

These are subdirectories within your home directory. If you give a directory name as an **argument** to the **ls** command it will list that directory instead.

```
$ ls Pictures
edgen_logo_lores_white.png  edgen_wallpaper.png
```

And, like the **date** command earlier, the **ls** command accepts options which modify the behaviour of the program. Here, we'll use the **-l** option to get a long listing, and the **-h** option to show file sizes in a human-readable style.

```
$ ls -l -h Pictures
total 532K

-rw-r--r-- 1 training users  24K Apr 10  2018 edgen_logo_lores_white.png

-rw-r--r-- 1 training users 505K Apr 10  2018 edgen_wallpaper.png
```

The meaning of the first few columns will be explained later, but you can see that the size of the files in kilobytes, and the date they were created, is displayed before the file name.

The shell remembers the commands you have run, and previous command lines can be recalled using the [Up arrow] key. This is particularly useful to run a similar command without writing the whole command out again.

You can move the cursor using the [Right arrow] and [Left arrow] keys. To jump directly to the beginning or end of a command, use the [Home] and [End] keys, respectively. Note that after editing the command you **don't** have to move the cursor back to the end before you press [Enter].

- ➔ Run the **ls** command line above
- ➔ Compare the result with what you see in the graphical file explorer
- ➔ Use the up arrow to recall the command, and combine the two options, so instead of "-l -h" you have "-lh". Do you see the same output?
- ➔ Type the command: **history**. What do you see?

**The mouse**

In the terminal, the mouse does not move the input cursor - you must use the arrow keys for that.

Having said this, the mouse can still be handy. Besides using the mouse to resize and scroll the terminal window, you can use it to **select and copy text**.

In this section we have learned the following commands:

| | |
|---|---|
| date | show the date and time |
| ls | list contents of a directory |
| history | Show a listing of previous command lines |

And the following concepts:

| | |
|---|---|
| prompt | A small piece of text printed by the shell indicating that it is ready to receive a command line. Conventionally ends with a $ sign. |
| command | a program or operation you tell the shell to run |
| options | settings that modify the behaviour of the command. By convention they are single letters preceded by a hyphen. |
| arguments | Items such as a file or directory you want the command to operate on |
| Command line history | Use the up arrow to recall previous commands, and the left and right arrows to edit the command. |

#### 1.2 Getting help within the shell

There are several ways to find out what a command does and what options are available for that command.

Most commands can be run with the **--help** or **-h** option to get a brief description of the command, e.g.:

```
$ ls --help
```

Some commands don't have a --help or -h option. For these commands, you can try the **help** function instead:

```
$ help cd

$ help help
```

If you run the second of those you will see that **help** "Display[s] information about builtin commands." So it only tells you what Bash knows about the command, and Bash itself only knows about basic commands.

```
$ man ls
```

The **man** command opens up a **manual page** (or "manpage") for a particular command. Manpages generally contain more detailed information than you'll get with the --help or -h flags, plus you can get information on just about any command on the system.

You can scroll through the man page one line at a time using the [Down arrow] and [Up arrow] keys, or one screen at a time using the [Page Down] key or space bar, and the [Page Up] or [B] key.

You can search in the manpage by typing a / followed by your search term and pressing [Enter]. This will highlight all occurrences of your search term in the manpage. You can use [N] and [Shift]+[N] to jump forwards and backwards between the highlighted occurrences.

Some manpages are more informative than others, but one tip when encountering a new command is to skip down to the bottom, as there are often examples of how to use the command near the end.

**To quit the manpage, use the [Q] key.** ([Ctrl]+[C] doesn't work in this case)

**Stopping a command with [Ctrl]+[C]**

When you want to kill (ie. quit immediately from) a job you're running in the terminal, use the [Ctrl]+[C] key combination.

Often this key combination fixes the problem when your terminal stops responding or when for some reason you've lost the command prompt. It can also be used to quickly abandon the current command you are typing and bring up a fresh prompt.

Note that in many applications like word processors this key combination is used to copy text, but in the shell it is used to stop programs.

In this section we have learned the following commands:

| | |
|---|---|
| *command* --help | show help for *command* |
| help *command* | show help for *command* |
| man *command* | show manual page for *command* |
| [Ctrl]+[C] | kill the current job |

#### 1.3 Files and directories

➔ Before continuing, we need to ensure some sample files are in place.
➔ Run the following command:

```
$ tar -xvaf /mnt/s3fs/tar/linux_course.tar.xz

Linux/
Linux/Downloads/
Linux/Downloads/Mus_musculus.GRCm38.dna.chromosome.Y.fa.gz
Linux/Downloads/Homo_sapiens.GRCh38.dna.primary_assembly.fa.gz
Linux/Downloads/SRR026762.fastq.gz
…
```

You should see a list of files being unpacked and copied, as above. If you get an error, check that you typed the command exactly as shown, including the spaces. Remember you can recall the command using the up arrow to edit it. It's not a problem if you run this command more than once.

Files on the Linux system are grouped into **directories** (also known as folders, just as in a Windows or Mac environment). The directories are arranged in a hierarchical tree structure. At the top is the **root directory** "/" that holds everything else. The root directory contains **files** and **subdirectories**. The subdirectories in turn can contain files and subdirectories, and so on.

Below is a small part of the directory structure on our system.

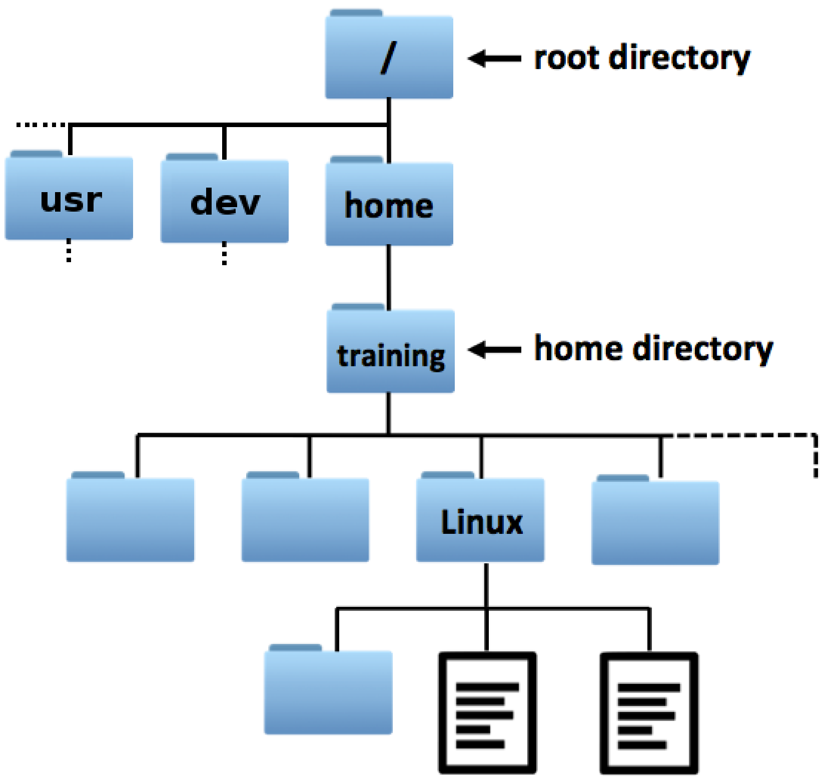

To identify any file or directory we can give an **absolute path** to that file by starting from the root and working down. These paths are written as a list of directory names separated by forward-slash characters.

In the command above, for example, /mnt/s3fs/tar/linux_course.tar.xz refers to a file named **linux_course.tar.xz** within a **tar** subdirectory within a **s3fs** subdirectory within a **mnt** subdirectory which lives in the "**/**" directory. (As it happens, the **s3fs** subdirectory is on a shared network drive. In Linux, all files are accessed within the same hierarchical structure, no matter what physical disk they are stored on.)

To avoid having to type out the full path to every file you work with, the shell maintains a current **working directory** and any commands, like `ls`, will run in that context.

```
$ pwd

/home/training
```

"pwd" means "print working directory". When you first start a shell, your **working directory** will be your **home directory**. In our case this is /home/training. That is, the **training** directory within the **home** directory within the root directory.

Note that, although there is a directory named simply **home**, this is **not** your home directory, but **/home/training** is!

If you refer to the name of a file or directory and don't start the path with a "/" then Linux will resolve the path **relative to** the working directory. So the following commands do the same thing.

```
$ ls /home/training/Linux

Downloads          mouse_exons.bed   Mus_musculus_snps_19_1_20000000.vcf
      sequence_alignment       drosophila_species.txt      toy.sam
      drosophila_species_cleaned.txt

$ ls Linux

Downloads   mouse_exons.bed   Mus_musculus_snps_19_1_20000000.vcf
sequence_alignment       drosophila_species.txt      toy.sam
      drosophila_species_cleaned.txt
```

💡 **Best practices for choosing good file and directory names**

1. Everything in Linux is **case sensitive**. If in doubt, stick to **lower case**.

The filename myfile is **not** the same as Myfile, MyFile, or myFile. Similarly, typing the command pwd is different to typing PWD.

2. File and directory names should **only contain letters**, **numbers**, **hyphens**, **underscores**, and **full stops**.

3. File and directory names should **not contain any spaces**.

In contrast to graphical environments, the command-line interface can become cumbersome when your file names contain spaces or special characters. For example, the shell will split the filename my file.txt to be two arguments, i.e. "my" and "file.txt". To use this filename, you will need to enclose the entire filename in quotation marks or escape the space with a backslash, so that the shell understands that the space is part of the name:

```
“my file”  or  my\ file
```

So, it’s best to try to avoid spaces and unusual characters altogether, even though Linux will not prevent you from putting them into file names.

In this section we have learned the following command:

| | |
|---|---|
| pwd | print working directory |

And the following concepts:

| | |
|---|---|
| Home directory | The place in the filesystem heirarchy where a given user on a Linux system keeps their own files |
| The root directory - “/” | The top of the unified file system heirarchy |
| Absolute path - eg “/home/training” | The location of a file, fully specified by starting at the root directory |
| Relative path - eg. “Linux/toy.sam” | A file location not beginning with “/” is interpreted relative to the current working directory |

#### 1.4 Navigating the file system

With the **cd** (change directory) command you can go to another working directory:

```
$ cd Linux
```

```
$ pwd
```

```
/home/training/Linux
```

Note that the shell command prompt now has changed,

from training@vm-01:~$ to training@vm-01:Linux$

**Tab completion**

You can save yourself a lot of typing by using **tab completion**. Tab completion means that partially typed file names are automatically filled in by pressing the [Tab] key. In the example above, if you type only cd L and then press the [Tab] key, your command will automatically be completed to cd Linux/ because Linux is the only subdirectory starting with the letter L. If nothing happens after pressing the [Tab] key, it may mean that there are several options to complete your command. For example, if you type cd D and then press the [Tab] key, nothing will happen, because there are several subdirectories that start with the letter D. Pressing the [Tab] key again will show all these options: Desktop/ Documents/ Downloads/. For successful tab completion, in this case, you have to type some more letters.

➔ Practise some tab completions, and try to use it from now on, as it will save you a lot of time and also will prevent you from making typos!

Use **cd ..** to go to the directory above your current directory (i.e. its **parent directory**). In this case, this puts us back in the home directory:

```
$ cd ..
$ pwd
/home/training
```

And if you repeat the **cd ..** command twice more this takes us right up to the file system root:

```
$ cd ..
$ cd ..
$ pwd
/
```

To go straight back to your home directory, use `cd` without any argument:

```
$ cd
$ pwd
/home/training
```

You may have noticed that the prompt uses a **~** character to indicate you are in your home directory. This is a standard shell shorthand, so **cd ~** is an equivalent command that takes you directly home.

In the previous chapter we mentioned **absolute and relative paths**. Remember an **absolute path** specifies a location from the root of the filesystem, starts with a **/**, and will always be valid - while a **relative path** specifies a location starting from the current location, does not start with a **/**, and will no longer be valid if you change directory.

The simplest and most common form of relative path is just the name of a single file or subdirectory in the current directory, but you can also chain multiple directory names and the "**..**" path element.

To demonstrate, first go the `Linux` directory within your home directory:

```
$ cd Linux
```

To go from here to /home/training/Downloads in a single step, we can can construct a path that goes back up to the parent directory in order to find the Downloads directory.

```
$ cd ../Downloads
```

This will work from the Linux directory, or any directory which is in /home/training, but if we started from somewhere else we'd need to construct a different route, or type the full absolute path, or use the **"~"** shorthand.

```
$ cd ~/Downloads
```

Which is equivalent to:

```
$ cd /home/training/Downloads
```

One common gotcha is this:

```
$ cd ~/Linux

$ ls

...list of files, as expected...

$ ls Linux

ls: cannot access 'Linux': No such file or directory

$ cd Linux

bash: cd: Linux: No such file or directory
```

If you are already working inside a directory, then referring to that directory by name doesn't work, as shown above. This is because the rules of relative paths say that they are relative to the working directory, and here the Linux directory is not **within** the working directory it **is** the working directory. The final two commands above refer to a directory named Linux *within* the directory named Linux, which we could create, but right now there is no such directory.

Finally, the single dot **.** is a shorthand for the working (current) directory, just as **..** means the parent directory. This seems pretty redundant now but we'll find a use for it later.

```
$ cd ./././

# Running this changes nothing!
```

💡 **Linux does not need to be in any given directory to access a file**

A common misconception for beginners is that Linux can only "see" a file if you cd to the directory where the file is stored, but really the ability to change working directory is only to avoid you having to type out long absolute paths. You can get Linux to generate an absolute path to any file with the **realpath** command:

```
$ cd ~/Linux

$ realpath toy.sam

/home/training/Linux/toy.sam
```

In this section we have learned the following commands and symbols:

| | |
|---|---|
| cd *dir* | change directory to subdirectory *dir* |

| | |
|---|---|
| cd .. | change directory to parent directory |
| cd | change directory to home directory |
| ~ | shorthand for home directory |
| . | shorthand for working (current) directory |

##### 1.4.1 Exercises on file system navigation

(1) Open a new shell window, which will put you in your home directory.

(a) List the contents of the sequence_alignment subdirectory within the Linux directory. Do you need to use `cd` to do this?

(b) What about changing to the Linux directory and then listing the contents of your home directory without changing back?

(2) Using --help, learn how the -a option affects the ls command.

(3) Using the man command, find the option to sort the list of files by modification time when using ls command (Hint: search the ls man page for the word “modification”).

(4) Look at the diagram showing the directory structure in section 3. Write out the **absolute** paths to the following directories: home (i.e. the directory with the name “home”), dev, training, and Linux.

(5) Similar to above, starting from your home directory (/home/training), write out the **relative** paths to the following directories: home (i.e. the directory with the name “home”), dev, training, and Linux.

##### 1.4.2 Answers to exercises

(1)(a)

```
$ cd Linux
$ cd sequence_alignment
$ ls
```

or, without changing directory

```
$ ls Linux/sequence_alignment
```

(1)(b)

```
$ cd ~/Linux
$ ls ..
```

or ls ~ or ls /home/training

(2)

```
$ ls --help
...
-a, --all  do not ignore entries starting with .
...
```

**Tip**: try running ls -a in your home directory.

Files and directories starting with a . are not shown unless the -a flag is used. They are referred to as hidden files and directories (or dotfiles). Aside from the directory shortcuts . and .. these mostly contain configuration settings and temporary files belonging to applications you run.

(3)

```
$ man ls
...
-t     sort by modification time, newest first
...
```

Remember - to quit the manpage, use the [Q] key.

**Tip**: try running ls -alt in your home directory.

(4)

home: /home

dev: /dev

training: /home/training

Linux: /home/training/Linux

(5)

home: ..

dev: ../../dev

training: .

Linux: Linux

#### 1.5 Moving, copying and deleting files

**Reminder**

Make sure that you are in the directory ~/Linux when starting a new chapter, unless it's explicitly stated that you have to be in another directory!

```
$ cd ~/Linux
```

Now we will have a look at how to create, copy, move, and remove files and directories within the shell.

First create two new directories, temp1 and temp2, using the **mkdir** (make directory) command:

```
$ mkdir temp1 temp2
```

And check whether the directories have indeed been created:

```
$ ls
temp1  temp2 ...
```

Change to the temp1 directory:

```
$ cd temp1
```

Create two (empty) files, myfile1.txt and myfile2.txt, using the **touch** command:

```
$ touch myfile1.txt myfile2.txt
```

And check whether the files have been created:

```
$ ls
myfile1.txt myfile2.txt
```

Files can be copied using the **cp** (copy) command:

```
$ cp myfile1.txt ~/Linux/temp2
```

Because we copied it, the `temp1` dirbectory still contains the file `myfile1.txt`:

```
$ ls
```

```
myfile1.txt myfile2.txt
```

Files can be moved using the **mv** (move) command:

```
$ mv myfile2.txt ~/Linux/temp2
```

Because we moved it, the temp1 directory doesn't contain the file myfile2.txt anymore:

```
$ ls
myfile1.txt
```

The mv command is also the command we use to **rename** files:

```
$ mv myfile1.txt myfile1_renamed.txt

$ ls
myfile1_renamed.txt
```

**Specifying cp and mv targets**

The cp and mv commands are entered in the format:

command *source target* (e.g. cp myfile1.txt ../temp2).

Note that:

If the target is an **existing directory**, the command will create a file with the same name as the source in the target directory. In this case you may have multiple source files.

If the target is an **existing file**, the command will **overwrite** the target file.

If the target **does not exist**, the command will create a new file with that name.

Files can be removed using the **rm** (remove) command:

```
$ rm myfile1_renamed.txt

$ ls
```

Go back to the directory ~/Linux:

```
$ cd ~/Linux
```

Directories can be removed using the **rmdir** (remove directory) command. However, rmdir can only remove empty directories.

Thus, directory temp1 can be removed:

```
$ rmdir temp1

$ ls
temp2
```

But directory Temp2 cannot be removed this way, as it contains two files:

```
$ rmdir temp2
rmdir: failed to remove 'temp2': Directory not empty

$ ls
temp2
```

**No news is good news**

You'll see that the above rmdir command showed an error message, but the previous commands (touch, rm, mv, cd, rmdir) just returned straight to the prompt. This is a tenet of the classic UNIX design - if a command worked as expected and has nothing unusual to report it says nothing. The core system commands all follow this pattern.

Often you can add a **-v** flag to get a verbose output, saying what was done.

One way to remove a non-empty directory is to first remove all the files it contains using rm, and then the directory itself using rmdir.

A quicker way is to remove the directory and its contents in one go using rm with the **-r** (i.e. recursive) option:

```
$ rm -r temp2

$ ls
```

So, rmdir may seem redundant but it is safer than rm -r, in that it will never remove any files, only empty folders.

In this section we have learned the following commands:

| | |
|---|---|
| mkdir *dir1* [*dir2...*] | create directory *dir1* [*dir2...*] |
| touch *file1* [*file2...*] | create empty file *file1* [*file2...*] |
| cp *file1 dir1* | copy *file1* to *dir1* |
| mv *file1 dir1* | move *file1* into *dir1* |
| mv *file1 file2* | rename *file1* to *file2* |
| rmdir *dir1* [*dir2...*] | remove empty directory *dir1* [*dir2...*] |
| rm *file1* [*file2...*] | remove *file1* [*file2...*] |
| rm -r *dir1* [*dir2...*] | recursively remove directory *dir1* [*dir2...*] and its contents |

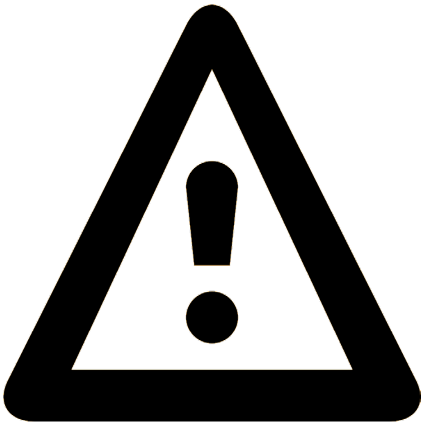

***Warning***

*When you remove or overwrite files from the command-line, they are gone. Forever. They are not put in a Trash or Recycle bin like in a graphical environment, so there is no easy way to get them back!*

You can of course make your own ~/Trash directory and move things to there instead of deleting them, then empty out the trash when you are sure. This is good practise.

You can also manipulate files via the Graphical User Interface (GUI) browser (we won't tell!) but this may not always be available, and when you are already in the right directory in your shell it is quicker to type a command than navigate in the GUI.

Finally you can add the -i flag to cp, mv and rm so you will be specifically asked if you really want to remove each file. “-i” here is short for “interactive” On some systems this may be set as the default, but you should never rely on it.

Get into the habit of using the safe rmdir when you think the directory is empty, and triple-checking whenever you use rm -r. A “classic mistake” is to type something like:

```
$ rm -r ~/myproject /temp_files.
```

That extra space in the command means the entire ~/myproject directory is going to silently vanish, followed by an error message saying that /temp_files cannot be found.

The danger is especially high if you are in a root shell (an administrative shell where file permissions, as described below, are not enforced). People have destroyed their whole systems with a single mistyped rm -r command. After this, recovery from backup is the only option.

#### 1.6 Shell "glob" patterns and wildcards

Sometimes, you want to operate on multiple files at once. The Bash shell supports wildcard characters "*" and "?" to enable this.

```
$ cd Linux
```

```
$ ls *.txt
drosophila_species.txt  drosophila_species_cleaned.txt
```

The * can represent any number of any characters. There are two files in the directory that end with .txt and so the shell expands the pattern for us and passes both the filenames as arguments to the **ls** command. It's important to note that this is a feature of the shell itself, not the **ls** program. A single * expands to all the filenames and subdirectory names in the current directory.

```
$ ls *

training@vm-01:Linux$ ls *
Mus_musculus_snps_19_1_20000000.vcf  mouse_exons.bed
drosophila_species.txt               toy.sam
drosophila_species_cleaned.txt

Downloads:
Homo_sapiens.GRCh38.dna.primary_assembly.fa.gz  SRR026762.fastq.gz
Mus_musculus.GRCm38.dna.chromosome.Y.fa.gz

sequence_alignment:
6991_1.fastq.gz
6991_2.fastq.gz
Escherichia_coli_bw25113.ASM75055v1.dna.toplevel.fa.gz
```

Here, the * expands to the names of the 5 files plus the 2 directories. The **ls** command prints back the filenames, and lists the contents of the directories (but would not list any subdirectories inside them).

The ? character is like * but matches only a single character. If you still have the temp1 and temp2 directories from the previous chapter you could say:

$ ls temp?
...or...
$ rmdir temp?

💡These multi-file matching patterns are called "glob" patterns, which is simply short for "global" because they are used to apply a single operation globally over many files.

A number of tools and programming languages aside from the shell support this type of pattern.

##### 1.6.1 Exercises on file management

(1) Go to the directory ~/Linux. In this directory, create a directory named directory1.

Within directory1, create two subdirectories, named subdirectory1 and subdirectory2.

In subirectory1, create two empty files, named file1 and file2.

Use the **tree** command to visualise what you have created. We've not used this command yet, but it's basically a version of **ls** that lists directories and their contents at once. You could use the relevant manpage to discover how it works, or just run it and see. You should see something a little like this:

```
/home/training/Linux/
...
 directory1/
  subdirectory1/
   file1
   file2
  subdirectory2/
```

(2) Copy file1 and file2 from subdirectory1 to subdirectory2. (**hint** - if you want to copy both files in one go, you can use the * or ? wildcard). Check whether both files have been copied.

(3) Stay in the directory ~/Linux. Remove subdirectory1 and the files it contains with a single command. Check whether the directory has been deleted.

##### 1.6.2 Answers to exercises

(1) This is one way to do it...

```
$ cd ~/Linux
$ mkdir directory1
$ cd directory1
$ mkdir subdirectory1
$ mkdir subdirectory2
$ cd subdirectory1
$ touch file1
$ touch file2

$ cd ~/Linux
$ tree directory1
...
```

It is possible to make all the directories and files in just two commands, using a shell feature we've not seen yet; nobody would expect you to come up with the following. But to show it's possible:

```
$ mkdir -p directory1/subdirectory{1,2}
$ touch directory1/subdirectory1/file{1,2}
```

(2) Again, this is just one possible way...

```
$ cd ~/Linux/directory1/subdirectory1
$ cp file1 ../subdirectory2
$ cp file2 ../subdirectory2
```

or with the ? wildcard

```
$ cp file? ../subdirectory2
$ ls ../subdirectory2
file1 file2
```

(3)

```
$ rm -r directory1/subdirectory1
```

or (two commands, but safer!):

```
$ rm directory1/subdirectory1/*
$ rmdir directory1/subdirectory1
```

#### 1.7 Viewing text files

So far we've moved empty files around and completely ignored the file contents. Let's finally take a look at some sequence data, and introduce some core Linux commands to access text files.

```
$ cd /mnt/s3fs/NERC_training/demo_data/misc/
$ ls
small.fastq
$ head small.fastq
@HWUSI-EAS721_0001:8:1:1021:5705#0/1
GATATTCAGCCATACCATGCTNATCCTCGGGATCGCNGTGATCCT
+
CCCCCCCCCCCCCCCCCCCAA#AAAAAAACCCCC>?#?>?<?;<C
@HWUSI-EAS721_0001:8:1:1021:10177#0/1
TGGTCACCGGTTGCAGAGTAANGCTCATCTCTTCTTNACCAGGGA
+
CCCCCCCCCCCCCACCCCA??#=AAAAAACCCCC??#A?;:9;;C
@HWUSI-EAS721_0001:8:1:1022:12400#0/1
CCACCGGTCCAGACAAAATCANCCACTTCCAGATCGCACTGCTGC
```

The FASTQ format is a standard way of storing raw sequencer reads. Each read is represented by four lines of text:

1. a header line, beginning with @
2. the sequence itself
3. a '+' spacer line
4. a string of gibberish-looking characters encoding the quality of each basecall

Since the FASTQ format is a type of text file and not a binary format we can display and filter it using standard Linux tools, even though none of these tools were written specifically with FASTQ files in mind. The **head** command above prints the first few lines of the file to the terminal. By default, 10 lines are printed so we are seeing the first two sequence records and half of the third.

This is very useful to peek at a large file. If you want to see the whole file you can use the **cat** (concatenate) command, but if the file is large it will just whizz up the screen too fast to read.

```
$ cat small.fastq
...lots of output...
```

The **less** command (the name is another pun - an improved version of the older **more** command) allows you to look at the file content one screen at a time:

```
$ less mouse_exons.bed
```

You have used the **less** command already, indirectly, because it is the text viewer used to read **man** pages. You can scroll around with the [Down arrow] and [Up arrow] keys, get help by pressing [H], and quit by pressing [Q].

Much like **head,** the **tail** command just returns the last 10 lines of the file, or you can specify the number of lines using the **-n** option:

```
$ tail -n 4 small.fastq
@HWUSI-EAS721_0001:8:1:1042:16579#0/1
TTCAGTACATTGTTGACGAGATTGTGGCTGCAGGGATCAAAGAAA
+
CCCCCCCCCCCCCCCCCCCCCCCCCCCCCC>?CCCBCCCCCC?CB
```

The **wc** (word count) command returns the number of lines, words, and bytes in a file:

```
$ wc small.fastq
 100  100 3292 small.fastq
```

If you're only interested in the number of lines, add the `-l` (lines) option:

```
$ wc -l small.fastq
 100 small.fastq
```

As we know there are 4 lines of text per sequence, this file must threfore contain 25 sequences.

In this section we have learned the following commands:

| | |
|---|---|
| cat *file* | output content of *file* |
| less *file* | output content of *file* one screen at a time |
| head *file* | output first 10 lines of *file* |
| head -n *num file* | output first *num* lines of *file* |
| tail *file* | output last 10 lines of *file* |
| tail -n *num file* | output last *num* lines of *file* |
| wc *file* | print number of lines, words, and bytes in *file* |
| wc -l *file* | print number of lines in *file* |

#### 1.8 Symbolic links

Alternatively referred to as a **soft link** or **symlink**, a **symbolic link** is a pseudo-file that links to another file or directory (called the **target** of the link).

Symbolic links are created using the **ln** command with the **-s** option. The **-r** option is also useful as it makes **ln** resolve paths automatically when making the link:

```
$ ln -sr ~/Linux/mouse_exons.bed ~/Public
```

Symlinks are shown in long-format directory listings, with an `l` in the first character and a `->` annotation that gives the target name. The target may be a relative or an absolute path, and relative paths are always relative to the directory containing the symlink:

```
$ ls -l ~/Public

lrwxrwxrwx 1 training training 29 Feb 27 13:55 mouse_exons.bed
-> ../Linux/mouse_exons.bed
```

💡 We're not covered permissions yet, but the "rwx…" part in the listing relates to the file permissions, and since symbolic links have no permissions of their own, so they show up as "lrwxrwxrwx".

If you try to read or edit the link, you will end up reading or editing the contents of the file where the link points. Deleting the symbolic link does not affect the target file. If, however, the target file to which the link points is removed or renamed, the link will stop working. This situation is called a "dangling symlink".

It is possible, and sometimes useful, to make symlinks to directories rather than files, and you can also make a link to to a link in which case Linux will follow the chain until it gets to a real file (or a hits a dangling link). However, things can get confusing if you have too many links like this. There is a **realpath** command which, for any file, will resolve the links and give you the "canonical", ie. link-free, absolute path to a file.

```
$ realpath Public/mouse_exons.bed
/home/training/Linux/mouse_exons.bed
```

In this section we have learned the following commands:

| | |
|---|---|
| ln -sr *file1 dir1* | Assuming dir1 exists, make a symbolic link to *file1* in *dir1* |
| ln -sr *file1 link1* | Assuming link1 does not exists, create a symlink named link1 pointing to file1 |
| realpath *file1* | Show the canonical absolute path to any file, resolving all links. |

#### 1.9 Zipping and unzipping files

When typing the commands below, remember to use **tab-completion** to avoid typing the long filenames in full, and note that several command lines are shown wrapped over two lines because they are too long to print on one line, but you need to type them as one line.

```
$ cd /mnt/s3fs/NERC_training/demo_data/raw_data/
$ cd 24130TA0003_03
$ ls
20221221_EGS2_24130TApool03_24130TApool03_PAM37551_2f2823fb_bar
code03_pass.fastq.gz
$ less 20221221_EGS2_24130TApool03_24130TApool03_PAM37551_2f2823
fb_barcode03_pass.fastq.gz
```

Like the file we saw in the previous chapter, this file also contains FASTQ sequence records. It's a lot bigger, both in terms of the number of sequences and their length, but the format is the same.

However, this file is also compressed with **gzip** (GNU zip). The **fastq.gz** extension has been used to indicate the zipped format. Compressed files use less disk space to store and can be transferred faster, and are commonly encountered in bioinformatics.

After quitting from **less** (press Q), try to use **head** on the same file:

```
$ head -n 6 20221221_EGS2_24130TApool03_24130TApool03_PAM37551_
2f2823fb_barcode03_pass.fastq.gz
*�����*1u%y�Y���6���K�>�:�x��K�|Ҳ�5j1z-
�8���`��Db�J���M������i��>i�6i�2n�&;$��n����a�������
0v������?���.��J��]����^�\
%¾(>�?������w1��o��F�N�h�wTon�B���y.����m��$��t��
�B�Ղ�)u�_}��3���RH�����9=ℸ
���a��wW��z����.��/��:Rj����JH���6�9RS�%���0�X�s�
�6ob���L<A-
��:��,�_n�1~n�kZ�}����R�X���Ut�',�wU�l��\b���hee�]ŵ���:
<�[�ő��OK[uau@dR�}�g   "C���u5���Hav�&k�&E���@QQKg�:�|)ol�
...
```

Ewwww! The **head** command, unlike the **less** command, does not know about gzipped files and just shows the compressed binary data directly. Printing a binary file to the terminal is neither useful nor pretty, and using **wc** to try and count the lines will give us a nonsense result. We'll need to unzip the file.

First, copy the file to your home directory (remembering that ~ is shorthand for home):

```
$ cp 20221221_EGS2_24130TApool03_24130TApool03_PAM37551_2f2823fb
_barcode03_pass.fastq.gz ~
```

This takes a little while as it's a big file. Have a look at the size of the file:

```
$ cd ~
$ ls -lh
...
-rwxrwxr-x 1 training users 2.3G Apr 7 10:21
20221221_EGS2_24130TApool03_24130TApool03_PAM37551_2f2823fb_barcode03_pas
s.fastq.gz
...
```

Note that with the option -h (human readable format) file/directory sizes are shown using unit suffixes: Byte, Kilobyte, Megabyte, Gigabyte, etc. So this file is taking up 2.3 gigabytes on disk.

Unzip (de-compress) the file using the **gunzip** command:

```
$ gunzip 20221221_EGS2_24130TApool03_24130TApool03_PAM37551_2f282
3fb_barcode03_pass.fastq.gz
```

This should take a minute or so.

**Questions:**

1. How big is the uncompressed file?
2. Confirm that you can now use the head command to peek at the first few lines of the file. What are the last 6 bases of the 10th sequence?
3. How many sequence records are in this FASTQ file?

You can zip the file again using the **gzip** command, but note that this will take several minutes:

```
$ gzip 20221221_EGS2_24130TApool03_24130TApool03_PAM37551_2f2823fb
_barcode03_pass.fastq
```

In most cases it is unnecessary to **gunzip** a compressed file before reading the contents, because we can combine the command **zcat** with other commands in one go, as we'll see in the next chapter.

Like cat, zcat prints the whole content of the file to the screen, but it decompresses it on-the-fly. You can try running zcat on this sample file but it will take a long time to print out as the file is large.

To interrupt the process, use [Ctrl]+[C].

A **tar** (tape archive) file, with extension .tar, is another common file format whereby entire directory structures and all of the files within them have been placed into a single file. Normally these are then compressed with **gzip** so you get a file with the extension .tar.gz (sometimes .tgz). Tar files, compressed or not, can be extracted using the **tar -xaf** command, eg:

```
$ wget https://ftp.gnu.org/gnu/tar/tar-latest.tar.gz
$ tar -xaf tar-latest.tar.gz
```

where: **-x** means extract files from an archive, **-a** automatically handles the decompression and **–f** specifies the archive file to read. It's also normal to add **-v**, the flag verbose for verbose output, so you get to see a list of what is being unpacked.

**Other archive file formats**

Linux tar files perform exactly the same function as zip files more normally used in Windows, and Linux can also pack and unpack .zip files as well as any other compressed format you are likely to encounter. Likewise, software to handle tar files is widely available for Mac and Windows so you should never have a problem exchanging files whatever format you use.

A reason for sticking to tar format on Linux is that tar knows about file ownership, permissions, symlinks and other quirks of the Linux filesystem.

In this section we have learned the following commands:

| | |
|---|---|
| gunzip *file.gz* | decompress *file.gz* to file |
| gzip *file.gz* | compress *file* to file.gz |
| zcat *file.gz* | output content of *file.gz* |
| tar -xvaf *file.tar.gz* | extract *file.tar.gz* listing the files as they are extracted |

#### 1.10 Pipes and redirects

Some of the real power of the Linux shell comes from the ability to string commands together to form "shell pipelines".

With the **|** (pipe) operator (which you can find on most keyboards at the bottom left next to the [Shift] key), the output of one command, written on the left of the pipe, can be used as input for another command, placed on the right of the pipe. The intermediate output is never saved to disk, it is passed directly from one command to another.

For example, if you want to count up the number of lines in a gzipped FASTQ file (see previous chapter) you can do this by first using **zcat** to get the real uncompressed content of the file, and then **piping** the output of **zcat** into **wc** to count the lines in the file:

```
$ zcat ~/Linux/sequence_alignment/6991_1.fastq.gz | wc -l
```

If you simply run **wc** on the original file, you'll get a result but it will be nonsense as wc does not know how to decompress the file before counting the contents. You might think that the **wc** command should be enhanced to work on gzipped files, as with **less**, but with pipes there is no need; you can simply join the commands together to make your own.

With the **>** (redirect) operator, the output of a command can be saved in a file, rather than being shown in the terminal.

For a simple example, let's say we want to create a file my_commands.txt, that contains the text "My commands:", followed by the contents of our command history.

To this end, first write a line saying "My commands:" to a new file. This can be done using the **echo** command, which simply prints out any text that you give it, then redirect this to the file:

```
$ echo My commands: > my_commands.txt
```

To check this text really is in the my_commands.txt file, cat it to the screen:

```
$ cat my_commands.txt
My commands:
```

If we now add the contents of our command history to the file using the > operator, it turns out that the original content of the file is overwritten and the first line is gone:

```
$ history > my_commands.txt
$ head my_commands.txt
   1 echo Hello World!
   2 ls
   3 history
   ...
```

To add output to the contents of an already existing file without overwriting it, we should use the **>>** (append) operator instead:

```
$ echo My commands: > my_commands.txt
$ history >> my_commands.txt
$ head my_commands.txt
My commands:
   1 echo Hello World!
   2 ls
   ...
```

Multiple pipes and redirection can be combined on one line:

```
$ echo My last 6 commands in reverse order: > my_commands.txt
$ history | tail -n 6 | tac >> my_commands.txt
$ cat my_commands.txt
My last 6 commands in reverse order:
  287  history | tail -n 6 >> my_commands.txt
  286  echo My last 6 commands: > my_commands.txt
  285  yes Linux is great
  284  head my_commands.txt
  283  history >> my_commands.txt
  282  echo My commands: > my_commands.txt
```

In this section we have learned the following shell features:

| | |
|---|---|
| *echo any text you like* | print back the given text (without redirection it will just appear in the terminal) |
| *command1 \| command2* | use output of *command1* as input for *command2* |
| *command > file* | redirect output of *command* to *file* |
| *command >> file* | append output of *command* to end of *file* |
| *tac file* | print the lines of file in reverse (backwards cat - not a commonly used command but sometimes handy) |

##### 1.10.1 Exercise - looking at your own data

It's now time to find your own sequence data file. Look under /mnt/s3fs/NERC_training/

```
$ cd /mnt/s3fs/NERC_training/
$ ls
```

You'll see demo_data and reference directories, as well as a **dated directory** for this specific course. Change into this dated directory, and then into the raw_data directory within it. Look for the subdirectory with your own sample number. Within this you'll see your raw sequences in a single .fastq.gz file. You may also see an .md5 file; you can ignore this.

Answer the following without using the **gunzip** command. You will need to use **zcat** with a pipe to another command.

1. How many lines (and therefore sequence records) are in this FASTQ file?
2. What are the last 6 bases of the 100th sequence?

💡 If you want to do simple arithmetic right in the shell you can use the **bc** command like so:

```
$ echo 4000 / 4 | bc
1000
```

It's possible to count the number of lines in the file and divide the result by 4 in a single command line, but we do not yet know enough shell syntax to make this work, so you'll need to just copy and paste the number into the new command line.

#### 1.11 Searching files with grep

The **grep** command in Linux is used to scan text files for any pattern you supply. We'll be using it later in the course. To try it, first take a look at the file **drosophila_species_cleaned.txt.**

```
$ cd ~/Linux
$ less drosophila_species_cleaned.txt

(remember to press 'q' to quit the viewer)
```

This text file contains a list of drosophila species (copied from Wikipedia). The name of the species is listed, as well as the entomologist who named the species, and the year. Which new drosophila species were named in 1980?

```
$ grep 1980 drosophila_species_cleaned.txt
D. altissima - Tsacas, 1980
D. anisoctena - Tsacas, 1980
D. bahunde - Tsacas, 1980
D. bakondjo - Tsacas, 1980
…
```

The grep command is searching for the text "1980", and if this appears one or more times within a line it prints the whole line. If the text is not found then it does not print that line. We can combine the above command with a pipe and **wc** to count the lines and tell us how many species were named in 1980.

```
$ grep 1980 drosophila_species_cleaned.txt | wc -l
19
```

Or, how many were named by Hardy?

```
$ grep Hardy drosophila_species_cleaned.txt | wc -l
317
```

D. Elmo Hardy was a busy man!

> 💡 **grep** does a lot more than searching for simple fixed strings. It supports a powerful pattern language called **regular expressions** - grep is short for "global regular expression print". Learning regular expressions is beyond the scope of todays course but they are well worth knowing about and there is more info in the bonus workbook.

#### 1.12 Installing software with Bioconda

It's time to use some dedicated bioinformatics software to look at the quality of the raw sequence data. So far, we've been using core tools that come with the Linux distribution, and there are also a bunch of extra tools installed on the teaching VMs for you, but often in bioinformatics you'll need to install your own software. Installing software can be surprisingly problematic, and the Bioconda project exists to try and make it easier by maintaining a catalogue of thousands of free software packages.

You can find the Bioconda home page at https://bioconda.github.io/

To get it set up on the VM, we're going to use some slightly different instructions compared to those on the official Bioconda site, because we think these work better. We'll use an installer that bundles two programs called **conda** and **mamba.** In the command shell:

```
$ cd
$ wget https://github.com/conda-forge/miniforge/releases/latest/download/Mambaforge-
Linux-x86_64.sh
...
Saving to: 'Mambaforge-Linux-x86_64.sh'
...
$ bash Mambaforge-Linux-x86_64.sh
```

The **wget** command above downloads files from the web on the command line. You could also obtain this file by using a web browser search for "mambaforge" then follow the links to find the download. Sadly, tab completion cannot help with web links, so be careful typing the address.

Accept the licence (press space to scroll through it) and press Enter to accept the default install location. When asked if you want to "run conda init" say "yes".

💡If you miss this last option it's not really a problem. Just run this at the shell prompt:

```
$ mambaforge/bin/conda init
```

Note the advice to "close and re-open your current shell". Do this now. Your prompt should now start with "(base)" indicating that conda+mamba is working.

##### 1.12.1 Setting up Bioconda and installing the packages

There are some setup instructions on https://bioconda.github.io/, but because we're using Mamabaforge we only need to run one of the commands:

```
$ conda config --add channels bioconda
```

You shouldn't see any output from this command.

This setup only has to be done once, and after this Mamabaforge will be ready to install bioinformatics packages using the mamba installer. We'll install part of the NanoPack toolset[1].

```
$ mamba install nanocomp==1.21.0 nanoplot==1.41.0 chopper==0.5.0 pandas==1.5.3
```

Mamba will have a think about what dependent packages it needs to make all this work. This takes about 2 minutes. Press Y to download and install everything. Sometimes, mamba will decide it can't make all the packages play nicely together, but in this case it should all be fine. There may well be more recent versions of the nanocomp/nanoplot packages available but we've tested these versions for this course.

> 💡You might well be wondering what "pandas==1.5.3" has to do with anything in the command above. Pandas is an extension to the Python language for data science, and is needed by the NanoPlot tool we want to run. When testing the course materials we discovered that Mamba was installing Pandas 2.0 and this is incompatible with NanoPlot 1.41, resulting in the program crashing. The workaround is to tell Mamba explicitly to install the older version of Pandas.
>
> At some point soon this will likely be fixed, but for now we have this workaround. It's not uncommon to hit such snags with bioinformatics software, and the solution for a new bioinformatician is to not be disheartened and make use of on-line forums to access help and find solutions.

Now we can try running the commands:

```
$ NanoComp -–help
$ NanoPlot --help
```

This reassures us the programs were installed and the commands are available. Even though we only asked for NanoComp, the NanoPlot package has been installed as a dependency.
To see where the programs actually got installed, we can use the **which** command.

```
$ which NanoPlot
/home/training/mambaforge/bin/NanoPlot
```

[1]https://github.com/wdecoster/nanopack

```
$ echo $PATH
/home/training/mambaforge/bin:/home/training/mambaforge/condabin:/home/training/bin:/usr/local/sbin:/usr/local/bin:/usr/sbin:/usr/bin:/sbin:/bin:/snap/bin
```

So the NanoPlot program has gone into the mambaforge/bin directory in /home/training. The reason that we can run this program without typing the full location into the command is that Conda has added this directory to the default PATH, and we can see what the PATH is by running the second command.

The PATH setting in the shell is a list of locations, separated by colons. You'll see that each is an absolute path to a place where the shell can look for programs. All the programs we've run so far, including Bash itself, are in one of these directories, or else they are built into the Bash shell.

```
$ which ls
/bin/ls
$ which bash
/bin/bash
$ which less
/usr/bin/less
$ which cd
$ type cd
cd is a shell builtin
```

A final thing to note is that the Conda system installed everything within the home directory, and thus did not require administrator level access. On these VMs you are the only user, but many Linux systems are shared and the ability to install your own software without affecting other users or needing the administrator password can be very useful.

#### 1.13 Useful Resources

**Directory of Linux commands** (http://archive.oreilly.com/linux/cmd/):

A directory of 687 Linux commands (taken from Linux in a Nutshell, 5th Edition), with for each command a description and list of available options.

**LinuxQuestions** (http://www.linuxquestions.org):

A community-driven, self-help web site for Linux users. The most popular section of the site are the forums, where Linux users can share their knowledge and experience. Newcomers to the Linux world (often called newbies) can ask questions and Linux experts can offer advice.

**Stack Overflow** (http://stackoverflow.com):

A question-and-answer website on the topic of computer programming.

**Biostars** (http://www.biostars.org):

A question-and-answer website with a focus on bioinformatics, computational genomics and biological data analysis.

*Before posting a question on one of the above websites, you should always first try to find the answer yourself by (1) doing a Google search and (2) searching the website for previously asked questions!*

Also, even in this internet age, it's worth getting a good book.

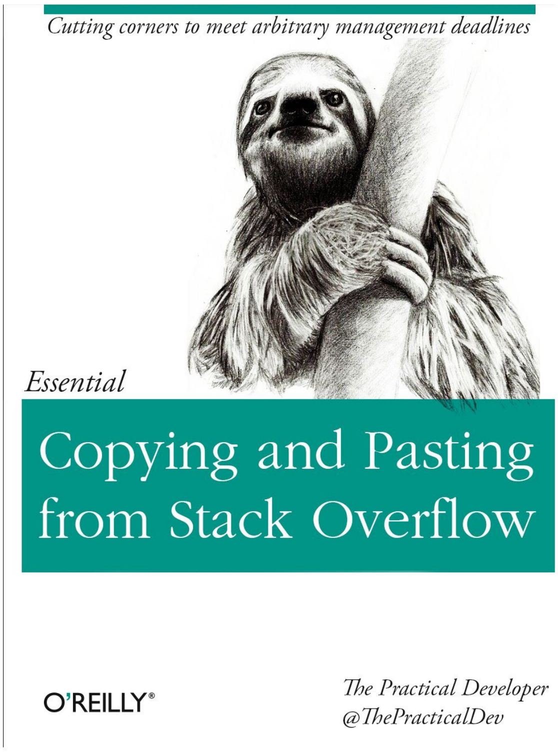

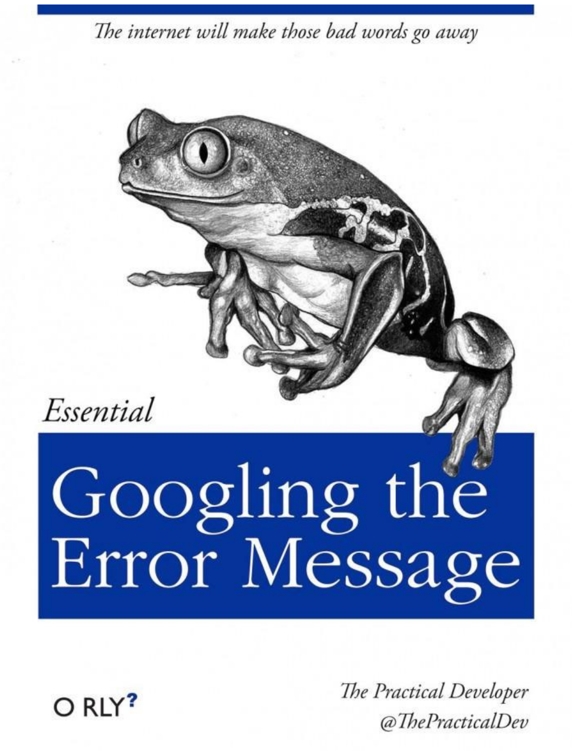

### Dry Lab: Day 1

### $ raw_data | quality

**Instructors:**

Tim Booth - Bioinformatician and Programmer

Nathan Medd - Training and Outreach Manager

### 2. Quality control and data preprocessing

During this section you will learn to:

- Assess the intrinsic quality of your raw reads (in fastq format) using metrics generated by the sequencing platform (e.g. quality scores)
- Pre-process data, i.e. trimming the low-quality bases, remove short reads.

#### 2.1 Software

You will be using the software we installed earlier from NanoPack; NanoComp, chopper, NanoPlot for this task. It is a set of Python scripts for visualising and processing long-read sequencing data (both PacBio and ONT). NanoPack and individual scripts are available through the public software repositories using pip and bioconda through conda (https://github.com/wdecoster/nanopack).

NanoPack can use multiple formats of raw data e.g. Fast5, FASTQ, ONT sequencing summary files, BAM (both mapped and unmapped), etc. But in these sections, we will focus on the base-called data in FASTQ format sequenced on the ONT platform.

#### 2.2 Data

First we will have to find our data on the system. This has been saved for you in our shared drive here:

/mnt/s3fs/NERC_training/june_2023/prom_data/raw_data/

Take a look:

```
$ ls /mnt/s3fs/NERC_training/june_2023/prom_data/raw_data/
```

You should see a collection of files, including one that has been labelled with your sample number, this is your raw data from the sequencer that has been uploaded this week:

<YOUR_FASTQ_FILE>.fastq.gz

Let's move to the directory "QC" and keep tidy any files we produce in this section:

```
$ cd ~/NERC_training/QC
```

We'll make a 'soft link' to your data in your home directory to save copying across a large data file:

```
$ ln -sr
/mnt/s3fs/NERC_training/june_2023/prom_data/raw_data/<YOUR_FASTQ_FILE>.fastq.gz
.
```

#### 2.3 Addressing QC with NanoPlot

We will use NanoPlot from NanoPack to assess the quality of your FASTQ file.

Help message can be displayed by:

```
$ NanoPlot -h
```

Once you've had a look at the options for running NanoPlot we'll go ahead and run it using our data in a fairly default setting (this may take around 15 mins):

```
$ NanoPlot -t 2 --fastq <YOUR_FASTQ_FILE>.fastq.gz --loglength -
o <YOUR_FILE>_nanoplot --plots dot
```

where:

-t Set the number of threads to be used by the script
--fastq Data is in one or more default fastq file
--loglength Show logarithmic scaling of lengths in plots
--plots Specify which bivariate plots have to be made (kde, hex,dot,pauvre)

Let's take a look at the output:

```
$ cd <YOUR_FILE>_nanoplot

$ ls
```

As you can see, NanoPlot has created a few images and a stats file in text format. All of these are summarized in a nice html report. Let's take a look at that:

```
$ google-chrome NanoPlot-report.html &
```

The first thing you'll see is a table containing some useful statistics about your data:

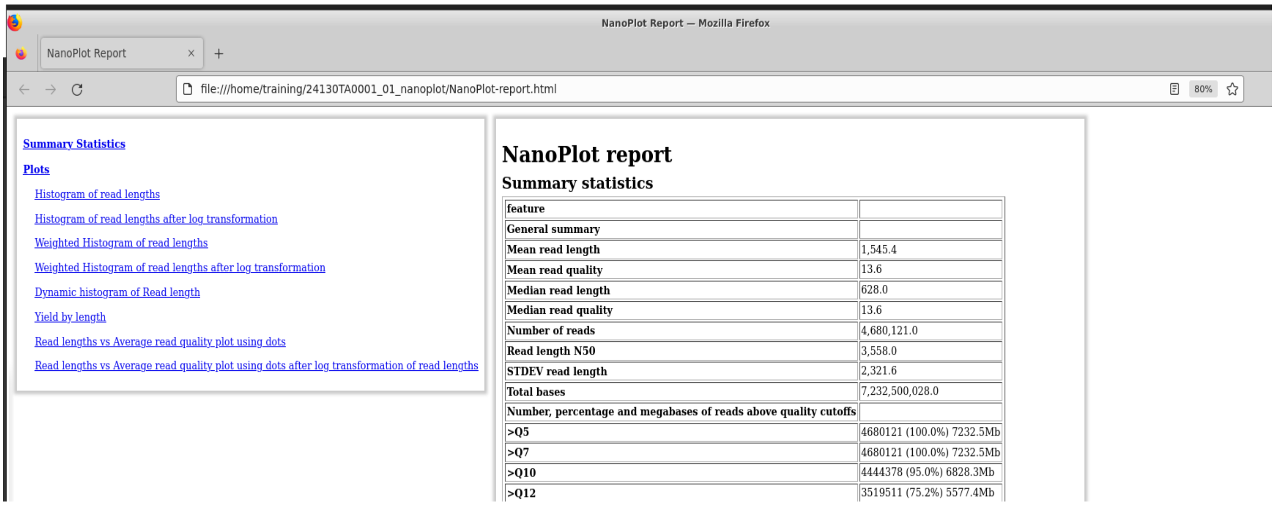

NanoPlot report

Summary statistics

| feature | |
|---|---|
| General summary | |
| Mean read length | 1,545.4 |
| Mean read quality | 13.6 |
| Median read length | 628.0 |
| Median read quality | 13.6 |
| Number of reads | 4,680,121.0 |
| Read length N50 | 3,558.0 |
| STDEV read length | 2,321.6 |
| Total bases | 7,232,500,028.0 |
| Number, percentage and megabases of reads above quality cutoffs | |
| >Q5 | 4680121 (100.0%) 7232.5Mb |
| >Q7 | 4680121 (100.0%) 7232.5Mb |
| >Q10 | 4444378 (95.0%) 6828.3Mb |
| >Q12 | 3519511 (75.2%) 5577.4Mb |

***Questions:***

1. ***How many reads are in the sample?***

2. ***What is the mean quality?***

3. ***What's the read length N50?***

4. ***Overall, how good is your data?***

**Add the answer to these questions to this spreadsheet:** [https://tinyurl.com/4hsufvn7](https://tinyurl.com/4hsufvn7)

#### 2.4 Filter reads with Chopper

We will filter the reads based on your average read quality generated above, <your_mean_qual>, and on a minimum read length using Chopper. This tool will also remove reads mapping to the lambda phage genome (control DNA used in nanopore sequencing).

We're going to have to pipe a couple of commands into each other here because Chopper needs an unzipped version of our raw data file, we then write out to a new '_trimmed.fastq.gzip' file using a redirect '>'. This should take 15-20 mins.

```
$ gunzip -c <YOUR_FASTQ_FILE>.fastq.gz | chopper -q <your_mean_qual> -l 500 | gzip >
<YOUR_FASTQ_FILE>_trimmed.fastq.gz
```

We will use NanoComp to compare the data pre and post filtering. NanoComp can also be used to compare the quality of multiple samples/runs in one go.

Unlike Illumina short reads, you may not see a massive difference in the quality improvement of the data but the aim here is to check if the filtering cutoffs were not too stringent and also if there is enough data left to carry out further analysis. This step should take ~20mins.

```
$ NanoComp --fastq <YOUR_FASTQ_FILE>.fastq.gz
<YOUR_FASTQ_FILE>_trimmed.fastq.gz -t 2 --names reads_raw reads_trimmed -o
<YOUR_FILE>_Nanocomp
```

Let's take a look at the results:

```
$ cd <YOUR_FILE>_Nanocomp

$ google-chrome NanoComp-report.html &
```

***Questions***:

1. ***Spot the differences! Fill out the same metrics from the previous question and compare results.***
2. ***How big is the loss in the yield?***
3. ***What is our average quality now?***

**Add the answer to these questions to this spreadsheet:** https://tinyurl.com/4hsufvn7

### Dry Lab: Day 2

### $ assembly | annotation

**Instructors:**

- Urmi Trivedi, Bioinformatics Team Leader, Edinburgh Genomics
- Heleen De Weerd, Bioinformatics analyst, Edinburgh Genomics

### 3. *De novo* nuclear genome assembly

During this session you will learn to:

- Generate a *de novo* genome assembly using Redbean (wtdbg2)
- Assess the quality of assembly

#### 3.1 Software

All these software are open source and can be downloaded from the links given below:

- Redbean (wtdbg2) https://github.com/ruanjue/wtdbg2
- QUAST: http://quast.sourceforge.net/
- Minimap2: https://github.com/lh3/minimap2
- Samtools: http://samtools.github.io/

#### 3.2 Data

We will be using the FASTQ data for the sample sequenced by you. We have filtered lambda phage reads from our data and also trimmed the remaining data using NanoFilt with length cutoff of 500 and average quality cutoff of 9.

```
$ cd /home/training/NERC_training/nuclear_assembly/

$ ls
```

Create a softlink to your trimmed file within this directory.

```
$ ln -s ../QC/<YOUR_FASTQ_FILE>
```

#### 3.3 Generate *de novo* genome assembly

Redbean is one of the most widely used tools for assembling genomes *de novo* from long noisy reads produced by PacBio or Oxford Nanopore Technologies (ONT). It assembles raw reads without error correction and then builds the consensus from intermediate assembly output. It is one of the fastest assemblers available and is designed for a wide range of datasets from small bacteria to large mammals.

wtdbg2 has two key components: an assembler wtdbg2 and a consenser wtpoa-cns.

##### 3.3.1. Wtdbg2

wtdbg2 assembles raw reads and generates the contig layout and edge sequences in a file "*prefix*.ctg.lay.gz". In order to run this command, you need to use the trimmed fastq file you generated yesterday. Furthermore, one of the mandatory parameters the program requires is estimated genome size. You can select this from the table below based on the barcode number you used during the course. These genome size estimates are for the species we are analysing here (but not the same individuals) based on flow cytometry, a non-sequenced based method. These data were downloaded from the Kew Plant DNA C-value database (https://cvalues.science.kew.org/)  Note that these are haploid values,  which is the standard way to report genome sizes.

| **Barcode number** | **Expected Genome size** |
|---|---|
| 1 | 1.45 GB |
| 2 | 1.45 GB |
| 3 | 1.1 GB |
| 4 | 1.1 GB |
| 5 | 600 MB |
| 6 | 1.4 GB |
| 7 | 300 MB |
| 8 | 1.5 GB |
| 9 | 1.5 GB |
| 10 | 1.5 GB |
| 11 | 1.5 GB |

| | |
|---|---|
| 12 | 900 MB |
| 13 | 600 MB |
| 14 | 1.5 GB |

Run wtdbg2:

```
$ wtdbg2 -i <YOUR_FASTQ_FILE> -o wtdbg2 -t 36 -x ont -g <estimated_genome_size>
```

where,

-i Long reads sequences file (can be multiple)
-o Prefix of output files
-t Number of threads
-x ONT preset (sets other params like k-mer sizes,etc. accordingly)
-g Estimated genome size (We need to select this value from the table shown above. This should correspond to your barcode ID and can be specified as: 1.4g, 800m, etc.)

```
training@vm-training_updateForNERC:nuclear_assembly$ wtdbg2 -i P_lancelota.fastq.gz -o wtdbg2 -t 36 -x ont -g 1.4g
--
-- total memory        3965164.0 kB
-- available           3483748.0 kB
-- 2 cores
-- Starting program: wtdbg2 -i P_lancelota.fastq.gz -o wtdbg2 -t 36 -x ont -g 1.4g
-- pid                     2996
-- date         Tue Apr 11 11:11:34 2023
--
[Tue Apr 11 11:11:34 2023] loading reads
40000
```

Check out the kmer-distribution:

```
********************** Kmer Frequency **********************
|
|
|
|
|
|
|
||
||
||
||
||
|||
|||
|||
||||
||||
|||||
|||||||
||||||||||||||||||||||||||||||||||||||||||||||||||||||||||||||||||||||||||||||||||||||||||||||||||||
**********************     1 - 201    **********************
Quatiles:
   10%   20%   30%   40%   50%   60%   70%   80%   90%   95%
     2     3     5     8    17    49   144   775  2782 10603
** PROC_STAT(0) **: real 148.634 sec, user 521.410 sec, sys 57.180 sec, maxrss 4595492.0 kB, maxvsize 7456560.0 kB
[Tue Apr 11 11:24:23 2023] - high frequency kmer depth is set to 11644
[Tue Apr 11 11:24:23 2023] - Total kmers = 167005956
[Tue Apr 11 11:24:23 2023] - average kmer depth = 8
[Tue Apr 11 11:24:23 2023] - 77742086 low frequency kmers (<2)
[Tue Apr 11 11:24:23 2023] - 2091 high frequency kmers (>11644)
[illegible]
```

This plot shows that most Kmers are composed of singletons 77M/167M showing the genome is extremely repetitive and contains erroneous reads. Also, the average kmer depth is extremely low (8) which shows the genome coverage is sparse. For an ideal assembly most k-mers are usually within 2~1000 and average k-mer depth is between 20 - 100.

***How does this distribution look for you?***

##### 3.3.1. wtpoa-cns

wtpoa-cns takes "*prefix*.ctg.lay.gz" as input and produces the final consensus in FASTA.

```
$ wtpoa-cns -t 32 -i wtdbg2.ctg.lay.gz -fo wtdbg2.ctg.fa
```

#### 3.5 Polishing genome assemblies

We will be using mapping based polishing methods to polish the nuclear genome assembly we have generated in the previous steps.

##### 3.5.1 Map the reads to assembly

We will use minimap2 for mapping the raw data to the Redbean assembly generated earlier. Minimap2 is a versatile sequence alignment program that aligns DNA or mRNA sequences against a large reference database. Minimap2 is tens of times faster than mainstream long-read mappers such as BLASR, BWA-MEM, NGMLR and GMAP (https://github.com/lh3/minimap2). It is compatible with the long reads produced from both PacBio and ONT.

```
$ minimap2 -ax map-ont -t 36 wtdbg2.ctg.fa <YOUR_FASTQ_FILE> | samtools sort -@36 >
```

```
minimap2_wtdbg2_sorted.bam
```

##### 3.5.1 Get mapping stats

We will explore samtools (https://github.com/samtools/samtools) to get the mapping stats and perform other operations such as sorting and indexing the BAM file.

```
$ samtools index minimap2_wtdbg2_sorted.bam

$ samtools flagstat minimap2_wtdbg2_sorted.bam > minimap2_wtdbg2_sorted.bam.stat

$ cat minimap2_wtdbg2_sorted.bam.stat
```

```
training@vm-training_updateForNERC:nuclear_assembly$ cat minimap2_wtdbg2_sorted.bam.stat
29710082 + 0 in total (QC-passed reads + QC-failed reads)
4680121 + 0 primary
19609392 + 0 secondary
5420569 + 0 supplementary
0 + 0 duplicates
0 + 0 primary duplicates
29573477 + 0 mapped (99.54% : N/A)
4543516 + 0 primary mapped (97.08% : N/A)
0 + 0 paired in sequencing
0 + 0 read1
0 + 0 read2
0 + 0 properly paired (N/A : N/A)
0 + 0 with itself and mate mapped
0 + 0 singletons (N/A : N/A)
0 + 0 with mate mapped to a different chr
0 + 0 with mate mapped to a different chr (mapQ>=5)
```

A read may map ambiguously to multiple locations, e.g. due to repeats. Only one of the multiple read alignments is considered primary, and this decision may be arbitrary. All other alignments have the secondary alignment flag.

##### 3.5.2 Polish

We are only selecting primary alignments and excluding supplementary/secondary alignments. This is done to get more confidence which will be the main basis for polishing/error correction. Later, the consensus is called again based on the mapping data.

```
$ samtools view -F0x772 minimap2_wtdbg2_sorted.bam | wtpoa-cns -t 36 -d wtdbg2.ctg.fa -
i - -fo wtdbg2.cns.fa
```

### 3.6 Assessment of assemblies

#### 3.6.1 Using QUAST

QUAST is a quality assessment tool for evaluating and comparing genome assemblies. QUAST can evaluate assemblies with a reference genome as well as without a reference.

Help message can be accessed by:

```
$ quast --help
```

Run QUAST:

```
$ quast -o quast -t 36 --labels raw,polished wtdbg2.ctg.fa wtdbg2.cns.fa
```

QUAST produces many reports, summary tables and plots for exploration of assembly quality. The key start point is the “report.html” file where all of these are summarized.

```
$ cd quast

$ google-chrome report.html &
```

View in Icarus contig browser

All statistics are based on contigs of size >= 500 bp, unless otherwise noted (e.g., "# contigs (>= 0 bp)" and "Total length (>= 0 bp)" include all contigs).

Worst Median Best — Show heatmap

| Statistics without reference | raw | polished |
|---|---|---|
| # contigs | 7072 | 7052 |
| # contigs (>= 0 bp) | 7072 | 7072 |
| # contigs (>= 1000 bp) | 7072 | 6997 |
| # contigs (>= 5000 bp) | 6351 | 4619 |
| # contigs (>= 10000 bp) | 2919 | 1839 |
| # contigs (>= 25000 bp) | 424 | 211 |
| # contigs (>= 50000 bp) | 37 | 24 |
| Largest contig | 131 313 | 181 451 |
| Total length | 80 504 411 | 59 160 116 |
| Total length (>= 0 bp) | 80 504 411 | 59 165 881 |
| Total length (>= 1000 bp) | 80 504 411 | 59 118 384 |
| Total length (>= 5000 bp) | 77 121 263 | 50 665 185 |
| Total length (>= 10000 bp) | 52 759 139 | 31 170 925 |
| Total length (>= 25000 bp) | 15 001 619 | 7 592 426 |
| Total length (>= 50000 bp) | 2 500 222 | 1 734 393 |
| N50 | 13 498 | 10 558 |
| N90 | 5854 | 4509 |
| auN | 17 418 | 14 716 |
| L50 | 1843 | 1685 |
| L90 | 5498 | 5160 |
| GC (%) | 37.05 | 37.13 |
| **Mismatches** | | |
| # N's per 100 kbp | 0 | 0 |
| # N's | 0 | 0 |

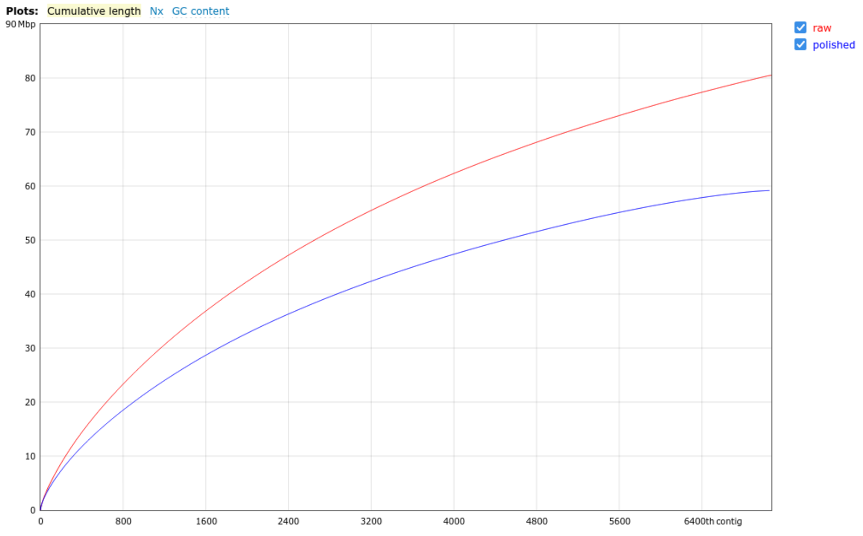

Contigs are ordered from largest (contig #1) to smallest.

***Questions:***

1. ***Is there any difference in the stats between these assemblies (E.g. higher N50 and less number of contigs) ?***

2. ***We have pregenerated the assemblies for the data across the class. Let's see which species/data have given us the best assembly! You can check the quast report for these -***

   ***Raw assemblies***

```
google-chrome
/mnt/s3fs/NERC_training/june_2023/prom_data/quast_nuclear_raw/report.html &
```

   Polished assemblies:

```
google-chrome
/mnt/s3fs/NERC_training/june_2023/prom_data/quast_nuclear_polished/report.html
&
```

### 4. Plastid assembly

During this session you will learn to:

- Generate a *plastome* assembly using ptGAUL
- Visualize plastid assembly

#### 4.1 ptGAUL:

You will be using ptGAUL for this task. ptGAUL (***p**las**t**id **G**enome **A**ssembly **U**sing **L**ong-read data) is a plastome assembly pipeline for long read data like Oxford Nanopore and PacBio.

The ptGAUL pipeline includes three major parts: filtering long reads, setting the depth of coverage, and assembling the filtered plastid data.

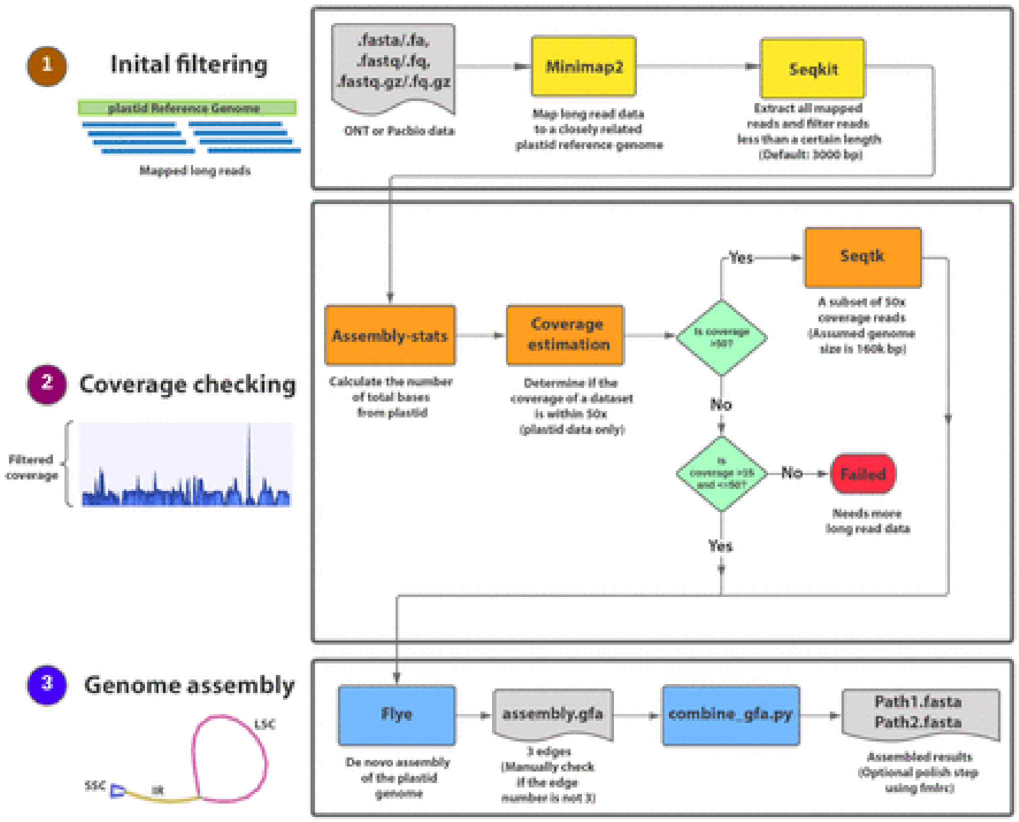

Along with the fastq file, other data that ptGAUL will require is the database. For this purpose, we have selected one thousand random sequences from Refseq plastidDB available at -

https://ftp.ncbi.nlm.nih.gov/refseq/release/plastid/. You can find the downsampled version of the database at -

```
$ cd /home/training/NERC_training/plastid_assembly/

$ ls plastid_DB
```

```
training@vm-training_updateForNERC:plastid_assembly$ ls plastid_DB/
plastid_db_1K.fna  plastid_db_1K.fna.fai
```

##### 4.1.1. Run ptGAUL

```
$ ptGAUL.sh -l <FASTQ_FILE_PATH> -t 36 -o ptgaul -r plastid_DB/plastid_db_1K.fna
```

where:

-l Input fastq file
-r Plastid database
-t Number of threads

Don't panic if you get:

```
==============================================
-------- Oops! Detect a weird result ----------
==============================================
```

Plastome can be linear, branched, or occasionally circular molecules. The circular molecules contain a large, inverted repeat (IR) and large and small single-copy regions (LSC and SSC). For such plastomes, they usually can have IR regions in different directions. Considering this, if flye produces between 1-3 contigs, ptGAUL attempts to combine these contigs and create a circular genotype with IR given in a different direction. If flye generates more than 3 contigs, the pipeline suggests manual checks before combining.

Check the output:

```
$ cd ptgaul/result_3000
```

```
training@vm-training_updateForNERC:result_3000$ ls
edges.fa      filter_name              new_filter_gt3000.fa
filter.paf    filter_reads             ptGAUL_final_assembly
filter_1.paf  filter_reads_final.fa    sorted_depth
filter_2.paf  flye_cpONT               total_length_of_plastid_reads
```

```
$ grep -c ">" flye_cpONT/assembly.fasta
```

```
$ quast -o quast_flye -t 36 flye_cpONT/assembly.fasta

$ cd quast_flye

$ google-chrome report.html
```

Has anyone got more than 3 contigs?

*We have pregenerated the plastid assemblies for the data across the class. Let's see which species/data have given us the best assembly! You can check the quast report for these -*

```
google-chrome
/mnt/s3fs/NERC_training/june_2023/prom_data/quast_ptgaul_flye/report.html
```

Let's see if ptGAUL has managed to stitch these contigs and generate a circular plastome

```
$ cd ../ptGAUL_final_assembly/

$ ls
```

Let's get assembly stats for these paths.

```
$ quast -o quast_ptgaul -t 36 –-labels path1,path2 path1.fasta path2.fasta

$ cd quast_ptgaul

$ google-chrome report.html
```

Usually, the stats for both paths are the same. ***What does yours look like?***

### Dry Lab: Day 3

### $ identity | phylogeny

**Instructors:**

- Urmi Trivedi, Bioinformatics Team Leader, Edinburgh Genomics
- Heleen De Weerd, Bioinformatics analyst, Edinburgh Genomics

### 5. Sequence Annotation

#### 5.1 PGA

We have an assembled plastid sequence, the next step for us to do is to annotate the sequence. This can be done in different ways, in our case we will use Plastid Genome Annotator (PGA). PGA is a tool specifically made for plastid annotation, however many different annotation tools are available.

PGA takes a number of genbank files as reference. In our case these files contain information on the plastid genome of a specific species as well as the genes which are present. It will also need the fasta files of the sequences we want to annotate, we will annotate both created paths.

We have downloaded the genbank files for different related species already.

```
$ cd /home/training/NERC_training/plastid_annotation/
```

We need to make a folder for our created sequence and link them to this folder.

```
$ cp -R /mnt/s3fs/NERC_training/reference/plastid_annotation_references/ ./

$ mkdir paths

$ ln -s
/home/training/NERC_training/plastid_assembly/ptgaul/result_3000/ptGAUL_final_assembl
y/*.fasta ./paths/

$ ls *
```

Once we have the reference ready and we have our sequence we can start the annotation.

```
$ perl /mnt/autosquash/PGA/PGA.pl -r
/home/training/NERC_training/plastid_annotation/plastid_annotation_references/ -t paths/ -
out annotated/
```

where,

-r is the folder containing reference genbank files
-t is the folder with the fasta sequences to be annotated
-out is the folder to which the results will be written

Once the tool is finished we can check the output folder, we should see our two sequences now are genbank files.

```
$ ls annotated/
```

We can look at these annotations using UGENE. This tool can be used to do different types of visualisation, comparison and many other things. If you have time, take a look at the functionalities of this tool. To start UGENE run the following command:

```
$ ugeneui
```

In UGENE, open the created annotation by clicking File → Open. Find your annotated file and open it. The image you see should look something like the following.

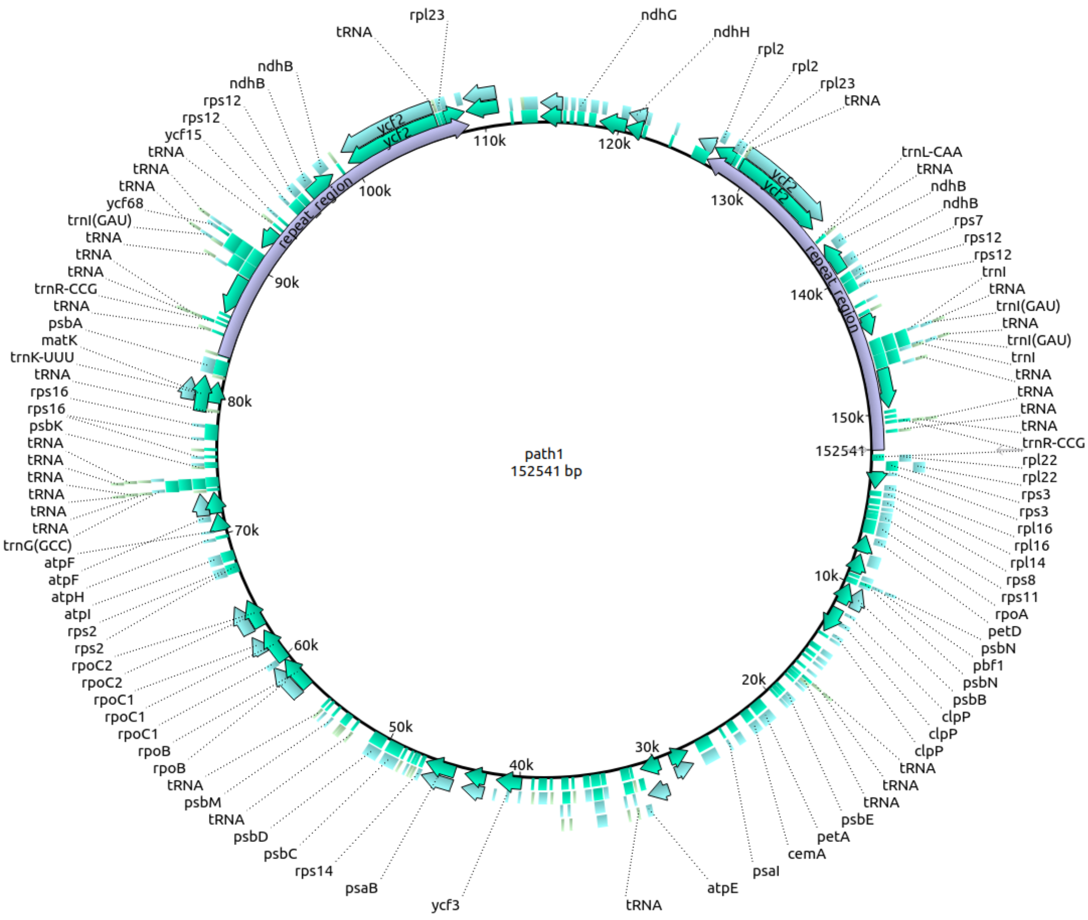

***Open both created paths, can you find differences between these two paths looking at the annotations?***

UGENE also has the option to create comparisons between two sequences using dotplots. For the creation of a dotplot you need two sequences, these will be compared to each other. In essence a dotplot will show you any position on the first genome which is similar to a position anywhere on the other genome as a dot, when positions do not match the graph will not show you anything.

Using the two different paths, create a dotplot by clicking Tools → Build dotplot. Select path1 and path2 as the two different sequences, make sure to use genbank files so you will also see the annotations for the sequences. Click next, check "Search inverted repeat" and click "OK". You image should look something like this:

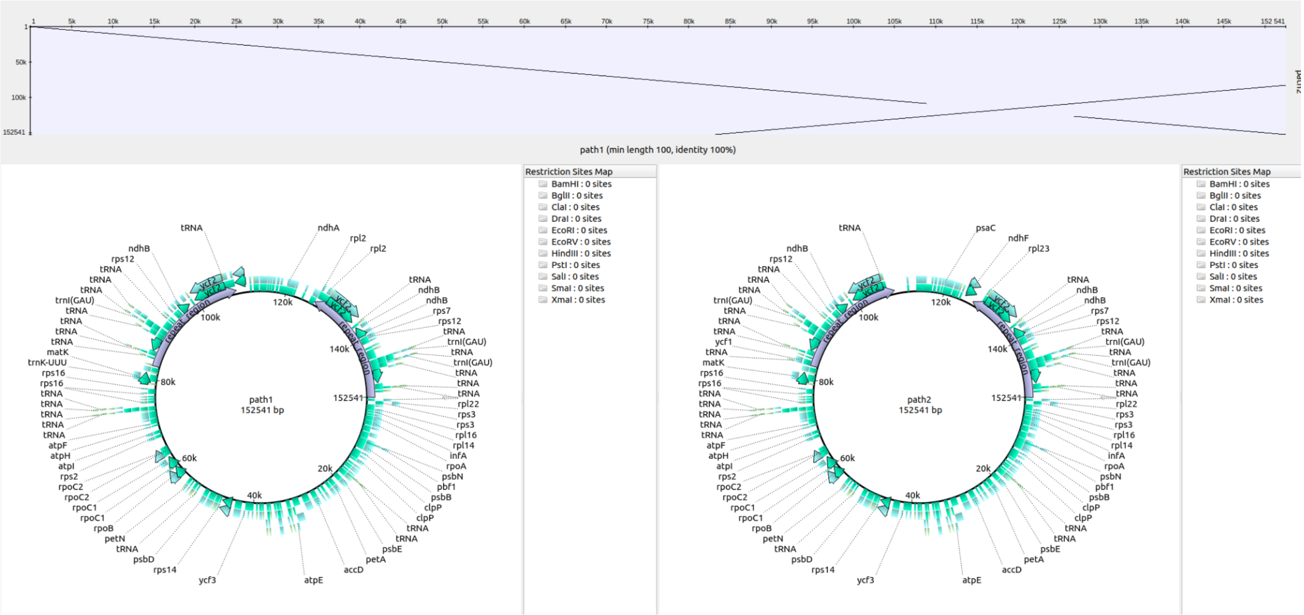

***Can you see differences between the two paths? If so, what are the differences?***

We want to see if our paths will have a similar pattern and order to previously published data. We have downloaded a reference to be able to compare our sequence to this reference.

Once again go to Tools → Build dotplot. Select one of your sequences and the reference sequence (located at /home/training/NERC_training/plastid_annotation/plastid_annotation_references/Cotinus_coggygria_NC_054342.1.gb). Make sure to again check "Search inverted repeat" and in this case also set the identity to 90%.

***Are there differences between your sequence and the reference sequence?***

***Check both paths in this manner, does one seem more similar to the reference? Make a note of this, you will need it later on.***

Do not worry if your sequences are different to the reference sequence, we can continue with these sequences.

In the case that the sequence has a different order, in a regular situation we would try to find out why the sequence is different, try different settings for the assembly of the plastid, try different tools

to see if another tool might be more suitable for this specific species or run, however in this case we do not have the time.
If you do have the time and want to take a look, you can rerun ptGAUL using different settings, use -c in ptGAUL to change the minimal coverage. You do not have to redo the annotation for the creation of the dotplot, you can create a dotplot between a fasta file and a genbank in UGENE without a problem.

In some cases, ptGAUL has made assumptions which are not true. We will take a look together to see if we can fix some of these assumptions.

#### 5.2 BOLD & BLAST

Now that we have a sequence and we have an annotation, we need to finally find out what we are actually working on. For this we will extract a barcode gene from one of the paths, which we will run through BOLD to get an idea of what we are looking at.

The barcode gene we will take a look at is matK. Open your genbank file in UGENE by going to File → Open and locating your genbank file.  We can look for the gene visually, however searching for it is an easier option. At the bottom of your screen below the visual version of our plastid, there is a list with feature information which can be opened. Right click the list and choose Find Qualifier. In the pop-up (shown in the figure below), in the field for value, we are going to put the name of the gene which we are looking for.

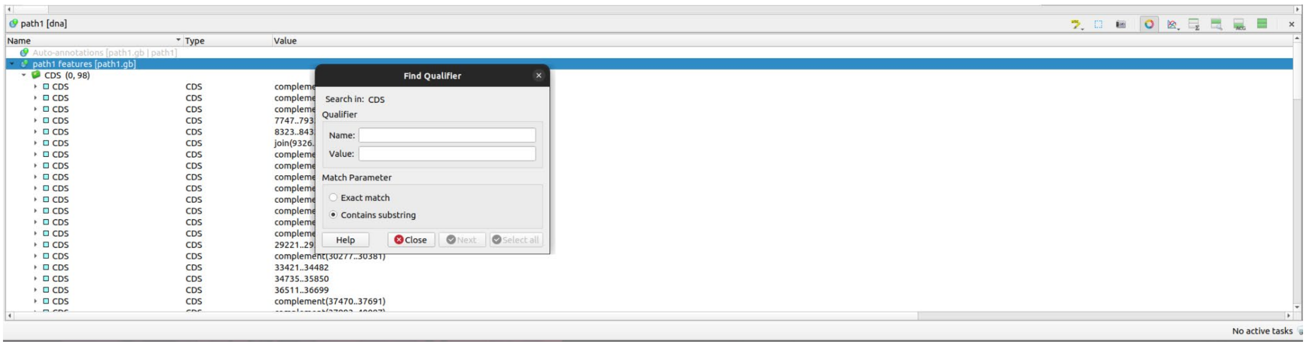

In this case we will look for matK. Click next and we should find the gene of interest. Once we have found it, we can click on the gene and see where it is on our plasmid. To copy the sequence, we can right click the gene in the feature list, go to copy/paste and select “Copy annotation sequence”. We can now paste the sequence anywhere, you can save the sequence in a separate file if you would like, however in this case we can paste the sequence into BOLD directly.

Go to https://www.boldsystems.org/index.php/IDS_OpenIdEngine, select “Plant Identification” and paste the sequence into the fasta sequence field. Continue by pressing Submit at the bottom of the form. It might take a few minutes before you get any results. The results should look something like the following.

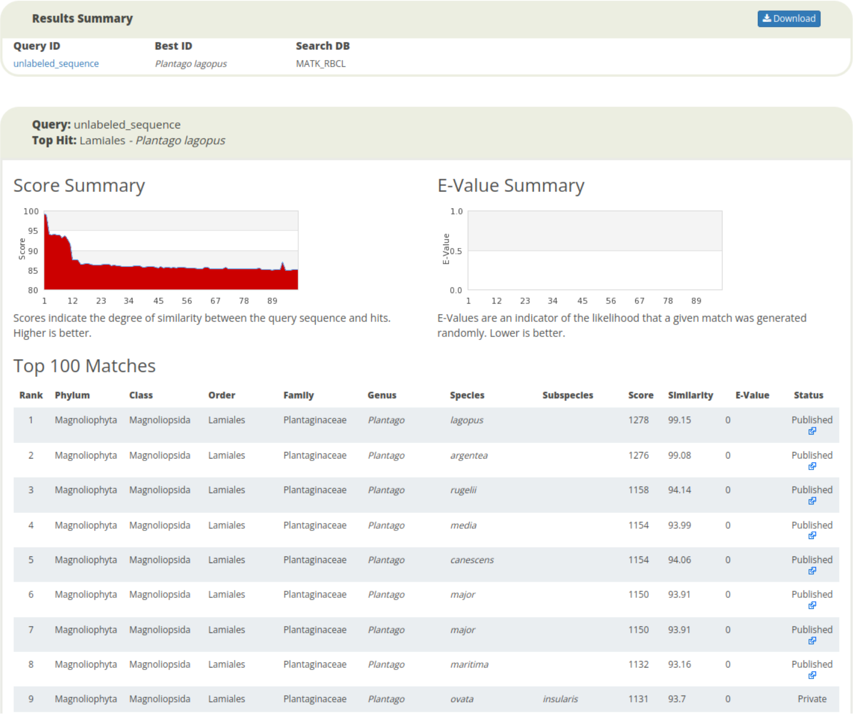

As a second identification method BLAST can be used (https://blast.ncbi.nlm.nih.gov/Blast.cgi). Click on nucleotide blast and paste the sequence into the top box. In this case we do not need to change any of the other settings, so we can click BLAST at the bottom to start the comparison.

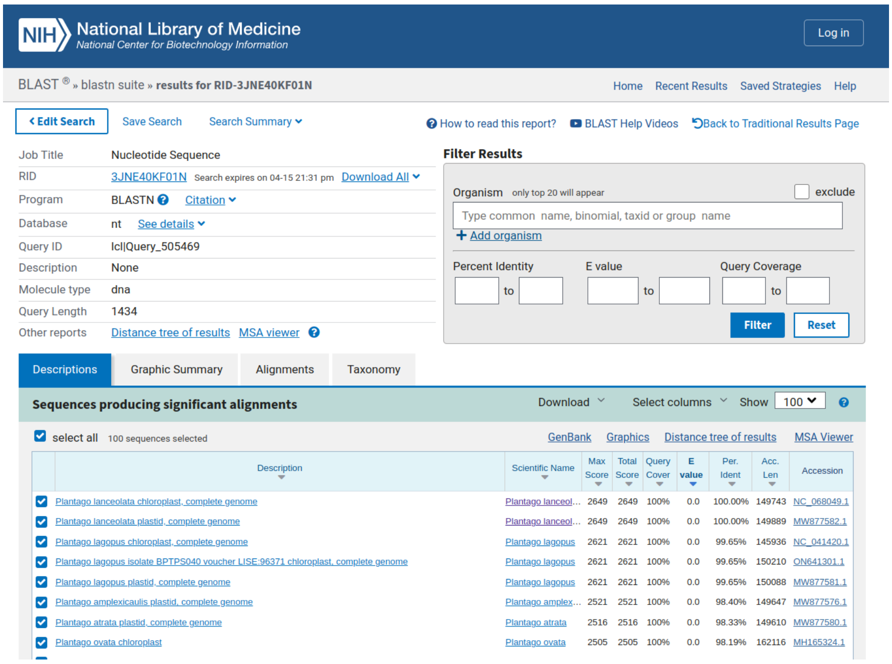

If you have time, do the same for the rbcL gene, do the results match?

**What species do you think you are working on? Add your prediction to this spreadsheet: https://tinyurl.com/4hsufvn7**

### 6. Phylogenetic Tree

We know what we are working with, but how does our sequence compare to others? For this we are going to make a phylogenetic tree, using different methods. We will use a multiple sequence alignment and a k-mer based approach. We will take a look at IQ-Tree as well.

```
$ cd /home/training/NERC_training/phylogenetics
```

#### 6.1 Multiple Sequence Alignment

We will start with the multiple sequence alignment. We have taken sequences from the public domain and created an alignment to save time. We can take a look at this alignment without adding our sequences first.

To visualise the multi sequence we can open the file using UGENE. At the bottom of the visualisation you will see an overview of how many sequences have a nucleotide per position. This means we can see highly conserved regions and regions which are more specific to a group or family.

Open the alignment in UGENE:
/home/training/NERC_training/NERC_training/demo_data/multiple_sequence_alignment

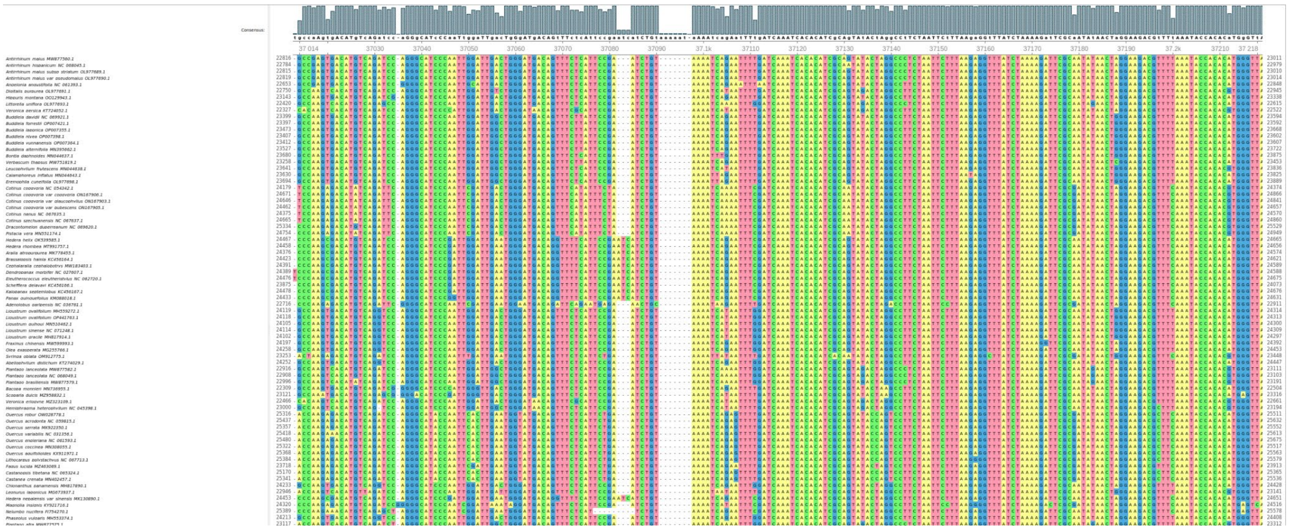

You will see a list of sequences annotated with their species name, at the bottom of the screen you will see how many sequences have a base per position.

***Are there regions which are only present in a few species?***

If needed you can zoom out by right clicking → Appearance → Zoom Out

For our comparison, we will only look at sequences which are within the family. For each family a file is available at /home/training/NERC_training/phylogenetics/family_alignments

You can take a look at the alignment file once again, however now we will add our own sequence to this. To be able to do this, we need to use the path which is most similar to the reference sequence. We took a look at this before, if you do not remember, take a look at the end of day2 of bioinformatics.
For the multiple sequence alignment we will use mafft.

```
$ mafft --add <YOUR_PATH_FILE> <FAMILY.aln> > FamilyPlusPath.aln
```

Once we have added our sequence, open up your alignment file and take a look using UGENE. Everyone's alignment will look different at this point.

We can make this multiple sequence alignment into a tree using UGENE. Right click the alignment, go to Tree → Build Tree. Different methods for tree building are available. For now we will use standard settings. Click Build.

***Does your sequence cluster close to the species we found using BOLD?***

**Have you changed your mind on the species you are working on? Add your prediction to this spreadsheet:** https://tinyurl.com/4hsufvn7

#### 6.2 Kmer based approach

If your sequence might not have had the same orientation as the reference file, your tree might have not looked completely right. Multiple sequence alignments assume that your sequences are in the same order and have the same structure. This could cause issues, however there is a solution for that. We can use a kmer based approach.

```
$ cp -R
/mnt/s3fs/NERC_training/reference/phylogenetics/kmer_references/ ./kmer_references_seq
uences

$ ls kmer_references_sequences
```

In this folder there should be a file named ListOfSequences.txt. This file contains a list of all the files in the folder. If we want to add our sequence to this list, we need to copy the fasta(!) files for one of both of our paths into this folder and add the name of the file to the ListOfSequences.txt file. Use what you have learned about Linux to do this on the command line, or use the graphical interface that this VM has to do this.

```
$ cp
/home/training/NERC_training/plastid_assembly/ptgaul/result_3000/ptGAUL_final_assembly/*.fasta ./kmer_references/
```

Once you have added your paths to the folder and the list, run the following command to create a kmer based tree:

```
$ SANS -f strict -i kmer_references/ListOfSequences.txt -N kmertree.nwk
```

where,

-f filter, this needs to be strict if we want a tree
-i is the input file with a list of one genome per line
-N Output Newick file (the tree file)

We can open this tree using UGENE or the interactive tree of life (https://itol.embl.de/).

***Which species is your sequence closest to?***

***Is this in line with what you found using BOLD?***

**Have you changed your mind on the species you are working on? Add your prediction to this spreadsheet:** https://tinyurl.com/4hsufvn7

#### 6.3 IQ-TREE

For the creation of trees, many different models could be used. These models have assumptions, assumptions on the type of data you are working with and the relationships between sequences. When working with our data, we want the best fit, but not spend many hours going through different models and evaluating every model, seeing which would fit best. IQ-TREE tests a large number of different models on a multiple sequence alignment to determine which model would work best for the dataset. In this case we are working with DNA sequences, which means that IQ-TREE will test 286 different models, and tell us which one fits best with our data.

To run IQ-tree there are two options, using the web version: http://iqtree.cibiv.univie.ac.at/ or the command line version. It is recommended to use the command line version, as the online version has limitations.

```
$ iqtree --help

$ iqtree -s FamilyPlusPath.aln -alrt 1000 -bb 1000
```

where,
-s Input alignment
-alrt Do the parametric aLRT test x times
-bb Do ultrafast bootstrapping x times

IQ-TREE will report the different models it is testing and the results of the test. The screen should look like the image below.

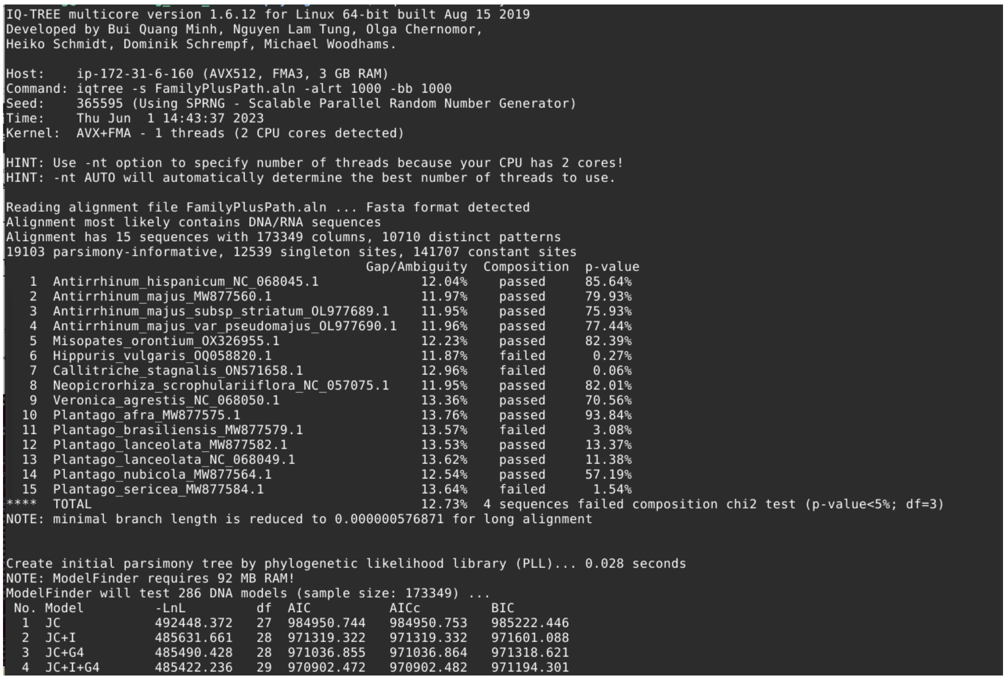

```
IQ-TREE multicore version 1.6.12 for Linux 64-bit built Aug 15 2019
Developed by Bui Quang Minh, Nguyen Lam Tung, Olga Chernomor,
Heiko Schmidt, Dominik Schrempf, Michael Woodhams.

Host:    ip-172-31-6-160 (AVX512, FMA3, 3 GB RAM)
Command: iqtree -s FamilyPlusPath.aln -alrt 1000 -bb 1000
Seed:    365595 (Using SPRNG - Scalable Parallel Random Number Generator)
Time:    Thu Jun  1 14:43:37 2023
Kernel:  AVX+FMA - 1 threads (2 CPU cores detected)

HINT: Use -nt option to specify number of threads because your CPU has 2 cores!
HINT: -nt AUTO will automatically determine the best number of threads to use.

Reading alignment file FamilyPlusPath.aln ... Fasta format detected
Alignment most likely contains DNA/RNA sequences
Alignment has 15 sequences with 173349 columns, 10710 distinct patterns
19103 parsimony-informative, 12539 singleton sites, 141707 constant sites
                                           Gap/Ambiguity  Composition  p-value
   1  Antirrhinum_hispanicum_NC_068045.1            12.04%    passed     85.64%
   2  Antirrhinum_majus_MW877560.1                  11.97%    passed     79.93%
   3  Antirrhinum_majus_subsp_striatum_OL977689.1   11.95%    passed     75.93%
   4  Antirrhinum_majus_var_pseudomajus_OL977690.1  11.96%    passed     77.44%
   5  Misopates_orontium_OX326955.1                 12.23%    passed     82.39%
   6  Hippuris_vulgaris_OQ058820.1                  11.87%    failed      0.27%
   7  Callitriche_stagnalis_ON571658.1              12.96%    failed      0.06%
   8  Neopicrorhiza_scrophulariiflora_NC_057075.1   11.95%    passed     82.01%
   9  Veronica_agrestis_NC_068050.1                 13.36%    passed     70.56%
  10  Plantago_afra_MW877575.1                      13.76%    passed     93.84%
  11  Plantago_brasiliensis_MW877579.1              13.57%    failed      3.08%
  12  Plantago_lanceolata_MW877582.1                13.53%    passed     13.37%
  13  Plantago_lanceolata_NC_068049.1               13.62%    passed     11.38%
  14  Plantago_nubicola_MW877564.1                  12.54%    passed     57.19%
  15  Plantago_sericea_MW877584.1                   13.64%    failed      1.54%
****  TOTAL                                         12.73%  4 sequences failed composition chi2 test (p-value<5%; df=3)
NOTE: minimal branch length is reduced to 0.000000576871 for long alignment

Create initial parsimony tree by phylogenetic likelihood library (PLL)... 0.028 seconds
NOTE: ModelFinder requires 92 MB RAM!
ModelFinder will test 286 DNA models (sample size: 173349) ...
 No. Model         -LnL         df  AIC          AICc         BIC
  1  JC            492448.372   27  984950.744   984950.753   985222.446
  2  JC+I          485631.661   28  971319.322   971319.332   971601.088
  3  JC+G4         485490.428   28  971036.855   971036.864   971318.621
  4  JC+I+G4       485422.236   29  970902.472   970902.482   971194.301
```

As output, IQ-TREE generates 9 different files. The FamilyPlusPath.aln.log contains output that has also been printed to our screen, this can be helpful if later on we want to check what happened. In FamilyPlusPath.aln.iqtree we will find the results for the model testing and the most likely tree. At the top of the ModelFinder chapter we find the model which best fits the data we are working with.

```
ModelFinder
-----------

Best-fit model according to BIC: TVM+F+R2
```

Using http://www.iqtree.org/doc/Substitution-Models we can check what the code means and if the assumptions which were made fit with our data. At the end of the file we can see a ASCII representation of our tree, but if we want to take a look and play around with the tree, we can also open FamilyPlusPath.aln.bionj, FamilyPlusPath.aln.contree or FamilyPlusPath.aln.treefile in UGENE or the interactive tree of life (https://itol.embl.de/). There might be a slight difference between the trees, the file which contains the tree we would be most certain about is the FamilyPlusPath.aln.contree. This file contains the consensus tree, which has been bootstrapped and tested.
The last file which might be of interest is the FamilyPlusPath.aln.mldist file, this contains the distances between sequences and can be used if we would want to go for a different method of tree building.

***Compare the tree generated by IQ-TREE to the one created using UGene. Has using IQ-TREE changed your tree?***

***Has using IQ-TREE changed your thoughts on the species you are working on?***

**After all these different steps, what is your final prediction on the species you are working on, add your prediction to this spreadsheet:** https://tinyurl.com/4hsufvn7

***Extra assignment***

If you want and have time, try to find the chloroplast of your favourite plant or flower. You need to find the Latin name of the plant, go to ncbi and find the sequence (https://www.ncbi.nlm.nih.gov/nuccore/), make sure to add chloroplast to your search. The size of the sequence should be around 150.000bp. Add the sequence to the folder and the name of the file to the file with the list of species. Rerun the SANS.

***How related is your favourite species to the plant you sequenced?***
