## Supplementary Protocol 1 for "A workflow for practical training in ecological genomics using Oxford Nanopore long-read sequencing"

### Lab Course Guide:

### NERC Advanced Training in Ecological Genomics

#### April – June 2023

Bioinformatics Demonstrators: Nathan Medd, Heleen De Weerd, Timothy Booth, Urmi Trivedi

### Introduction to Course

Genomic sequencing is used in a multitude of environmental applications, from the monitoring of rare and overlooked species using metagenomics, to detecting responses to climate change using population genomic sequencing. This two-week intensive course will provide a practical introduction to ecological genomics, covering the types of wet lab methods (DNA extraction, library preparation and next generation sequencing) and bioinformatic analyses (data QC, genome assembly, phylogenetics) necessary to go from samples to sequence to results. In this course, you will track a single sample through all stages of the workflow, learning practical skills at each stage. The work will take place at Edinburgh Genomics, using cutting edge genomic equipment, with sequencing using nanopore to obtain results in real time.
After completing the workshop, you should be able to: (1) prepare your own samples and libraries for genomic sequencing, (2) perform bioinformatic analyses of next generation sequencing data, (3) apply your new skills to other sequencing methods and analysis problems.

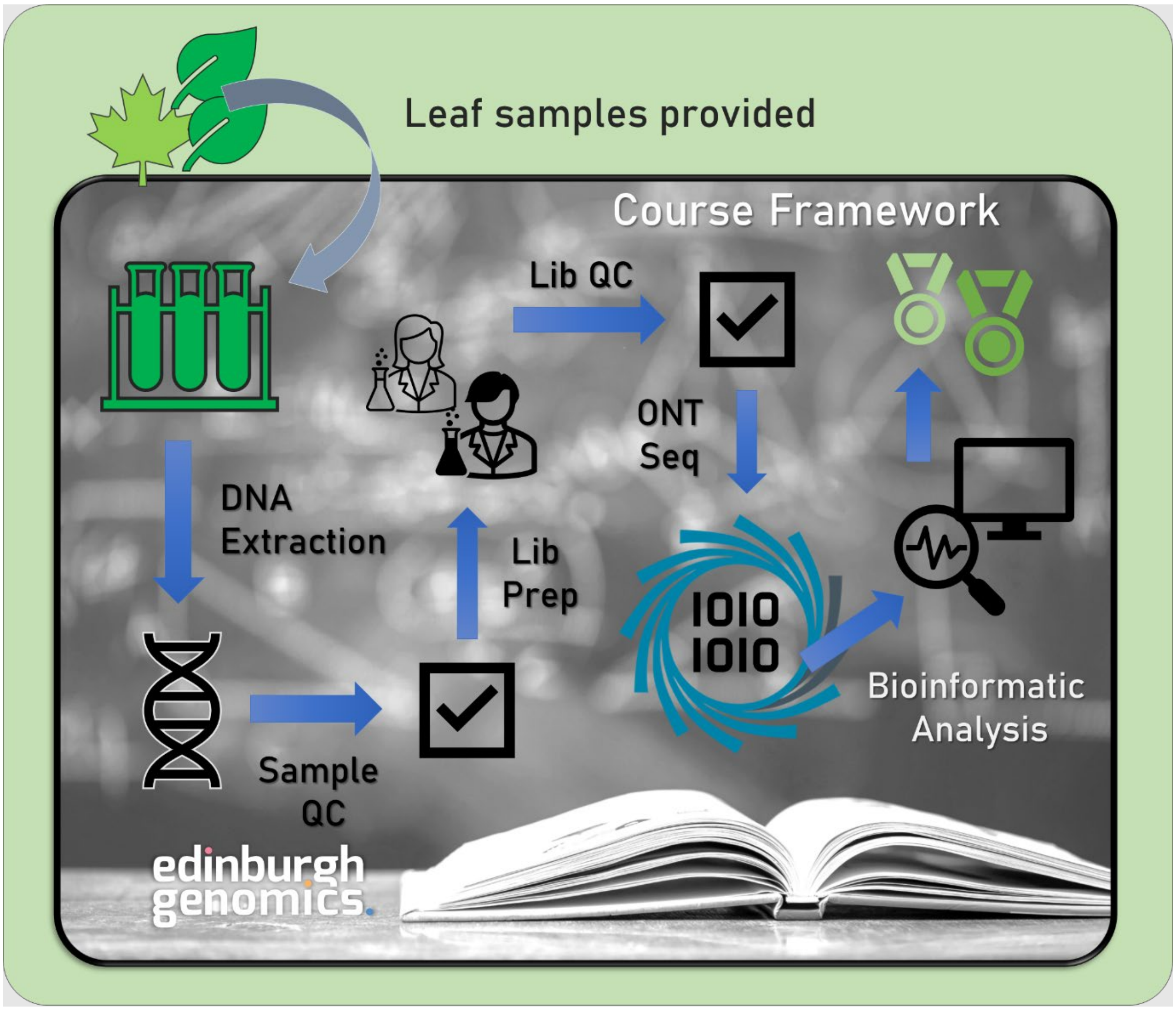

### Overview Timetable

**1. Monday**
09.30 Arrival, setup
10.00 Intro to the course, background to ecological genomics. DNA extraction (AT)
10.30 Coffee
11.00 Lab intro, tour, Q&A (JSL)
12.00 Lunch
13.00 Sample grinding (RF)
14.00 DNA extraction & clean-up (RF)
16.30 DNA extraction discussion
17.00 Finish

**2. Tuesday**
09.30 Background to library prep (JSL)
10.30 Coffee
11.00 DNA QC
11:30 DNA QC Discussion
12:00 Lunch
13:00 Library preparation
15:00 Library QC
16:00 Library prep results discussion, panel Q&A on user sample types.
17:00 Finish

**3. Wednesday**
09.30 Background to genomic sequencing (AT)
10.30 Coffee
11.00 Flongle class demo
12.30 Discussion of results
13.00 Finish

*Bioinformatics week follows on Wednesday – Friday – see separate guide*.

### Health & Safety Overview

While the following protocols have been specifically chosen/designed to avoid most harmful chemicals, it is critical to follow proper health & safety protocols while in the lab. You will have been provided with a full risk assessment the week prior to beginning the course – make sure you have read this fully and understand what to do in case of an accident.

- Make sure you are always wearing proper safety equipment when in the lab, including a lab coat and gloves. Safety goggles are also available on request.
- Only carry out the lab work when a demonstrator is present. If you have any doubts, always check with a demonstrator first.
- Important safety features (e.g. the eye wash station) will be pointed out to you when you first enter the lab. Make sure to familiarise yourself with these.
- There is a fire alarm test at ~4PM on Tuesday. Any alarm outside of this time should be treated as real and the building should be evacuated. Lab work should be abandoned quickly and safely, and lab coats and gloves removed before quickly exiting the building via the fire doors.
- Hazardous chemicals will be labelled. Pay special attention when working with these.
- Be careful when handling tubes and vials to avoid spills. When you are not using a tube or vial it is always best to seal it fully as this also avoids contamination.
- Dry ice is incredibly cold and can ‘burn’ quickly when in contact with bare skin. Even when gloved, avoid contact entirely if possible.
- No food or drink is allowed in the lab.

### Lab Work Overview

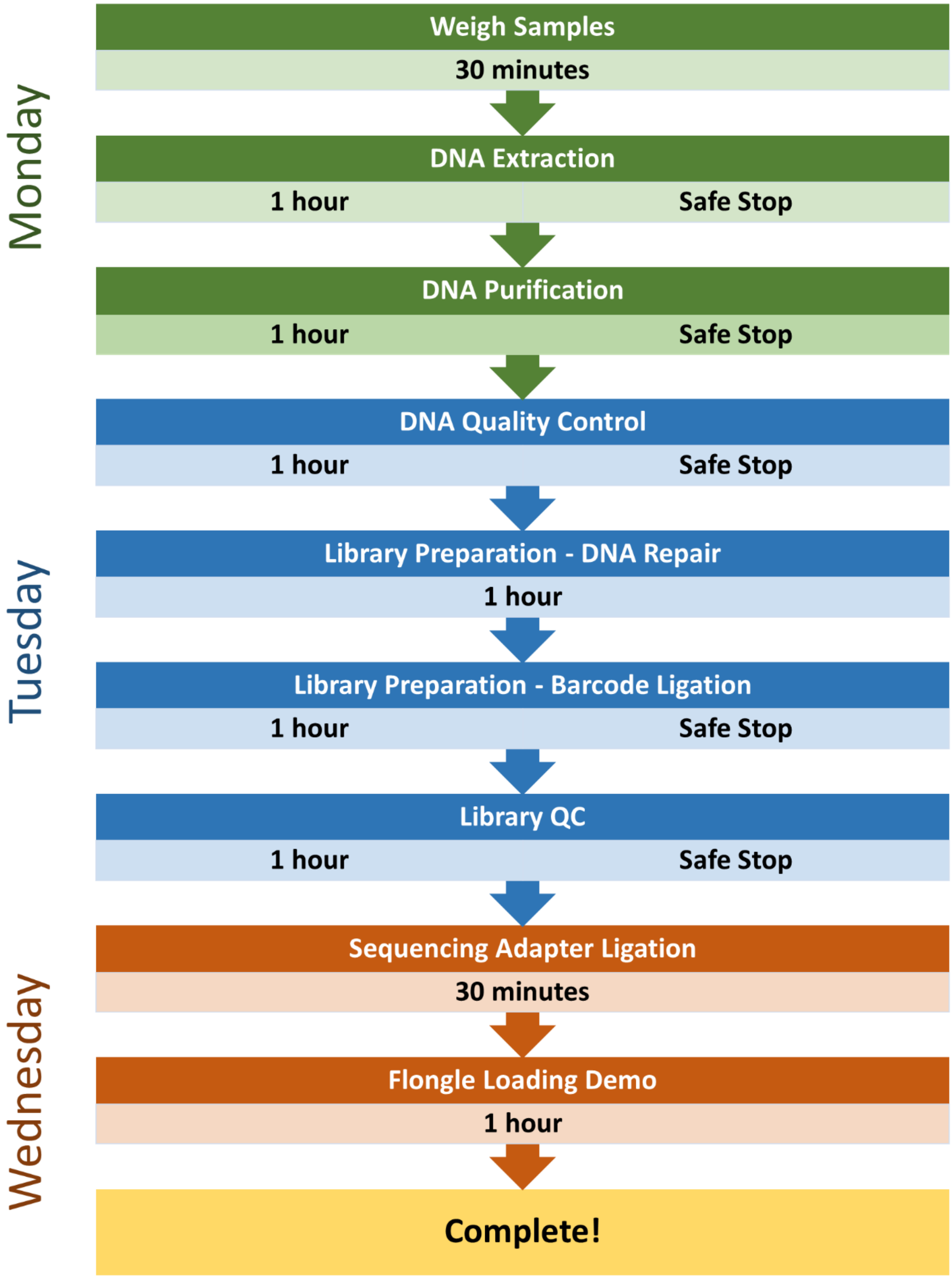

#### 1. Day 1 – Weighing Samples

- *Ensure you have a bucket of dry ice ready. The level of dry ice should be high enough to submerge an Eppendorf tube. NB. Dry ice should not be directly handled, even with gloved hands.*

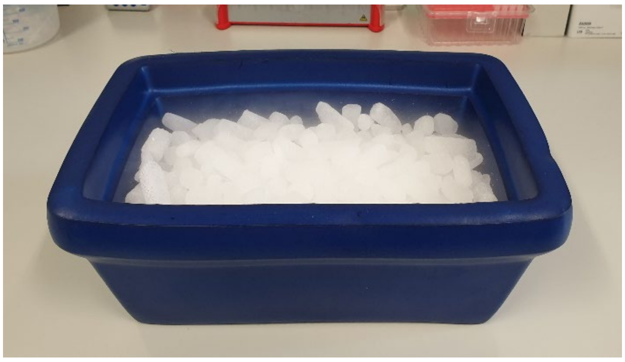

☐ Locate your grinding tube and disposable pestle.

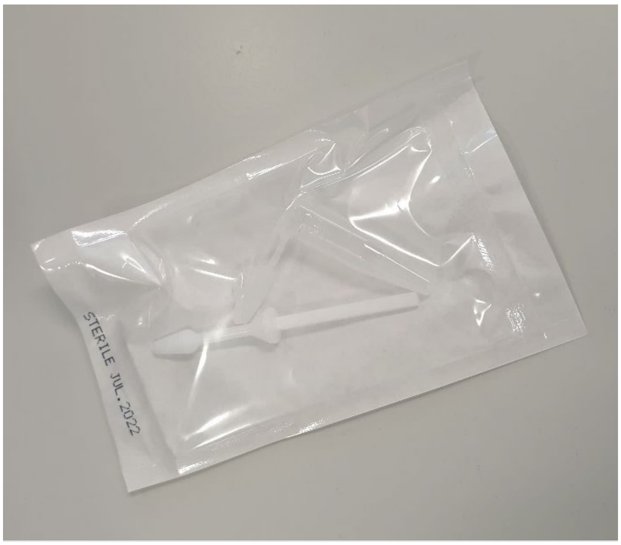

☐ Carefully remove the grinding tube from the sealed packet and submerge it in the dry ice to cool. NB. In the image below I have marked the tube for clarity.

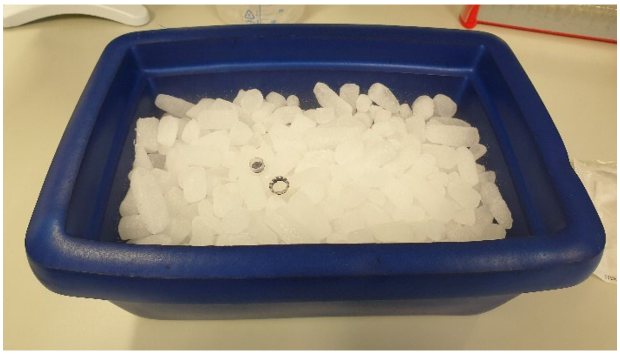

☐ Locate the scales and weighing boats. You may have to wait for other trainees to finish this step. Slide open a side-door and place a weighing boat in the centre of the scales.

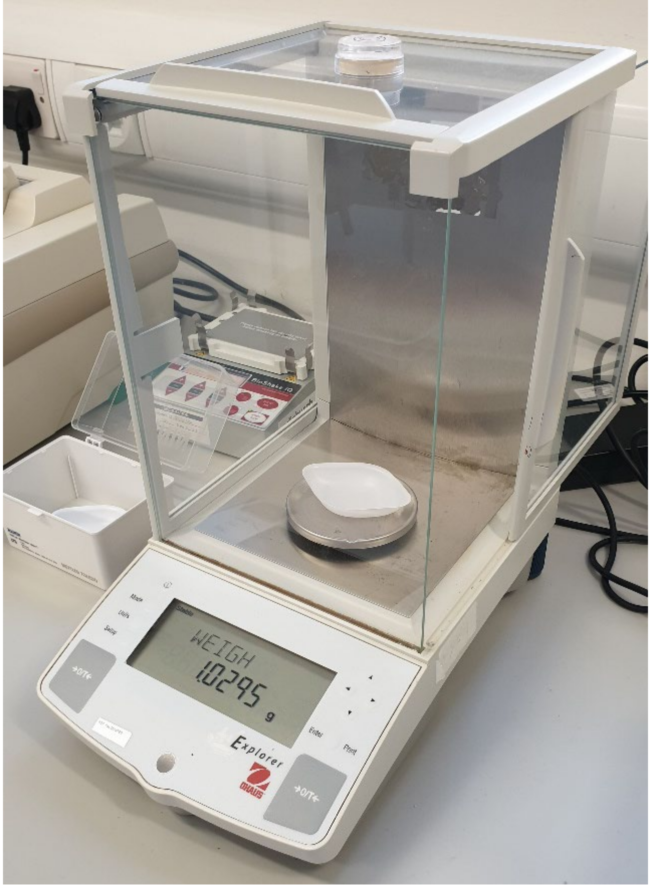

☐ Close the side-door and tare the scales by pressing the 'O/T' button. The weight reading should become close to zero, although this will fluctuate slightly.

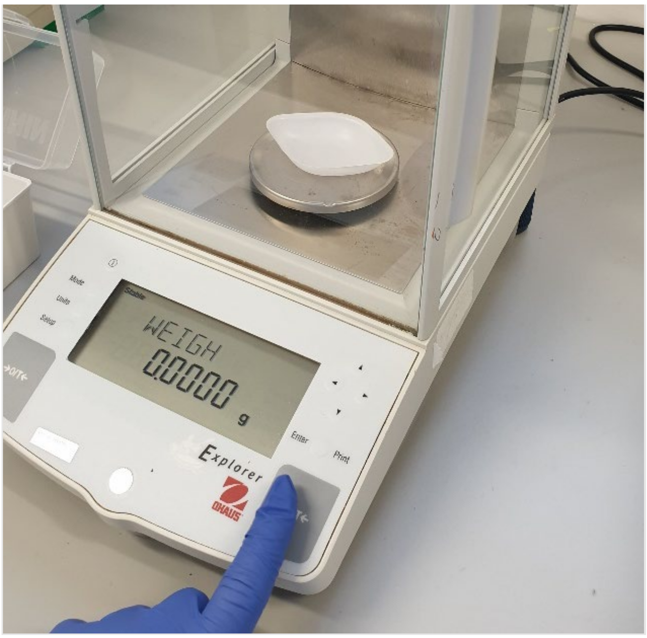

☐ Locate the tweezers and your allocated plant sample. This consists of several dried leaves in grains of silica.

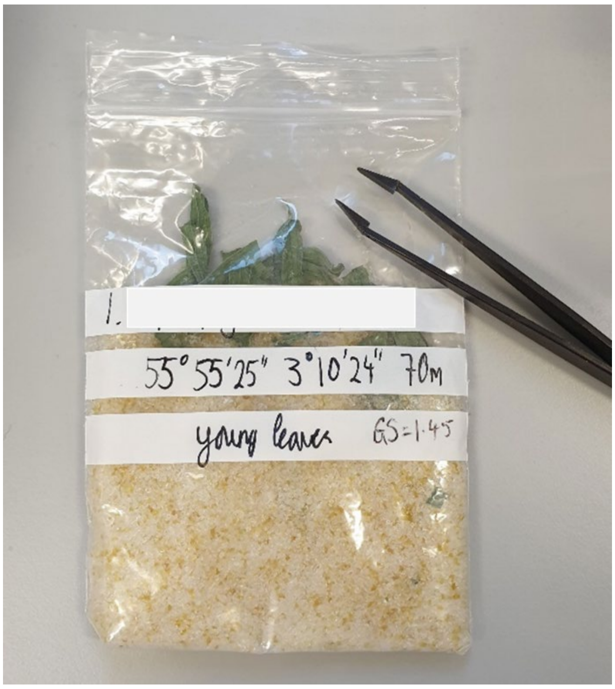

☐ **Carefully** remove leaves, or portions of leaves, from the silica grains and place on the weighing boat on the scales. NB. Leaves are dry and brittle and will break if too much pressure is applied! Try to shake off any grains attached to the leaves if possible. Attempt to get approximately 15mg of dried leaf tissue (0.0150 g). You may deliberately break leaves using pressure from the tweezers to achieve this.

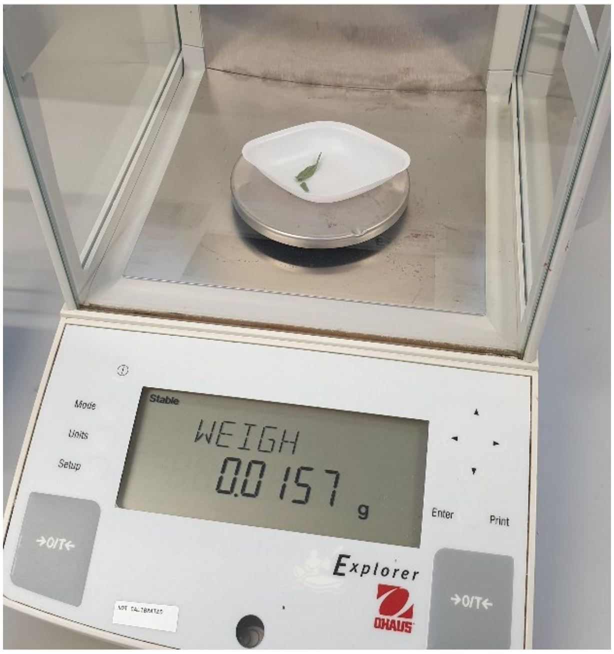

☐ Transfer the weighing boat with ~15mg leaf tissue over to your dry ice. By applying pressure to either side of the weighing boat, you can create a funnel. Use this to empty the weighing boat into your chilled grinding tube. Allow leaf sample to cool for ~1

minute prior to beginning the grinding stage.

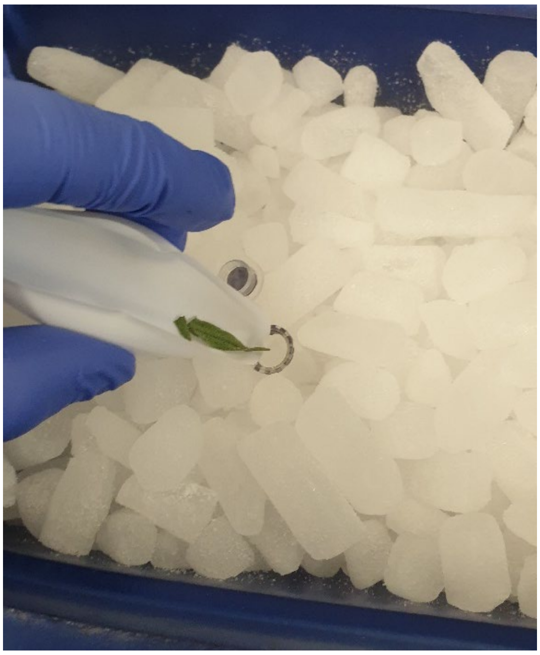

#### 2. Sample Grinding

☐ Locate 'PowerMasher II' device. Remove the disposable pestle from the sleeve and insert it into the transparent plastic holder as shown. Ensure the pestle is fully secured.

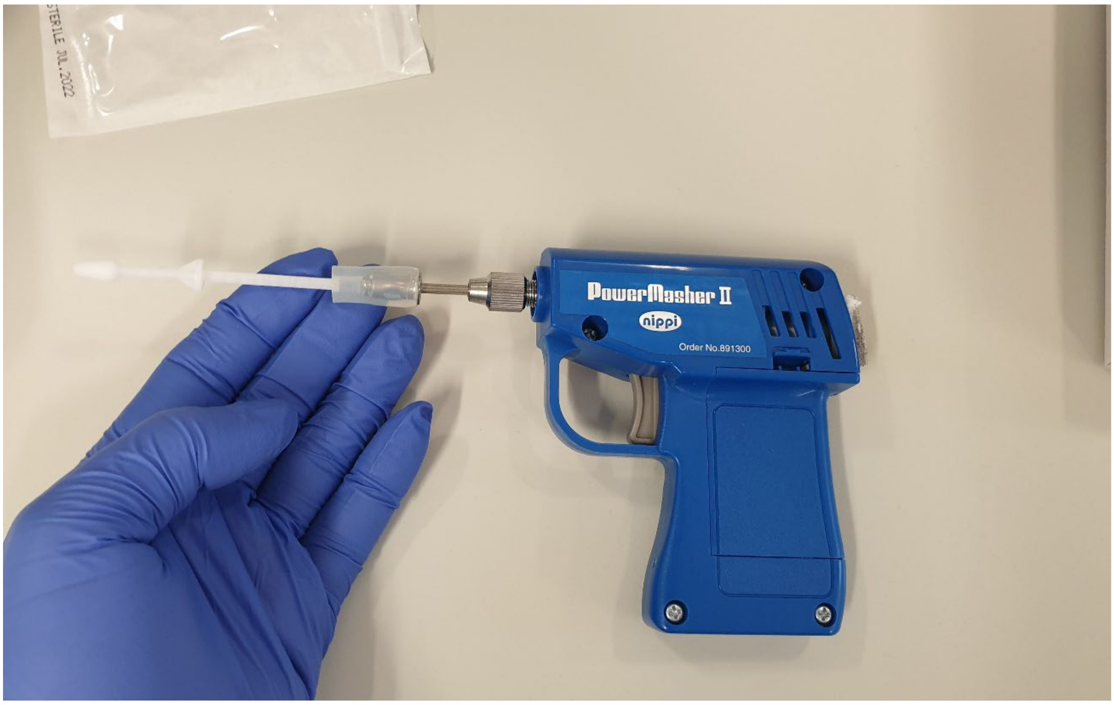

☐ Returning to the grinding tube containing your leaves, use the end of the pestle to compress the leaf sample into the bottom of the tube.

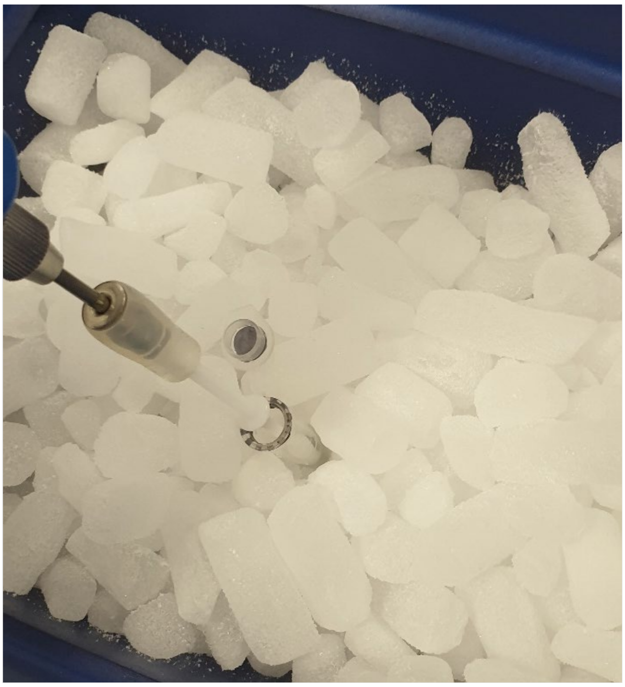

☐ Perform 10 second bursts of grinding by pressing the trigger on the PowerMasher II while simultaneously compressing the leaves into the bottom of the grinding tube. After grinding for 10 seconds, take a 10 second break, this is to ensure the heat generated does not thaw the sample which can impede the creation of a fine powder.

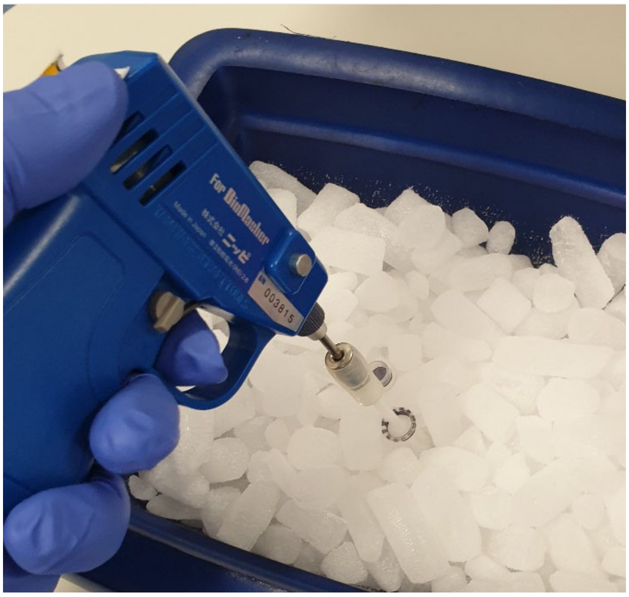

☐ During each 10 second break, use the pestle to push down any leaf fragments which have travelled up the side of the tube, ensuring all leaf matter is in contact with the rough surface at the bottom of the grinding tube.

☐ Repeat steps III and IV for 10 minutes, or until a fine powder has been obtained. Verify this with one of the lab demonstrators. A fine powder will give a higher yield from the DNA extraction in the following steps. Keep this on dry ice until proceeding.

### 3. DNA Extraction

- *DNA extractions will be performed with the DNeasy Plant Mini Kit (QIAGEN). You will find aliquots of the necessary reagents from the kit on your bench, along with the required plasticware. You should also have a small tray of ice ready (not dry ice).*

☐ Remove the grinding tube with leaf powder from the dry ice and place in a tube rack.

☐ Using a P1000 pipette, add 400 µl AP1 to the grinding tube. Use the pipette tip to dislodge any clumps of leaf material at the bottom of the grinding tube.

☐ Immediately, using a P20 pipette, add 4 µl RNase A to the grinding tube.

☐ Seal the grinding tube using the attached lid.

☐ Using the vortexer on your bench, vortex the tube for 10 seconds. Press the tube into the pad to activate. NB. You may have a different model of vortexer than below. The leaf powder should now be homogenously suspended in the lysis mixture.

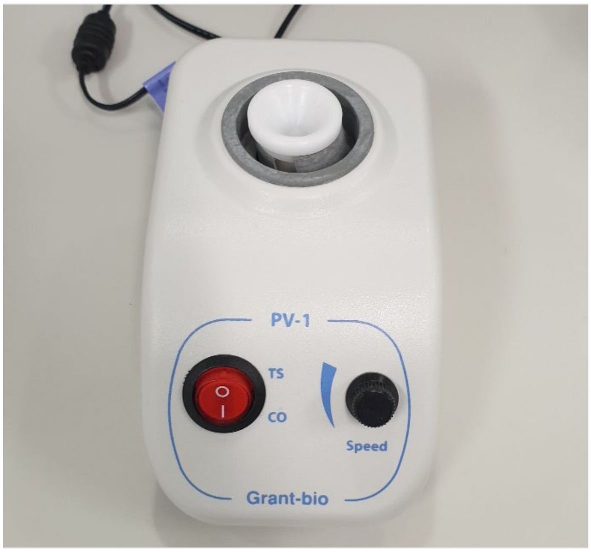

☐ Transfer the lysis mixture into a heat block set to 65°C for 5 minutes (set a timer). After 5 minutes, vortex the tube again as described above.

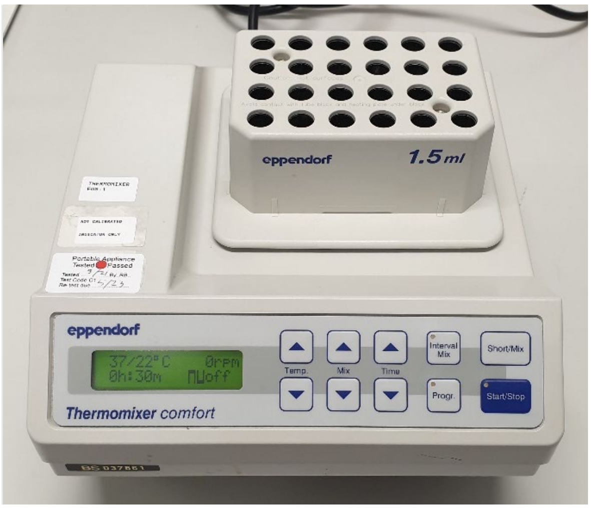

☐ Return the vortexed tube to the heat block for another 5 minutes at 65°C, then transfer to ice.

- ☐ Immediately add 130 µl P3 using a P200 pipette. Switch to a 200µl wide-bore pipette tip and mix thoroughly by pipetting up and down 10 times. Leave this to incubate for 5 minutes on ice to neutralise the lysis reaction.
- ☐ Transfer the tube containing the lysate to a centrifuge and run at 20,000 x g (rcf) for 5 minutes. A centrifuge **must** be balanced (approximately equal weights on opposite sides) before it is run – speak to a demonstrator if you are unsure. Make sure to seal the centrifuge with the detachable lid.

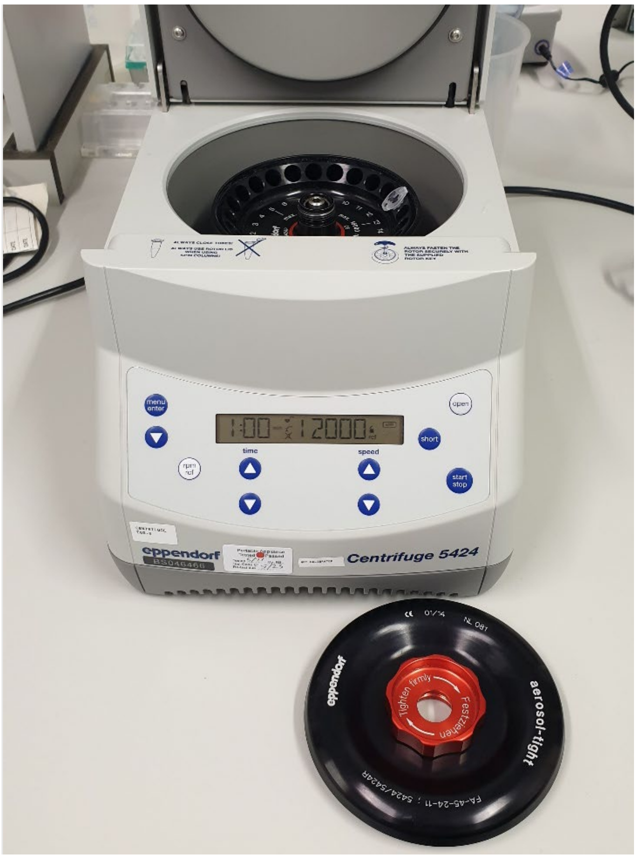

- ☐ Remove the tube from the centrifuge. A 'pellet' of unwanted leaf material should be present at the bottom of the tube, the DNA will remain in solution. Using a P1000 pipette, transfer the solution, avoiding the pellet, into a 'QIAshredder spin column' in a 2ml collection tube (sealed packet, provided).
- ☐ Centrifuge the QIAshredder spin column in collection tube for 2 minutes at 20,000 x g (rcf).
- ☐ Remove the column and collection tube from the centrifuge. A faint pellet may be visible in collection tube. Retain the collection tube containing the flow-through and discard the spin column.
- ☐ Using a P1000 pipette, transfer the flow-through into a new 1.5ml Eppendorf tube, using gentle aspiration to avoid disturbing the pellet, if present. There should be ~ 450 - 500 µl of flow-through.
- ☐ Using a P1000, add 700 µl of AW1 to the flow through and mix by pipetting 5 times.
- ☐ Locate the 'DNeasy Mini spin column' in 2ml collection tube (sealed packet, provided). Transfer 650 µl of the flow-through/AW1 mixture into the spin column.

☐ Centrifuge this for 1 minute at 7000 x g (rcf). Discard the flow-through in the collection tube (while retaining the collection tube and spin column).
☐ Add the remaining AW1 mixture to the spin column in the collection tube and centrifuge again for 1 minute at 7000 x g . Discard the flow-through.
☐ Place the spin column into a new 2ml collection tube (old collection tube can be discarded).
☐ Add 500 µl AW2 and centrifuge for 1 minute at 7000 x g. Discard the flow-through in the collection tube (retain tube and spin column).
☐ Add an extra 500 µl AW2 to the spin column plus collection tube and centrifuge for 2 minutes at 20,000 x g.
☐ Transfer the spin column to a new 1.5ml Eppendorf tube, being careful to avoid any flow-through. If flow-through comes into contact with the spin column this can be removed by dabbing with a tissue.
☐ For elution, add 55 µl AE to the spin column in 1.5ml Eppendorf tube. Incubate at room temperature (15 – 25°C) for 2 minutes. Centrifuge for 1 minute at 7000 x g.
☐ Repeat the above step, adding 55 µl AE to the spin column in the same 1.5ml Eppendorf tube. Incubate at room temperature (15 – 25°C) for 2 minutes. Centrifuge for 1 minute at 7000 x g.
☐ This Eppendorf now contains your eluted DNA. Label it clearly with your initials and store it on ice until proceeding to DNA purification.

- ***Safe stop point.***

#### 4. DNA Purification

- *We will be cleaning up our DNA extractions using the DNeasy PowerClean Pro Cleanup Kit (QIAGEN). For plant samples especially, it is necessary to remove certain compounds including polysaccharides and polyphenols which co-extract with the DNA and can reduce sequencing yield. You will find aliquots of the necessary reagents from the kit on your bench, along with the required plasticware.*

☐ Transfer 100 µl of your DNA extraction into a clean 2ml collection tube.
☐ Add 50 µl of CU to the DNA extraction and vortex briefly (5 seconds) to mix.
☐ Add 50 µl of IR and vortex briefly to mix.
☐ Centrifuge the tube at 13,000 x g for 2 minutes.
☐ A pellet (which contains potential contaminants) should be visible in the tube. Taking care to avoid this pellet, use a P200 to transfer ~190 µl of supernatant to a clean 2ml collection tube.
☐ Add 400 µl of SB to the supernatant and vortex briefly to mix.

- ☐ Using the benchtop mini-centrifuge ('minifuge'), spin down the tube for ~2 seconds to remove any liquid from the cap. The minifuge is activated by closing the lid.

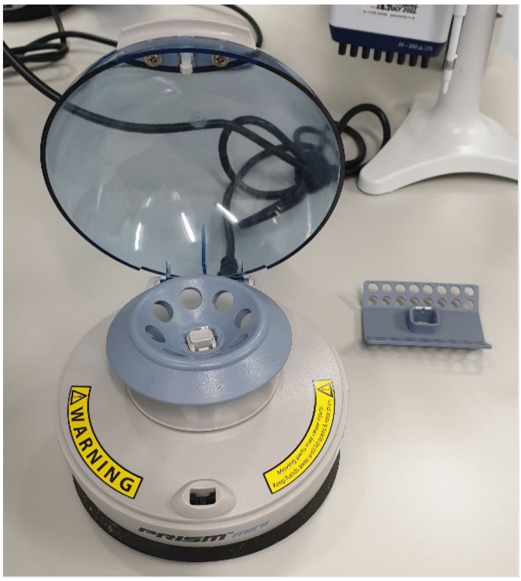

- ☐ Transfer the mixture (~590 µl) into a provided MB Spin Column and centrifuge at 10,000 x g for 1 minute. Discard the flow-through (retaining the spin column).
- ☐ Add 500 µl of CB to the MB Spin Column and centrifuge at 10,000 x g for 1 minute. Discard the flow-through.
- ☐ Repeat the previous step (X).
- ☐ Centrifuge the MB Spin Column at 20,000 x g for 2 minutes.
- ☐ Avoiding the flow-through, transfer the MB Spin Column to a new 2ml collection tube.
- ☐ Add 30 µl of EB to the (white) centre of the spin column filter membrane. Incubate for 2 minutes at room temperature.
- ☐ Centrifuge at 10,000 x g for 1 minute.
- ☐ Discard the MB Spin Column, keeping the collection tube. This collection tube contains your purified DNA – seal it and label it clearly with your initials. It will be stored at -20°C ready to go into library preparation tomorrow.

- ***Safe stop point.***

### 5. Day 2 – DNA QC

- *Prior to starting library preparation, we need to check the quantity and quality of the DNA you have extracted. For long-read sequencing, we want to start with long DNA (>10kb) and so we will size the DNA extraction using a Tapestation (Agilent) and Genomic Screentapes. This is essentially a very compact and quick gel electrophoresis system which produces an automated digital report. We also want to start the protocol with equal inputs of DNA (400 ng), and so we measure the concentration (ng/µl) with a Qubit Broad-range DNA Assay Kit. The Qubit reagent contains a fluorescent dye* *which binds to the DNA and allows quantification in the Qubit device.*

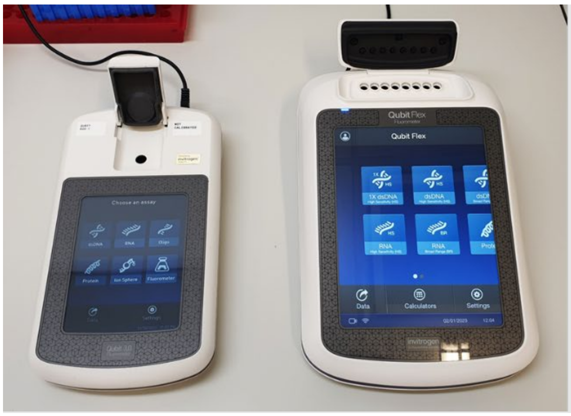

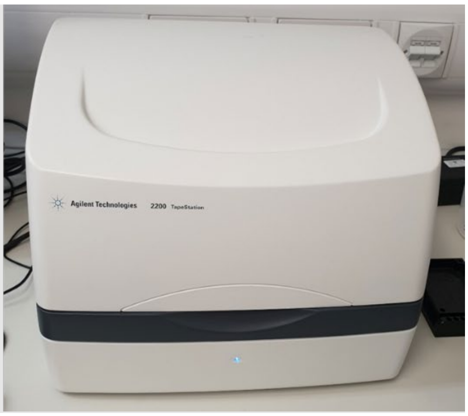

- *For Qubit, first we prepare a master-mix of the Qubit reagent diluted 1 in 200 in Qubit buffer. This dilution has been provided for you. 199 µl of this reagent has been aliquoted into 8-well strips (also provided).*

☐ To this, you need to add 1 µl of your DNA extraction in wells 1 – 3, using a P2 pipette. Seal the tube strip and mix by ‘flicking’ the strip several times. Spin down this tube strip briefly (2 seconds) using the bench-top minifuge with the tube-strip adapter.

☐ At your turn, place the tube strip into the Qubit Flex with your sample to the left. Using the touch screen, select the left-most 3 positions if not already selected. Press Next. Ensure sample volume is set to 1 µl. Press Run Samples – this will take a few moments. The concentration of your 3 replicates will appear on screen – take a note of these measurements and use them to calculate an average concentration for your sample. Relay this average concentration to one of the demonstrators.

- *The lab demonstrators will prepare the samples for sizing on the Tapestation to allow them to be run together. Hand your sample to a demonstrator following the Qubit stage. Your sample will be mixed with Tapestation buffer and then run on a Screentape – a small plastic device which contains several gel-filled channels for electrophoresis. The demonstrators will go over the results with you when ready.*

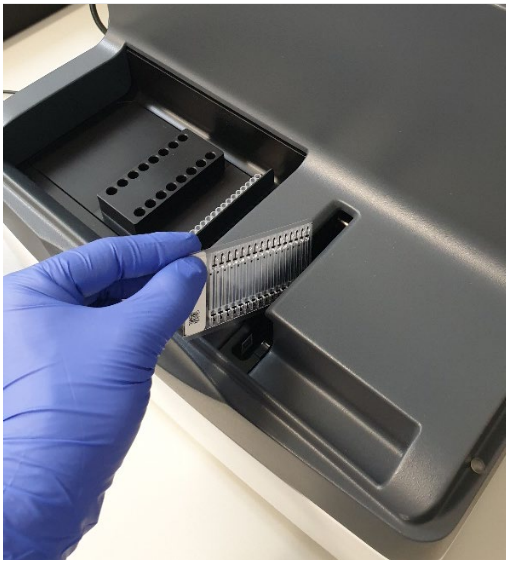

### 6. Library Preparation

- *We will be using the Native Barcoding Kit 24 V14 (SQK-NBD114.24, Oxford Nanopore) to perform library preparation. An overview of the process is given below. For this course we will not be carrying out fragmentation (which reduces the size of the DNA). However, we will first repair the DNA you have produced using an enzyme cocktail to remove minor damage and then end-prep – add A-overhangs. These overhangs allow ligation of barcodes in the following step (required to identify your sample from the pool bio-informatically following sequencing). An adapter complex is then added, which contains the motor protein which drives the DNA fragment through the nanopore.*

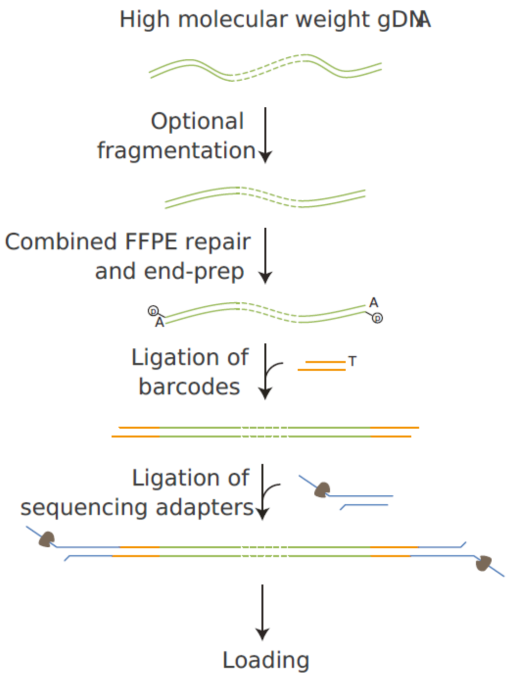

(Source: ONT)

- *You will be provided with the library preparation reagents in a labelled box placed on ice.*

☐ You will need to prepare a 400 ng dilution of your DNA extraction in 24 µl of nuclease-free water (16.7 ng/µl). If you have a lower concentration than 16.7 ng/µl, simply transfer 24 µl to a new tube. If you have less than 100 ng DNA in 24 µl (4.2 ng/µl), see a demonstrator.

- *This dilution is based on the average concentration you calculated during sample QC yesterday. The formula for calculating the volume of sample you require ($V_1$) is: $C_1V_1 = C_2V_2$, re-arranged: $V_1 = C_2V_2/C_1$ . From above, $C_2$ is 16.7, $V_2$ is 24 and $C_1$ is your average concentration. $24 - V_1$ will give you the volume of nuclease-free water for dilution. Check your sums with a demonstrator if unsure.*

☐ Once you have diluted your sample, seal the tube and mix by flicking the tube several times. Briefly spin down the tube in the bench-top minifuge.

☐ Add the following components to your DNA in a 1.5ml Eppendorf, in the order listed:

| Reagent | Volume (µl) |
|---|---|
| DNA | 24.0 |
| DNA Repair/End-prep Master-mix | 6.0 |
| Total | 30.0 |

- ☐ Pipette mix 10 times using a wide-bore 200 µl tip (set to 20 µl).
- ☐ Transfer the mixture to the 20°C heat block (Thermomixer) and set a timer for 5 minutes.
- ☐ After 5 minutes, transfer the tube to the 65°C heat block and set another 5 minute timer.
- ☐ After 5 minutes, return the tube to room temperature on your bench.
- *Locate the tube of AMPure XP beads. These are magnetic beads which bind DNA and are used for purification (i.e. separating your DNA from the enzymes you've just added). There will be a brown precipitate in this tube, which is the beads.*
- ☐ Resuspend the beads by thoroughly vortexing on the bench-top vortexer for at least 20 seconds. The solution must appear homogenous before continuing.
- ☐ Add 30 µl of the re-suspended AMPure XP beads to your DNA/DNA Repair mixture. Mix thoroughly using a wide-bore tip.
- ☐ Incubate on a rotator mixer for 5 minutes at room temperature. If the mixer is currently running, you may briefly stop it using the power switch on the side. Tubes are pushed into the metal holders. Ensure the cap is tightly sealed.

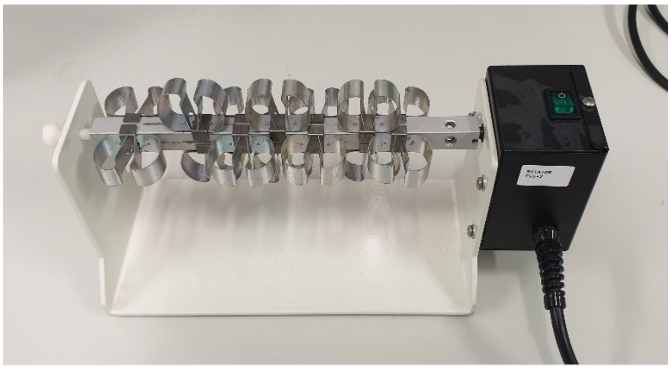

- ☐ After 5 minutes, remove your tube from the rotator and briefly spin down in the minufuge. Place your tube onto a magnet and wait until the beads pellet and the supernatant becomes transparent (approximately 2 minutes).

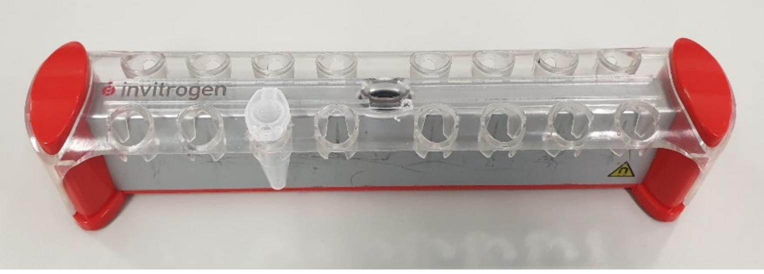

- ☐ With your tube still on the magnet, pipette off the supernatant and discard.
- ☐ Wash the beads with 500 µl of 80% ethanol (provided in bijou). Pipette the ethanol down the side of the Eppendorf **opposite** the DNA pellet. Wait 30 seconds and then gently remove the ethanol.

☐ Repeat previous step.
☐ Remove the tube from the magnet and briefly spin it down in the minifuge.
☐ Return the tube to the magnet and allow the beads to pellet (~30s). Gently remove any residual ethanol at the bottom of the tube with a P20 pipette.
☐ Remove the tube from the magnet and place in a tube rack.
☐ Using a P20, add 9 µl of nuclease-free water and re-suspend by pipetting 10 times. Incubate on your bench for 2 minutes at room temperature to elute your DNA.
☐ Return the tube to the magnet and allow beads to pellet. The supernatant now contains your DNA. Carefully transfer 8 µl of supernatant to a fresh 1.5 ml Eppendorf tube. Store on ice if not immediately proceeding to Barcode Ligation.

- ***Safe stop point.***

### 7. Native Barcode Ligation

- *You have been provided with a Native Barcode for your sample and the Ligase on ice. You should take a note of your barcode number.*

☐ Add the following components to your end-prepped DNA in the order listed:

| Reagent | Volume (µl) |
|---|---|
| **End-prepped DNA** | 7.5 |
| **Native Barcode** | 2.5 |
| **Blunt/TA Ligase Master Mix (LIG)** | 10.0 |
| **Total** | 20.0 |

☐ Mix thoroughly using a wide-bore 200 µl tip (Set P200 to 20 µl and pipette mix 10x).
☐ Place the tube in a heat block set to 20°C for 20 minutes.
☐ Remove tube from heat block and return to bench.
☐ Add 2 µl of EDTA to stop the reaction.
☐ Resuspend AMPure XP beads by vortexing (20s).
☐ Add 20 µl of resuspended AMPure XP beads to the reaction and mix thoroughly with a wide-bore pipette.
☐ Incubate on a rotator mixer for 5 minutes at room temperature. If the mixer is currently running, you may briefly stop it using the power switch on the side.
☐ After 5 minutes, remove your tube from the rotator and briefly spin down in the minufuge. Place your tube onto a magnet and wait until the beads pellet and the supernatant becomes transparent (approximately 2 minutes).
☐ With your tube still on the magnet, pipette off the supernatant and discard.
☐ Wash the beads with 500 µl of 80% ethanol (provided in bijou). Pipette the ethanol down the side of the Eppendorf **opposite** the DNA pellet. Wait 30 seconds and then gently remove the ethanol.
☐ Repeat previous step.
☐ Remove the tube from the magnet and briefly spin it down in the minifuge.
☐ Return the tube to the magnet and allow the beads to pellet (~30s). Gently remove any residual ethanol at the bottom of the tube with a P20 pipette.

☐ Remove the tube from the magnet and place in a tube rack.

☐ Using a P200, add 30 µl of nuclease-free water and re-suspend by pipetting 10 times. Incubate at 37 °C for 5 minutes (in a heat block) to elute your DNA.

☐ Return the tube to the magnet and allow beads to pellet. The supernatant now contains your DNA. Carefully transfer 28 µl of supernatant to a fresh 1.5 ml Eppendorf tube. Store on ice if not immediately proceeding to Library QC.

- ***Safe stop point.***

#### 8. Library QC

- *Library QC will be performed similarly to sample QC. The concentration of your library will be measured using Qubit and the size will be measured using the Tapestation. Based on the size and concentration we can calculate the molarity of the prepared library. We can use the molarities to calculate the volume of each library required to create a balanced pool (the pool is the combined barcoded sequencing libraries). This ensures we get the desired coverage of each sample based on their genome size.*
- *As before, for Qubit we have prepared a master-mix of the Qubit reagent diluted 1 in 200 in Qubit buffer. This dilution has been provided for you. 199 µl of this reagent has been aliquoted into an individual Qubit tube (to preserve library we will be only taking 1 measurement).*

☐ To this Qubit tube you need to add 1 µl of your library using a P2 pipette. Mix using a P200 pipette and seal the tube. Spin down this tube briefly (2 seconds) using the bench-top minifuge.

☐ At your turn, place the Qubit tube into the Qubit 3.0. Using the touch screen, press Read Tube in the bottom right. After a few moments the Qubit will display the concentration. Take a note of the concentration of your library. Relay this concentration to one of the demonstrators.

- *The lab demonstrators will prepare the libraries for sizing on the Tapestation to allow them to be run together.*

☐ Hand your sample to a demonstrator following the Qubit stage. Your sample will be mixed with Tapestation buffer and then run on a Screentape, as before. The demonstrators will go over the results with you when ready (the run takes approximately 15 minutes).

- *Your library QC results will be used to create a balanced pool for adapter ligation and loading tomorrow.*

#### 9. Day 3 – Adapter Ligation

- *A 700ng pool of your barcoded libraries has been created, balanced based on the genome size of the species. The average size of this pool has been measured. As there is now only one tube, this stage of the protocol will be performed by a demonstrator.*
- *In addition to the barcoded pool, the following is required on ice: Elution Buffer (EB), Quick Ligation Buffer, T4 Ligase and Native Adapter (NA). We are also working with*

*Long Fragment Buffer (LFB) at room temperature (this is used to remove small fragments <3kb) and a vial of AMPure XP beads.*

- ☐ Prepare the adapter ligation mix by adding the following components in the order shown to the barcoded pool:

| Reagent | Volume (µl) |
|---|---|
| **700ng pooled barcoded library** | 60 |
| **Native Adapter (NA)** | 10 |
| **Quick Ligation Buffer** | 20 |
| **Quick T4 DNA Ligase** | 10 |
| **Total** | 100 |

- ☐ Mix the adapter ligation mix 10x using a P200 wide-bore pipette tip (set to 70µl) and briefly spin down in the minifuge.
- ☐ Incubate the reaction for 10 minutes at room temperature.
- ☐ Resuspend the AMPure XP beads by thoroughly vortexing (at least 20 seconds). The solution must appear homogenous.
- ☐ Once the 10 minute incubation is complete, add 50 µl of resuspended AMPure XP beads to the reaction using a standard 200 µl pipette, then mix thoroughly using a wide-bore 200 µl tip.
- ☐ Incubate the tube on the rotator for 5 minutes.
- ☐ Transfer the tube to a magnetic rack and allow the beads to pellet and the supernatant to become transparent. This may take a couple of minutes.
- ☐ Remove and discard the supernatant.
- ☐ Immediately remove the tube from the magnetic rack and re-suspend the beads in 250 µl Long Fragment Buffer (LFB) using a P1000 pipette.
- ☐ Return the tube to the magnetic rack and allow the beads to pellet. Remove and discard the supernatant.
- ☐ Repeat the previous 2 steps (XI, XII).
- ☐ Remove tube from magnetic rack and allow the pellet to dry for 1 minute. Immediately re-suspend in 28 µl Elution Buffer (EB). Pipette mix 10x using a wide-bore tip. Incubate the elution at 37°C for 10 minutes in a heat-block.
- ☐ Transfer the tube to the magnetic rack and leave for 1 minute for the beads to pellet. The supernatant now contains the library – **do not discard!** Transfer 27 µl of the supernatant, slowly to avoid aspirating the beads, into a fresh 1.5ml Eppendorf tube.

- *We will quantify 1 µl of this library using the Qubit 3.0 and high-sensitivity DNA assay. Based on the average size of the library, we will then calculate the amount of femtomoles present. We will load 1µl of the recovered library on the Flongle as a test, and the remainder (ideally 10-20 femtomoles) on the PromethION flow cell.*

### 10. Flongle Loading Demonstration

- *The Flongle is a small flow-cell used mainly for testing libraries. It connects via an adapter to the MinION device. We will demonstrate loading of the Flongle with the prepared library, and go over some of the initial QC results and data.*

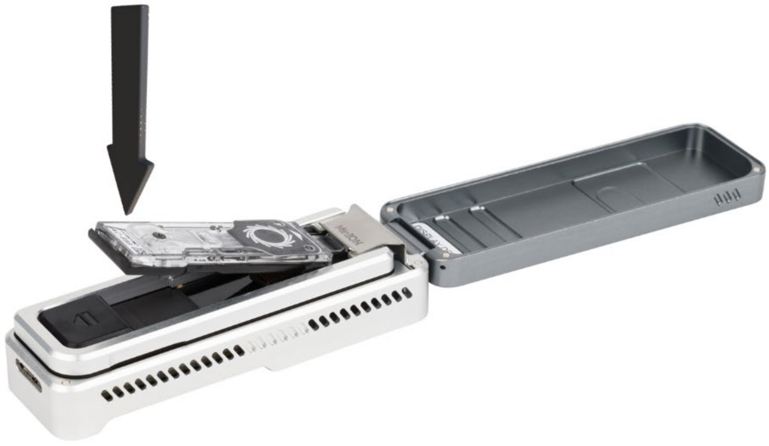

(Source: ONT)

- *The Flongle has several openings and ports. The most critical components to take note of are the plastic seal tab and the sample port.*

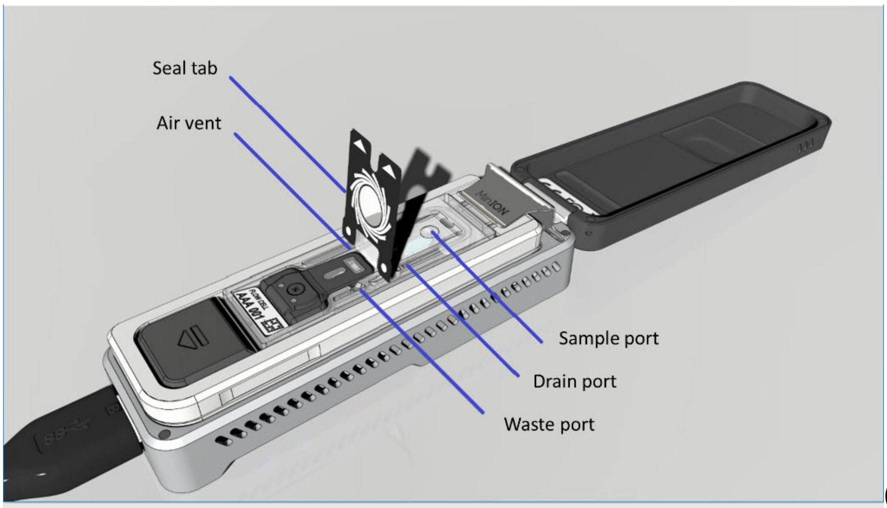

(Source: ONT)

- ☐ Firstly, we prepare a priming mix consisting of 117 µl Flow Cell Flush (FCF) and 3µl Flow Cell Tether (FCT). This contains the ATP which drives the motor protein and the tether which allows it to bind to the pores.
- ☐ We peel back and stick down the seal tab to expose the sample port.
- ☐ Nanopore flow cells are extremely sensitive to air exposure. To ensure we do not introduce any air into the sample port, we aspirate out ~30µl of the storage solution using a P200 pipette.
- ☐ We then prime the flow cell using the 120 µl priming mix. After taking the priming mix up into the pipette, we twist the pipette plunger to expel a tiny amount of mix and

ensure there is no air at the bottom of the pipette tip. We gently introduce the priming mix into the Flongle via the sample port by twisting the pipette plunger.

☐ The DNA library is then prepared for loading:

| Reagent | Volume (µl) |
|---|---|
| **Sequencing Buffer (SB)** | 15 |
| **Library Beads (LIB)** | 10 |
| **EB** | 4 |
| **DNA Library** | 1 |
| **Total** | 30 |

☐ The sequencing mix is then introduced into the flow cell similarly to the priming mix and the seal tab is returned to its original position. The sequencing platform lid is closed and the sequencing run is set-up in the MinKnow software.

☐ ONT sequencing devices output sequencing data in real time, following a short ~5 minute delay where the flow-cell comes to temperature and is checked at the start of a run. Within the first 15 minutes we will have QC results for the flow cell and some data on the first fragments to pass through the pores!

### Working with MinKNOW

- *MinKNOW is ONT’s software for setting up and monitoring sequencing runs.*

#### Flow Cell QC

1. The first stage of starting a run is to check your flow cell. ONT guarantees flow cells to have a minimum number of active pores, and will replace flow cells which do not meet this standard within 3 months of receipt.
2. After correctly inserting your flow cell into the MinION device, select ‘Flow cell check’ on the display.

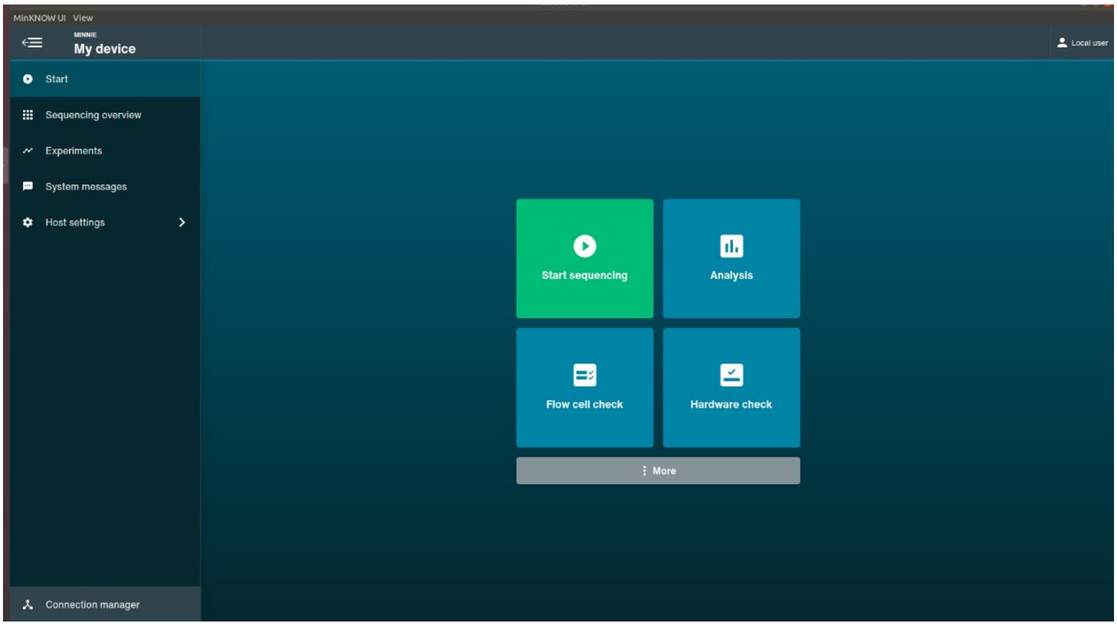

3. Enter the Flow cell ID and Flow cell type then select Start.

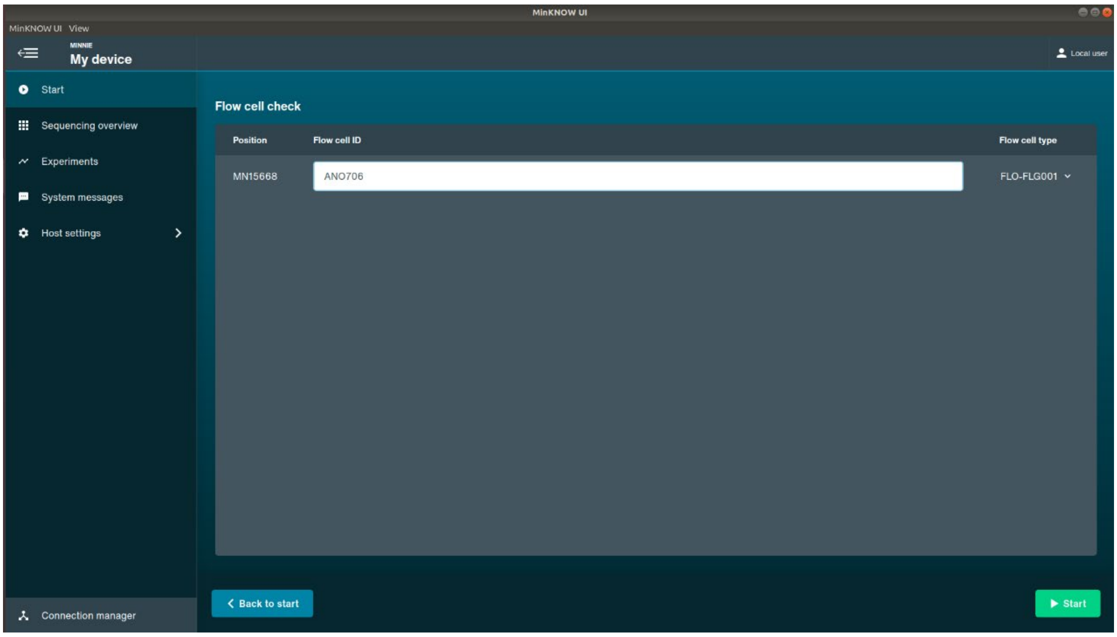

4. After ~10 minutes, a total number of pores found will be displayed under the flow cell. For a Flongle, there should be a minimum of 60 active pores (MinION – 800

pores, PromethION – 5000 pores).

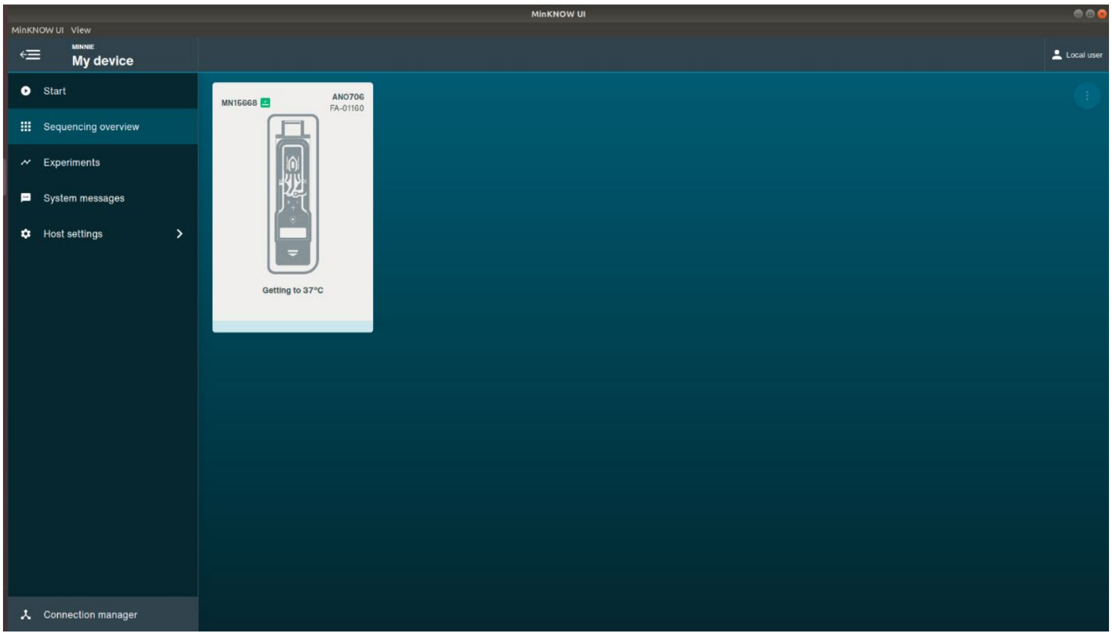

#### Setting up a run

1. Return to the 'Start' screen using the left-side menu and then select 'Start sequencing'.

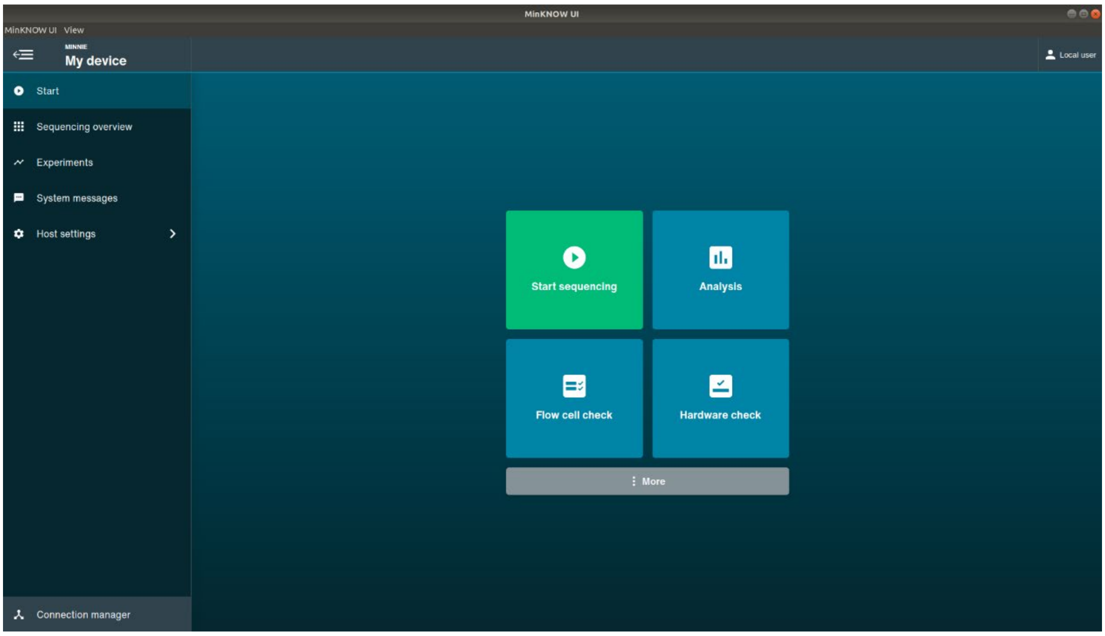

2. Select the library preparation reagents used to create your libraries. These can be filtered using the Sample type, PCR-free and Multiplexing options. Click 'Skip to final

review once you have selected your reagents.

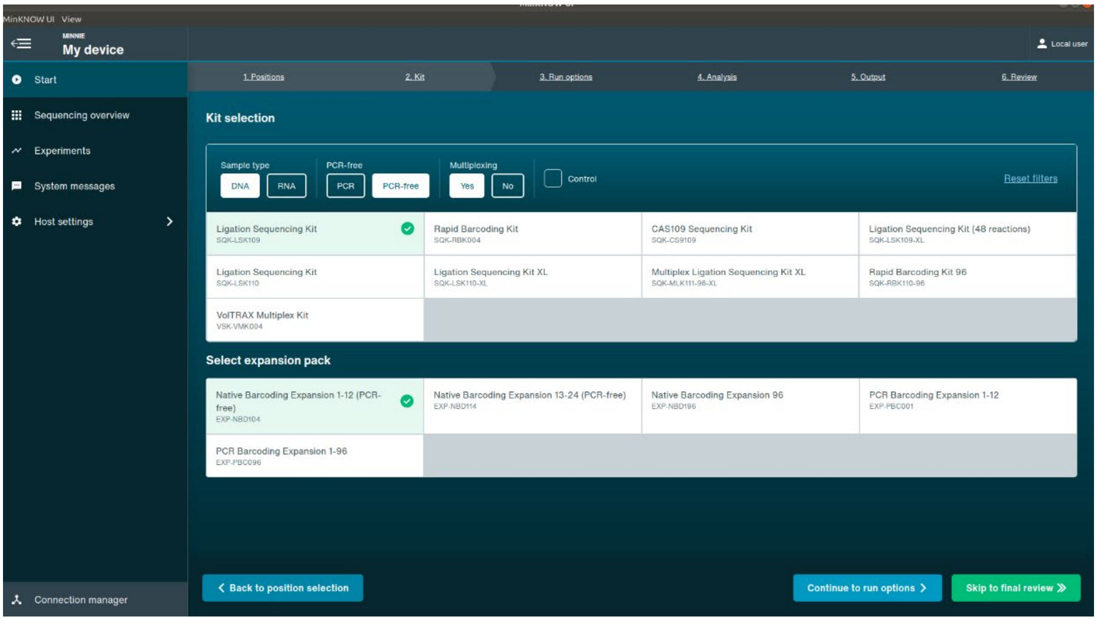

3. It is generally safe to leave the Run options, Analysis and Output settings at their default values except in specific use cases, e.g. methylation calling. Click 'Start' to being sequencing.

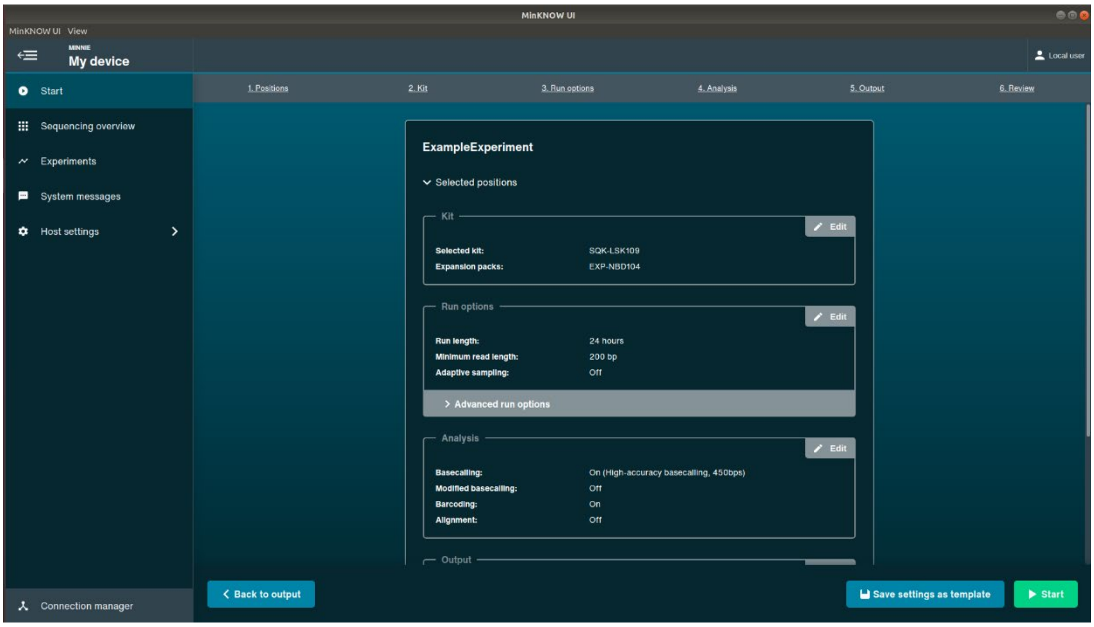

#### In-run QC Output

1. MinKNOW provides a selection of useful panels to monitor the output of your sequencing run in real-time as it progresses.
2. The channel states panel provides a visual overview of the flow cell and highlights sequencing and available pores in green. The more green, the more productive the

flow cell.

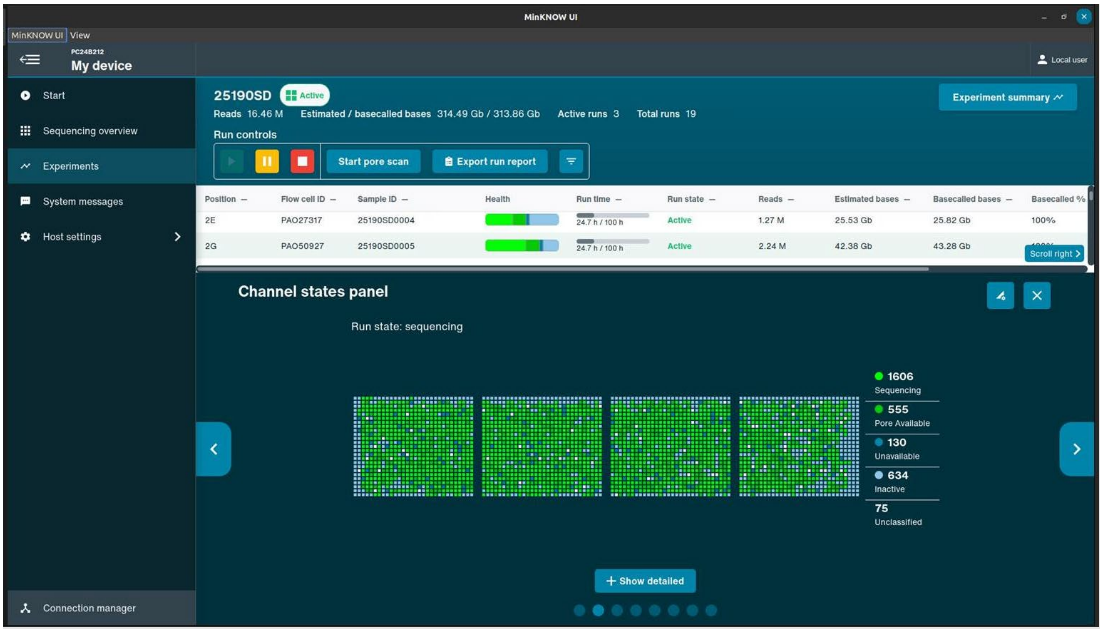

3. The pore scan results can be used to check how the health of the flow cell has progressed over time. While the health of the flow cell does gradually decrease over the course of a run, a rapid increase in saturated or unavailable pores can indicate problems with the library.

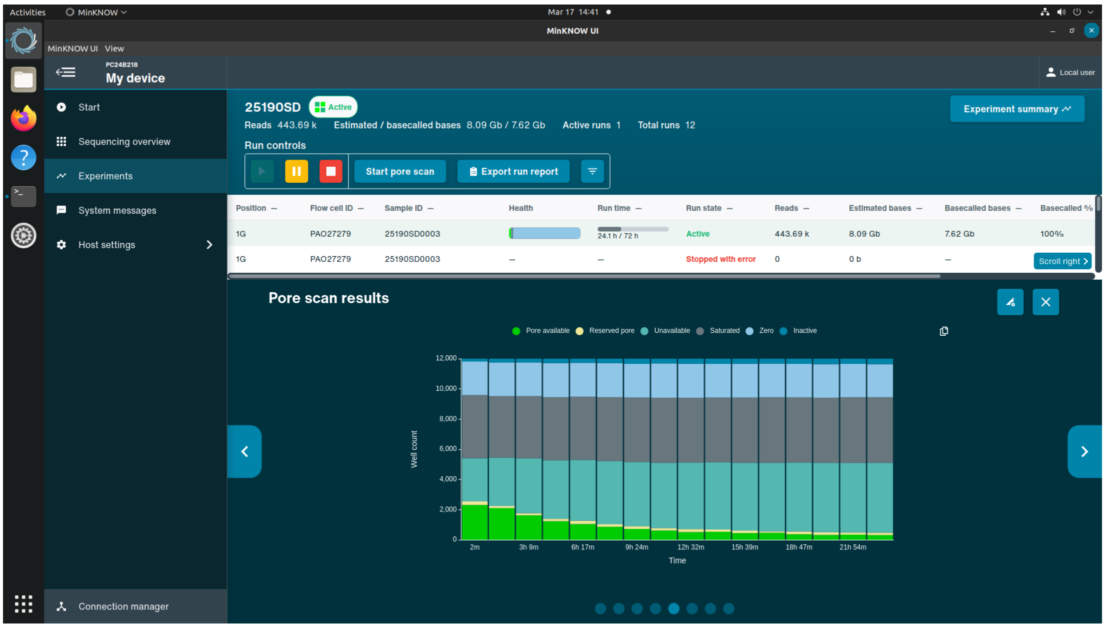

4. The read length histogram gradually fills as the run progresses. Longer fragments take longer to sequence and so may not be visible in the first ~30 minutes of a run. However, after an hour of sequencing, it provides an accurate read-out of the size distribution to expect in the final data. Ideally, the size profile should match the QC

results prior to sequencing.

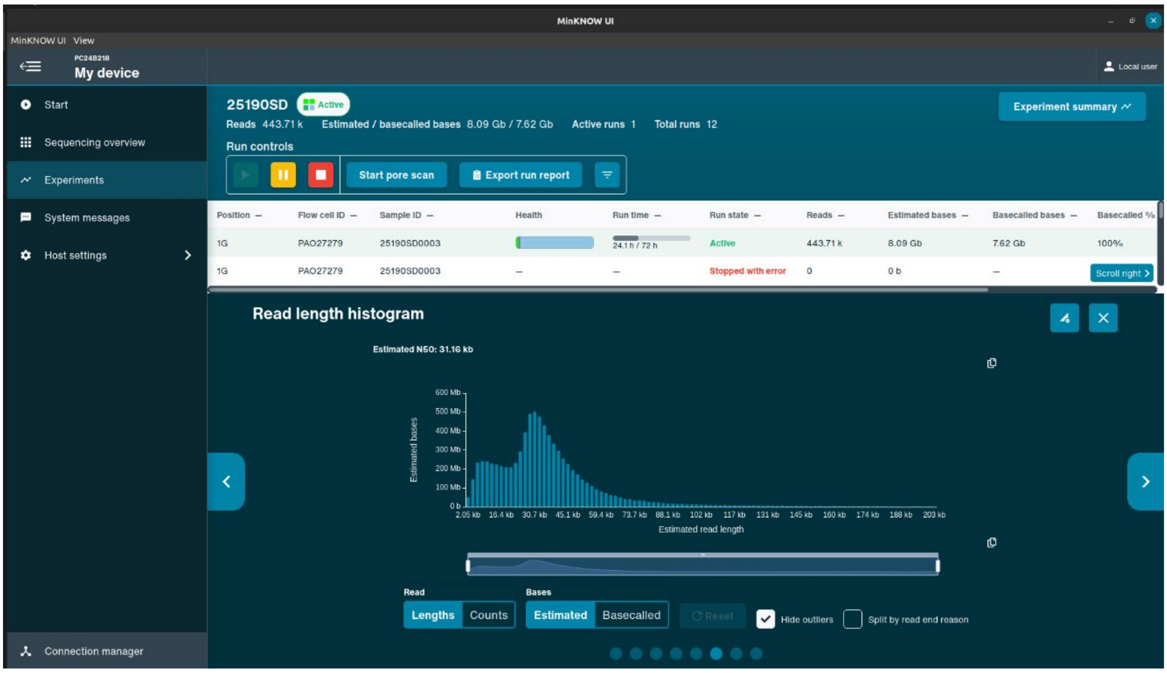

5. Finally, the Translocation speed panel is very useful for detecting when a flow cell needs to be ‘refuelled’, i.e. it requires additional ATP. The green area represents the expected translocation speed range. If the median translocation speed falls below the green area, the run should be paused and the flow cell flushed with additional buffer.

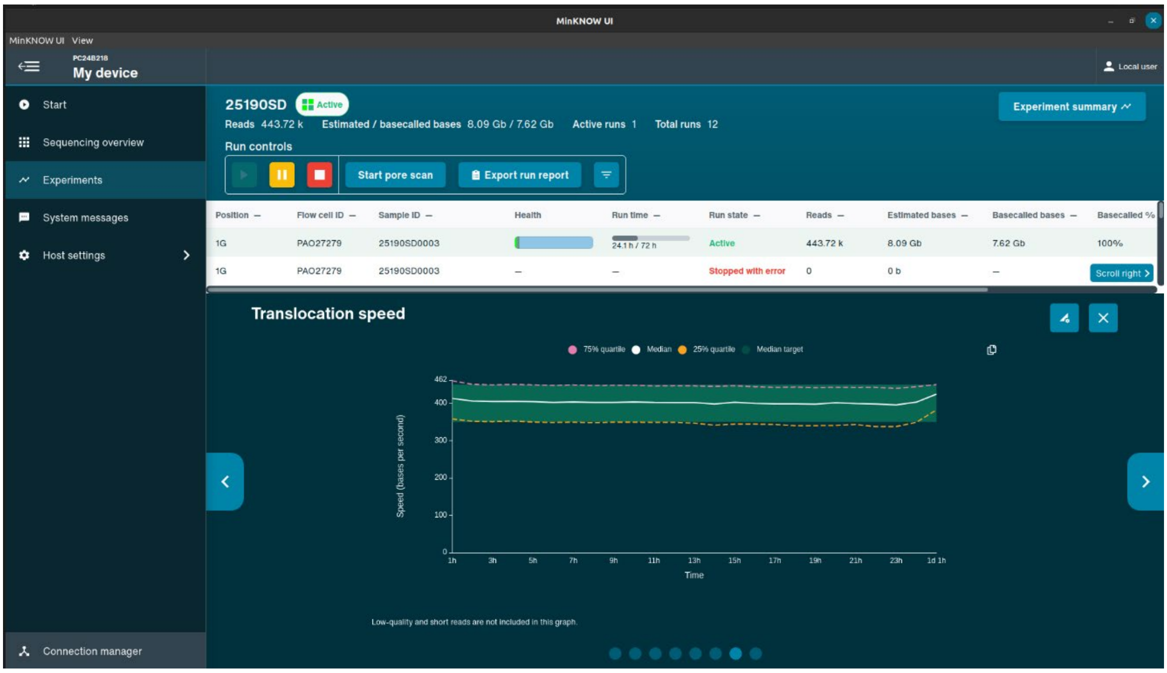
